# The size of helical pitch is important for microtubule plus end dynamic instability

**DOI:** 10.1101/2020.12.29.424770

**Authors:** Chenshu Liu, Louis Prahl, Yu He, Yan Wang, Ruijun Zhu, Yinghui Mao

**Affiliations:** Physiology Course, Marine Biological Laboratory, 7 MBL Street, Woods Hole, MA, 02543; Department of Pathology and Cell Biology, Columbia University Irving Medical Center, 630 West 168^th^ Street, New York, NY, 10032; California Institute for Quantitative Biosciences; Department of Molecular and Cell Biology, University of California Berkeley, Berkeley, CA, 94720; Department of Biomedical Engineering, University of Minnesota, Minneapolis, MN 55455; Miller Institute, University of California Berkeley, Berkeley, CA, 94720; Department of Applied Physics, Yale University, New Haven, CT 06511; Pinterest, 1099 Stewart Street, Seattle, WA 98101; Department of Physiology, University of California, San Francisco, San Francisco, CA, 94158; Department of Bioengineerinig, University of Pennsylvania, Philadelphia, PA 19104

**Author notes:** Lead Contact Chenshu Liu, **Email:**.

**Keywords:** Microtubule, Dynamic instability, Helical pitch, Seam, Computational simulation, Stochastic, Evolution

## Abstract

Microtubule (MT) dynamic instability is a conserved phenomenon underlying essential cellular functions such as cell division, cell migration and intracellular transport, and is a key target of many chemotherapeutic agents. However, it remains unclear how the organization of tubulin dimers at the nanometer scale translates into dynamic instability as an emergent property at the micrometer scale. Tubulin dimers are organized into left-handed helical MT lattice, and most present-day MTs converge at a 1.5 dimer helical pitch that causes a seam in an otherwise symmetric helix. Because presently there are no experimental methods that can precisely manipulate tubulin subunit with sub-dimer resolution, the impact of helical pitch on dynamic instability remains unknown. Here by using stochastic simulations of microtubule assembly dynamics we demonstrate that helical pitch plays essential roles in MT plus end dynamic instability. By systematically altering helical pitch size, one half-dimer at a time, we found that a helical pitch as small as one half-dimer is sufficient to inhibit short-term MT length plateaus associated with diminishing GTP-tubulin cap. Notably, MT plus end dynamics quantitatively scale with the size of helical pitch, rather than being clustered by the presence or absence of helical symmetry. Microtubules with a 1.5 dimer helical pitch exhibit growth and shrinkage phases and undergo catastrophe and rescue similar to experimentally observed microtubules. Reducing helical pitch to 0 promotes rapid disassembly, while increasing it causes microtubules to undergo persistent growth, and it is the 1.5 dimer helical pitch that yields the highest percentage of MTs that undergo alternating growth and shrinkage without being totally disassembled. Finally, although the 1.5 dimer helical pitch is conserved among most present-day MTs, we find that other parameters, such as GTP hydrolysis rate, can partially compensate for changes in helical pitch. Together our results indicate that helical pitch is a determinant of MT plus end dynamic instability and that the evolutionarily conserved 1.5 dimer helical pitch promotes dynamic instability required for microtubule-dependent cellular functions.

## INTRODUCTION

Microtubules (MTs) are dynamic, hollow polymers that are essential for cellular mechanics and signaling in diverse contexts such as chromosome segregation, cell migration and intracellular transport [1-6]. MTs are self-assembled from tubulin subunits and are polarized with a minus end and a plus end. While MT minus ends are usually anchored at the MT organizing center, the more dynamic MT plus ends interact with various subcellular structures and actively undergo growth and shrinkage[7, 8]. Growth and shrinkage of MTs occurs through net addition and removal of tubulin subunits at the MT plus end, respectively. In many eukaryotic cell types, MTs switch between periods of steady but relatively slow growth and periods of rapid shortening, a phenomenon called “dynamic instability”[9, 10]. It is key to MT functions in diverse cellular processes such as the efficient “search and capture” of kinetochores during mitosis and meiosis [11, 12], and it is tightly regulated by many microtubule-associated proteins [13-16]. It is also the focus of many microtubule-targeting agents (e.g. Taxol) that confers chemotherapeutic treatment by attenuating dynamic instability [17, 18]. Dynamic instability has been empirically described by parameters such as the rates of growth and shrinkage, and the frequency of switch from growth to shrinkage (“catastrophe” events) or of switch from shrinkage to growth (“rescue” events) [19]. Despite the discovery of dynamic instability over three decades ago, its underlying mechanisms are still not fully understood.

High resolution structural studies of MTs at different dynamic states suggests that the structural arrangement of tubulin dimers at the MT plus-end encodes dynamic information [20-23]. In many eukaryotic cells the MT lattice is formed by 13 parallel linear strands of α/β-tubulin heterodimers called protofilaments (PFs)[24, 25] (**Figure 1A**). When an α/β-tubulin heterodimer is added to the plus end of an individual PF, the β-tubulin monomer carries a guanosine triphosphate (GTP) that can be hydrolyzed and exchanged. Hydrolysis from GTP-tubulin to GDP-tubulin occurs after the α/β-tubulin heterodimer is incorporated into the lattice, and it is required for dynamic instability, since GDP-tubulin-bearing dimers are preferentially removed from plus-ends, leading to disassembly [26]. In most MTs, neighboring PFs are staggered and there is a pitch of three tubulin monomers (1.5 dimers) per turn of the helix, so that adjacent subunits in the MT lattice form a left-handed helix [27]. This is called a 13-3 (13-PFs 3-mononer start) B-lattice MT, where the majority of contacts between neighboring PFs’ subunits are homotypic (i.e. α-tubulin in contact with α-tubulin, β-tubulin in contact with β-tubulin) with one exception: those interactions between subunits of PF1 and PF13 are heterotypic, as each α-tubulin is in contact with a β-tubulin and vice versa, forming a local A-lattice at the seam between PF13 and PF1 (**Figure 1A**). This broken helical symmetry at the seam, caused by the 1.5 dimer helical pitch, remains one of the most conspicuous yet poorly understood structures of the 13-3 B-lattice MTs.

**Figure 1.**
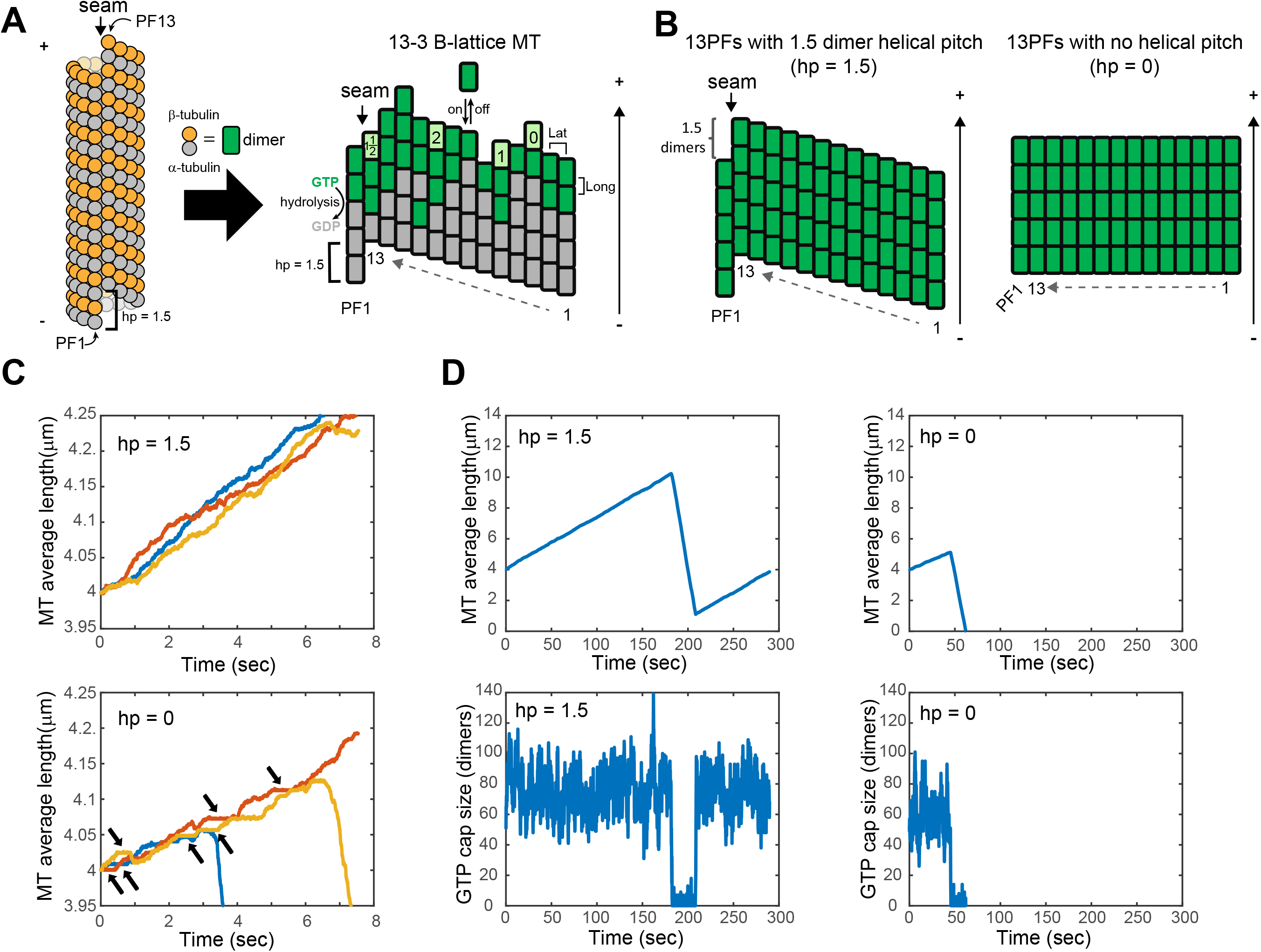
Helical pitch of the 13-3 B-lattice MT is required for ordinary MT plus end dynamics. (A) Schematics showing the organization of α-/β-tubulin dimers in a 13-3 B-lattice MT and the position of the seam between PF1 and PF13. For simplicity, in the ‘flattened’ MT lattice, each α- /β-tubulin heterodimer is represented by rectangles that is the subunit of the MT lattice. Subunits with GTP-tubulin are in green, and after hydrolysis, subunits with GDP-tubulin are in grey. In addition to a hydrolysis event, an “on/off” event is also shown at the plus end of PF7. Lateral and longitudinal bonds between subunits are also shown (labeled as “Lat” and “Long” respectively). Four subunits colored in light green are representative examples to show the number of lateral neighbors for each of them. ‘+’ and ‘-’ indicates MT plus end and minus end, respectively. ‘PF’, protofilament; “hp”, helical pitch. “hp = 1.5” stands for the 1.5 dimer (3 monomer) helical pitch. (B) Schematic showing the structural features of a regular MT with 1.5 dimer offset (helical pitch) at the seam between PF1 and PF13, in contrast to an ‘*in silico*’ engineered MT with no helical pitch or the seam. Hydrolysis is not shown here. (C) Short-term simulation reveals occasional short-term plateaus when plotting MT average length as a function of time in MTs without helical pitch, but not in regular MTs. Arrows point to such plateaus. (D) Long-term simulations show MT dynamic instability, which is altered in the absence of helical pitch. The sizes of “GTP cap” (total numbers of GTP-tubulin in the MT) were also plotted as a function of time, below each corresponding MT length kymograph.

Earlier work speculated that the helical organization of tubulin subunits in MT lattice could be more important for MT dynamics than the mechanical properties of the MTs [28]. More recently, the qualitative contribution of A-lattice content in MTs has been shown to be important for MTs’ dynamics, especially catastrophe frequency, based on EB1/Mal3 induced ectopic A-lattice formation [29]. However, the exact position and fraction of A-lattice in Mal3/EB1 treated MTs is unknown. Furthermore, the question remains: why does the prevalent B-lattice MT need a helical pitch? Under certain circumstances MTs formed *in vitro* from purified tubulin might not comply with the 13-3 B-lattice structure yet still have an overall B-lattice[30]. These MTs can have 9 ∼ 16 PFs and their structures can correlate with a helical pitch that is not 1.5 dimers[30-33]. Because these “noncanonical” MTs can only be verified with electron microscopy based on quantitative measurement of PFs number, helical symmetry and pitch size, it is difficult to perform time-lapse measurement of MT dynamics, either *in vivo* or *in vitro*, while knowing the exact underlying helical pitch sizes especially for “noncanonical” MTs[34]. It can also be experimentally challenging to manipulate the geometry of *in vitro* assembled MTs so that structural features of the lattice (e.g. PF number) other than the helical pitch itself remain unchanged[30, 33]. Finally, because most present-day microtubules have a 1.5 dimer helical pitch, the few exceptional cases with polymorphic MT lattice structures provide limited variations for systematic experimental dissection of the roles of helical pitch[35]. Therefore, it remains a mystery why present-day MTs have a helical pitch and how does the size of helical pitch affect MT dynamic instability.

Computational modeling using kinetic and thermodynamic parameters extracted from experimental measurements of MT self-assembly dynamics can recapitulate microtubule plus end dynamic instability [36-38]. Starting from a previous thermo-kinetic model of MT assembly[36], here we performed stochastic simulations to directly examine the role of helical pitch in MT plus end dynamic instability. By systematically altering helical pitch size in the model, we found that a helical pitch as small as one monomer is sufficient to inhibit short-term MT length plateaus associated with diminishing numbers of GTP-tubulin near the plus end. Moreover, by combing through a series of helical pitch sizes, we found that changes in MT plus end dynamic instability are due to quantitative scaling with pitch size, rather than qualitative dependence on axial stagger or helical symmetry. Importantly, while the 1.5 dimer helical pitch confers intermediate levels of growth/shrinkage rates and catastrophe/rescue frequencies, it yields the highest percentage of MTs that maintain dynamic instability by undergoing alternating growth and shrinkage without complete disassembly. Interestingly, although the 1.5 dimer helical pitch is conserved among most present-day MTs, other parameter sets that yield similar dynamic instability parameters may be found in some cases, indicating there could have been more than one way to make a microtubule. Our results have implications for the evolution of eukaryotic cytoskeletons with consistent structural features.

## RESULTS

### Stochastic simulations of MT plus end dynamic instability uncovers a role of helical pitch in suppressing short-term MT length plateaus

To simulate MT dynamic instability *in silico*, we started from a single state polymer model where tubulin dimers are the basic subunits of MTs. These subunits are allowed to be added to or removed from the existing MT lattice (“on” or “off” events) in a stochastic manner with a first-order kinetic rate of association (k_on_) and dissociation rate (k_off_) determined by k_on_ and the lateral and longitudinal bond free energy values (ΔG_long_/ΔG_lat_) between subunits (**Figure 1A** and see **Methods**). To simulate a single-state polymer, subunits are assumed to have identical nature (i.e. all subunits use the longitudinal ΔG_long_ and lateral ΔG_lat_ bond energies for GTP tubulin) (**Figure 1B**). These conditions are reminiscent of *in vitro* microtubule assembly experiments using the non-hydrolyzable GTP analogue GMPCPP[37, 39, 40]. Using these kinetic Monde Carlo simulations, we reproduced previous model and *in vitro* experimental measurements (**Figure S1**). Next, based on the single state model, a complete model was developed, where tubulin dimers are either GTP-tubulin or GDP-tubulin and the identity change occurs via hydrolysis (Figure A, and see **Methods**). Importantly, all subunits inside the MT lattice were considered when stochastically deciding which subunit should undergo the “off-event” to be removed from the MT (**Figure S2**, also see **Methods**).

The free energy cost for removing subunits from the MT depends on their configurations inside the MT lattice, in particular how many lateral neighbors each subunit has. Based on the position of a given subunit – on which PF and how deep it is on that PF, the exact number of lateral neighbors can be counted (**Figure 1A** and **Figure S2**). Subunits at the MT plus-end in PF2-PF12 can have zero, one or two lateral neighbors. Because of the one and half tubulin dimer offset between PF1 and PF13 at the seam (the 1.5 dimer helical pitch, hp = 1.5), we made additional adjustments when counting lateral neighbors for subunits on PF1 and PF13, to reflect the one half increment in the number of lateral neighbors (**Figure 1A, Figure S2B**). The number of lateral neighbors for subunits at the tip or embedded in the lattice were similarly accounted for. In the hp = 0 configuration, no offset exists between PF1 and PF13 and all PFs share the identical neighboring interactions, hence the MT plus-end is “blunt” with zero helical pitch (**Figure 1B, Figure S4**). Short-term simulations (8 seconds of simulated time) revealed interesting differences between ordinary MTs with 1.5 dimer helical pitch and the MTs without helical pitch. When MT average length was plotted as a function of time (kymograph, **Figure 1C**), there was an apparent shorter MT average length by the end of simulation and more frequent rapid shortenings when helical pitch was absent. In the hp = 0 condition, there were also more frequent short-term ‘plateaus’ in the kymographs where the MT average length remained approximately constant. The occurrences of these plateaus tended to appear where the standard deviation between all 13 PFs’ lengths was near-zero and average number of GTP-tubulin dimers per growth phase (“GTP-cap size”) is decreasing (**Figure S3**). Long-term simulations (5 minutes of simulated time) recapitulated the dynamic instability behavior at MT plus end (**Figure 1D**) for both the hp = 1.5 MT and the “blunt” (hp = 0) MT. The GTP-cap had a mean size of 72.5 ± 7.0 subunits in total for a regular MT during growth phase (which is on average 5.6 layers per PF, consistent with suggestions from *in vitro* experimental data that the cap must be smaller than 200 dimers [40, 41]), while it drops to zero during the rapid shortening phase. In the absence of helical pitch, catastrophes still occurred, but long-term dynamics as well as GTP-cap size were reduced (**Figure 1D**). Thus, our stochastic simulation recapitulates MT plus end dynamic instability and shows that helical pitch is required for preventing short-term MT length plateau.

### MT plus end dynamics quantitatively scales with helical pitch size and is not clustered by the presence or absence of helical symmetry

Under ordinary conditions, the regular MT’s 13-3 B-lattice is constructed with each of the neighboring PFs staggered from each other by 0.9 nm. This is encoded by the three-start configuration where the three tubulin monomers, each 4nm long, create a 12 nm pitch size (0.9 × 13 = 11.7 nm) and a helical discontinuity (i.e. a seam) between PF1 and PF13. When we created the “blunt” MT plus-end, the neighboring PFs were not staggered at all, thus no helical pitch or seam was encoded in the helically symmetric B-lattice. In other words, the making of “blunt” MTs changes at least three aspects of plus end structures, which are not mutually exclusive– a reduced helical pitch size, a lost seam (or regained helical symmetry), and a total loss of axial stagger between PFs (i.e. being blunt). Because B-lattice MTs with reduced helical pitch size or with regained helical symmetry can still be achieved without being categorically “blunt” (**Figure S4**) [27], it is of great interest to test (1) if changes in plus end dynamics are only due categorically to the absence of axial stagger (i.e. “blunt” MTs), or (2) if plus end dynamics are specifically altered depending on whether cylindrical symmetry is broken, or alternatively (3) regardless of the presence or absence of a seam, do MT plus end dynamics respond quantitatively to graded helical pitch sizes? Therefore, we sought to systematically examine the consequences of altering helical pitch size on MT plus end dynamics.

We started with zero helical pitch (hp = 0) which is the case for “blunt” MTs, and then systematically increased hp by half-dimer steps until reaching the upper limit of the size of helical pitch beyond which homotypic inter-monomer contacts between PFs would no longer be possible. Therefore, the upper limit scenario would be when inter-PF staggering is monomer length (4 nm) and the total helical pitch size for a 13-PF B-lattice MT is 13 monomer, or 6.5 dimers (i.e. subunits; see **Figure S2B** and **Methods** for more details). All the simulations were carried out with a predetermined total number of steps for approximately 5min, which is the average half-life of MTs in interphase cells as well as MTs in kinetochore-fibers of the mitotic spindle [42-46]. As the output, kymographs from independently repeated simulations show apparent differences in the slope of MT growth phase (corresponding to MT growth rates) and in catastrophe and rescue frequency, as the size of helical pitch varies (**Figure 2**). This sets up the stage for quantitative analysis of the relationship between helical pitch size and MT plus end dynamic instability.

**Figure 2.**
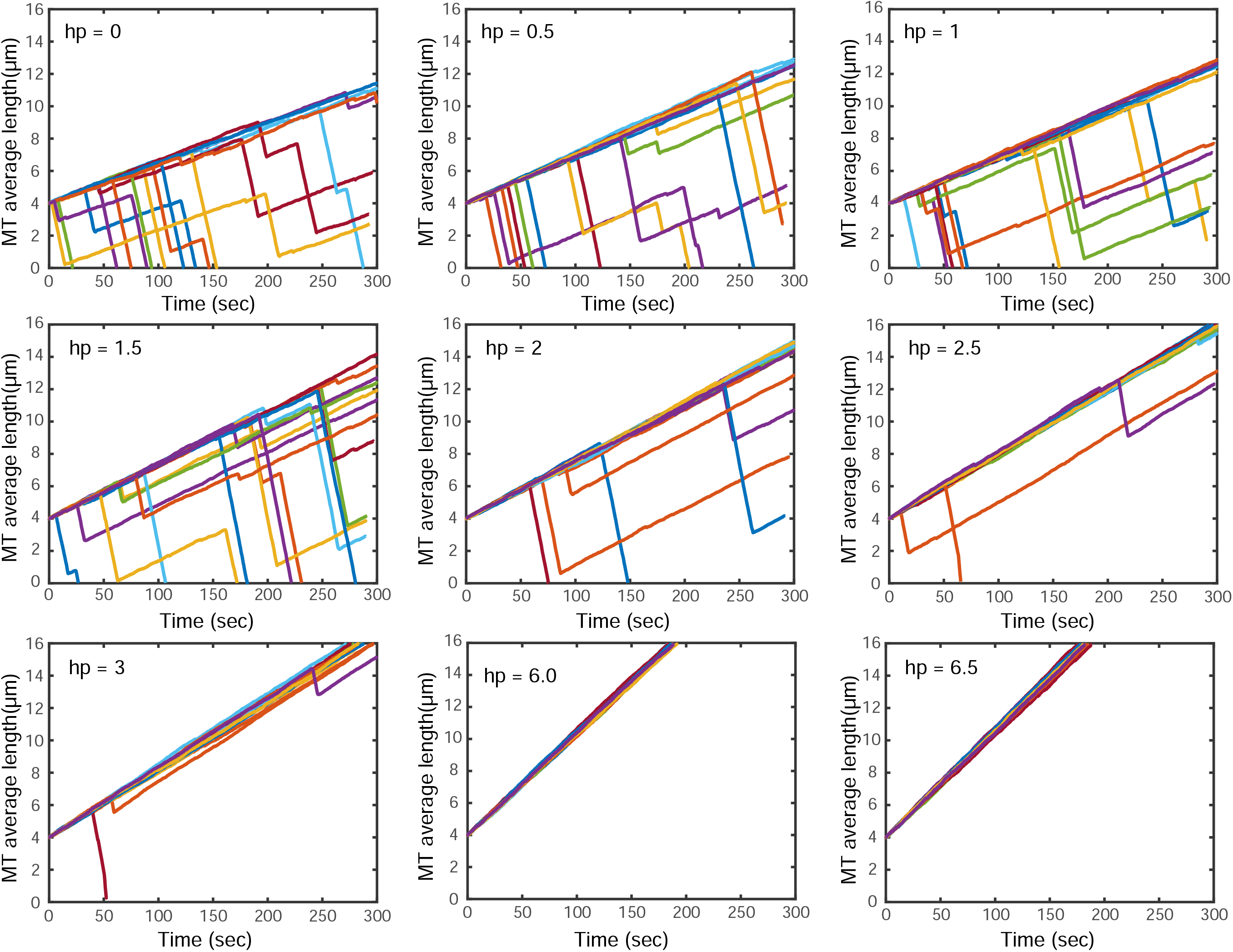
Modeling MT plus end dynamic instability with systematically altered helical pitch sizes. A series of helical pitch numbers were used to simulate MT plus end dynamics under each condition. Totally 18 simulation runs (color-coded) were shown for each individual helical pitch size, ranging from 0 to 6.5 dimers.

To investigate how MT plus end dynamic instability varies with helical pitch size, we analyzed the simulations’ raw data of MT average length and GTP cap size as functions of elapsed time, with an automated algorithm to identify catastrophe/rescue events (**Figure S5A and B**; see **Methods** for more details). Quantitative analyses revealed that the MT growth velocity progressively increases as pitch size increases, while MT shortening velocity only increases slightly as pitch size increases (**Figure 3, A&B**, also see **Supplemental Statistics Data**). Meanwhile, catastrophe frequency decreases as helical pitch size increases, while rescue frequency increases as pitch size increases (**Figure 3, C&D**). Notably, only helical pitch sized at odd number of monomers can give rise to the A-lattice seam between PF1 and PF13, or in other words, increasing pitch size at 1 monomer (half a dimer) intervals, creates MTs with alternating states of helical symmetry (i.e. presence of absence of seam). Importantly, the velocities and frequencies do not appear to be clustered based on the presence or absence of seam structures (i.e. helical symmetry) (**Figure S6**). Together with the continuous scaling between helical pitch size and dynamic instability parameters, this suggest that MT plus end dynamic instability quantitatively scales with helical pitch sizes and is not determined by either the presence or absence of the seam.

**Figure 3.**
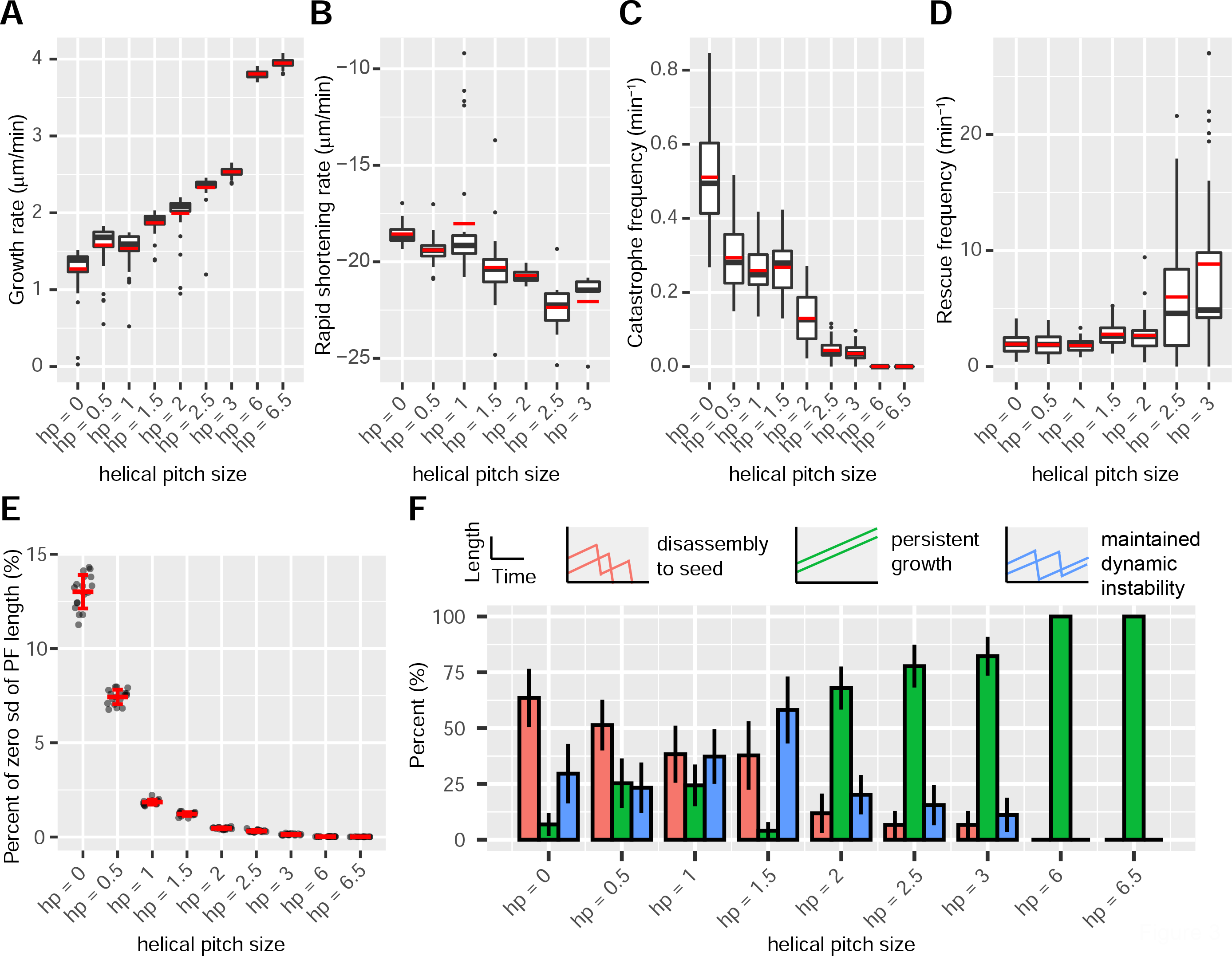
Aspects of MT plus end dynamic instability as functions of helical pitch sizes. (A and B) Velocity of growth phase (growth rate) and of rapid shortening phase (rapid shortening rate) as a function of helical pitch sizes. Boxplots (Turkey) are plotted from the velocity of individual phase (growth/rapid shortening) pooled together from totally 36 simulations per helical pitch condition, medians (black crossbars) and means (red crossbars) are shown. Krustal-Wallis and post hoc Wilcoxon pairwise tests were performed for assessing variations among and between helical pitch sizes and clustering was performed to test if the presence/absence of seam has any role in determining the velocities (see **Methods**, and the **Supplemental Statistics Data** for more details). (C and D) Boxplots (Turkey) showing catastrophe frequency and rescue frequency as a function of helical pitch sizes. Medians (black crossbars) and means (red crossbars) are shown. The lack of rescue frequencies at hp = 6 & 6.5 are because of the absence of rapid shortening phase. One-way ANOVA and post hoc pairwise t-tests were performed for assessing variations among and between helical pitch sizes and clustering was performed to test if the presence/absence of seam has any role in determining the frequencies (see **Methods**, and the **Supplemental Statistics Data** for more details). (E) Scatter plot overlaid with Mean ± SD showing the percentage of time where the standard deviation between all PFs’ lengths is zero, during the whole simulation run, as a function of helical pitch sizes. Krustal-Wallis and post hoc Wilcoxon pairwise tests were performed for assessing variations among and between helical pitch sizes and clustering was performed to test if the presence/absence of seam has any role in determining the percentage of time where the standard deviation between PF lengths is zero (see **Methods**, and the **Supplemental Statistics Data** for more details). (F) The percentage of total simulation runs that either disassemble to the seed (red), undergo persistent growth (green), or undergo maintained dynamic instability (blue), by the end of the 5 min simulation period, as a function of helical pitch sizes. Chi-squared test and post hoc pairwise comparisons of proportions were performed for assessing variations among and between helical pitch sizes. The percentage of “maintained dynamic instability” in MT with 1.5 dimer helical pitch is significantly higher than others (see **Methods**, and the **Supplemental Statistics Data** for more details). Means ± SD are plotted.

Microscopic microtubule length changes are the net result of large numbers of subunits addition or removal that culminate with fluctuations of individual PF’s length[18, 37]. Because MT average length plateaus were observed in short term simulations when MTs were blunt (**Figure 1C, Figure S3**), we decided to look more closely at the percentage of time when all the PFs are of the same length (zero standard deviation among PF length) during each simulation as helical pitch varies in size. In the absence of helical pitch, the time when plateau occurs accounts for as high as 13.0 ± 0.9 % of total time per simulation (**Figure 3E**). Interestingly, the percentage of “MT length plateau” drops exponentially as helical pitch size increases, and even a helical pitch as small as one monomer (i.e. half a dimer) is sufficient to partially inhibit short-term MT length plateaus associated with diminishing numbers of GTP-tubulin near the plus end. Notably, the percentages of “MT length plateau” does not appear to be clustered according to the presence or absence of seam structures (i.e. helical symmetry) (**Figure 3E, Figure S6C & S6C’**). This suggests two things: (1) the occurrence of “MT length plateau” is not determined by the presence or absence of helical symmetry, but scales quantitatively with helical pitch size; and (2) as the size of helical pitch increases, the 13 PFs within the whole MT tend to grow less consistently and this could potentially result in higher likelihood of persistent growth, as high degrees of length differences between PFs could create more binding sites biased toward “on-events” (**Figure 2**). Because short-term MT length plateaus are associated with diminishing GTP-tubulin cap (**Figure S3**), we next examined the size of GTP-tubulin cap at MT plus end during each growing phase (see **Methods** for details). Indeed, mean GTP-tubulin cap size per growth phase increases with helical pitch size (**Figure S8A**). Because GTP-bound tubulin subunits confer higher levels of mechanical stability in the MT lattice than GDP-bound subunits [36, 47], the positive scaling between GTP cap size and helical pitch size rather than clustering based on helical symmetry (**Figure S6F & S6F’, Figure S8A**) is consistent with MT dynamics being biased toward persistent growth at higher helical pitch sizes (**Figure 2**). Together, modeling MT plus end dynamic instability with systematically altered helical pitch sizes suggests that MT plus end structures and dynamics quantitatively scale with helical pitch sizes.

### The helical pitch with 1.5 dimer supports maintained MT dynamic instability

Given the role of helical pitch in suppressing short-term plateau and the quantitative scaling between MT plus end dynamics and helical size, we then asked: is there anything special about the 1.5 dimer helical pitch conserved among most present-day eukaryotes? The 1.5 dimer helical pitch confers intermediate levels of growth and shrinkage rates, as well as of catastrophe and rescue frequencies (**Figure 3, A to D**). Meanwhile it also results in a wide range of growth time before the onset of rapid disassembly (**Figure S8B**). This provides a possible solution for a diverse range of MT-based cellular activities that require varied MT growth time and length [48].

To further measure the diversity of MT plus end dynamics, we developed simple criteria to categorize MT kymographs into several sub-types and quantified the percentage of each among all independent simulations. The arbitrary cut-off time for all kymographs is approximately 5 min after the simulations started for reasons discussed above[42-46]. Based on the distinct behaviors of MT kymographs by the end of the 5min mark, we defined three categories of MTs that are mutually exclusive: “disassembly to seed” refers to those MTs that are completely disassembled (zero length) within the 5min time window; “persistent growth” refers to those MTs that don’t undergo discernable rapid disassembly and keep growing throughout the 5min time window; “maintained dynamic instability” refers to those MTs that undergo at least one catastrophe and one rescue and are still growing by the end of the simulations (**Figure 3F** and **Figure S5C**; see **Methods** for details). As a result, almost half of the MTs (38%) disassemble to seed at 1.5 dimer helical pitch upon the 5 min mark, consistent with 5 min being the average half-life of MTs. Importantly, the percentage of categories differ significantly according to helical pitch sizes, and there is a local peak for the percentage of maintained dynamic instability, at the expense of persistent growth, when the helical pitch is 1.5 dimers (**Figure 3F**). This result suggests that, while the 13-3 B-lattice MTs are not likely to grow persistently over the entire 5 min simulation time window, they tend to have high levels of sustained fluctuation where a MT can readily restore its growth phase after rapid shortening.

### The 1.5 dimer helical pitch is not unique and could probably have been bypassed if evolution had started differently

Microtubule dynamic instability, as characterized by parameters we have examined so far, is an emergent property manifested at the systems level. Although the 1.5 dimer helical pitch of MTs is unique under present-day parameters as it confers the highest percentage of maintained dynamic instability, it remains curious whether the well conserved 1.5 dimer helical pitch among most present-day eukaryotes results from deterministic force of natural selection, or random chance. The first step to address this is to test if the 1.5 dimer helical pitch is unique in achieving the range of emergent properties of present-day MTs. To do that, we performed simulations with 2.0 dimer helical pitch and asked the question: are there scenarios where the hp = 1.5 dynamics can be recapitulated for microtubules with a different helical pitch? Because varying the hydrolysis rate (k_H_) of GTP-tubulin can clearly affect MT dynamic instability [36, 49], and that at 1.5 dimer helical pitch, maintained dynamic instability is much less likely to happen when hydrolysis rate is less than 0.95 molecule^-1^s^-1^(**Figure S7**, top row), we therefore focused on hydrolysis rate as the second variable and examined how helical pitch (hp) and hydrolysis rate (k_H_) together affect the emergent properties of MT dynamic instability.

At 2.0 dimer helical pitch, we performed systematic simulations over a range of artificial hydrolysis rates that are equal to or higher than 0.95 molecule^-1^s^-1^ (**Figure S7**). Resultant growth and shortening rates, catastrophe and rescue frequencies were summarized in diamond graphs for comparison with the “control” scenario where the historical outcome being hp = 1.5 dimers and k_H_ = 0.95 molecule^-1^s^-1^ (**Figure 4A, Figure S8**) [36, 48, 50]. Interestingly, at least two scenarios (hp = 2.0, k_H_ = 1.10; hp = 2.0, k_H_ = 1.15) recapitulated the rates and frequencies of the “historical control”. When we quantified the percentage of each emergent subtype (“disassembly to seed”, “persistent growth”, or “maintained dynamic instability”), the scenario where the helical pitch is 2.0 dimers and the hydrolysis rate is 1.15 molecule^-1^s^-1^ has the highest percentage of “maintained dynamic instability”, which turns out to be indistinguishable from that of “historical control” (**Figure 4B**), despite differences in helical structures and average GTP-cap sizes (**Figure S8**). These results suggest that, the 13-3 B-lattice MT with 1.5 dimer helical pitch might not be unique after all, given that simple changes in thermodynamic and kinetic parameters can create structurally distinct MTs with similar emergent dynamic instability characteristics. Indeed, when varying helical pitch sizes at a given artificial hydrolysis rate constant (k_H_ = 1.15 molecule^-1^s^-1^), the percentage of maintained dynamic instability no longer peaks at 1.5 dimer helical pitch, indicating that if evolution had started from a different hydrolysis rate as “permissive” preceding change, a contingent helical pitch size different from 1.5 dimer might as well have resulted (**Figure 4C, Figure S9 to S11**). Together, our results indicate that if MT structures had evolved from a different “starting point” (e.g. a set of parameters including 2.0 dimers helical pitch thus no seam, or a hydrolysis rate that’s higher than present-day value), outcomes with different MT structure could have resulted that have quite similar salient features that characterize MT plus end dynamic instability today.

**Figure 4.**
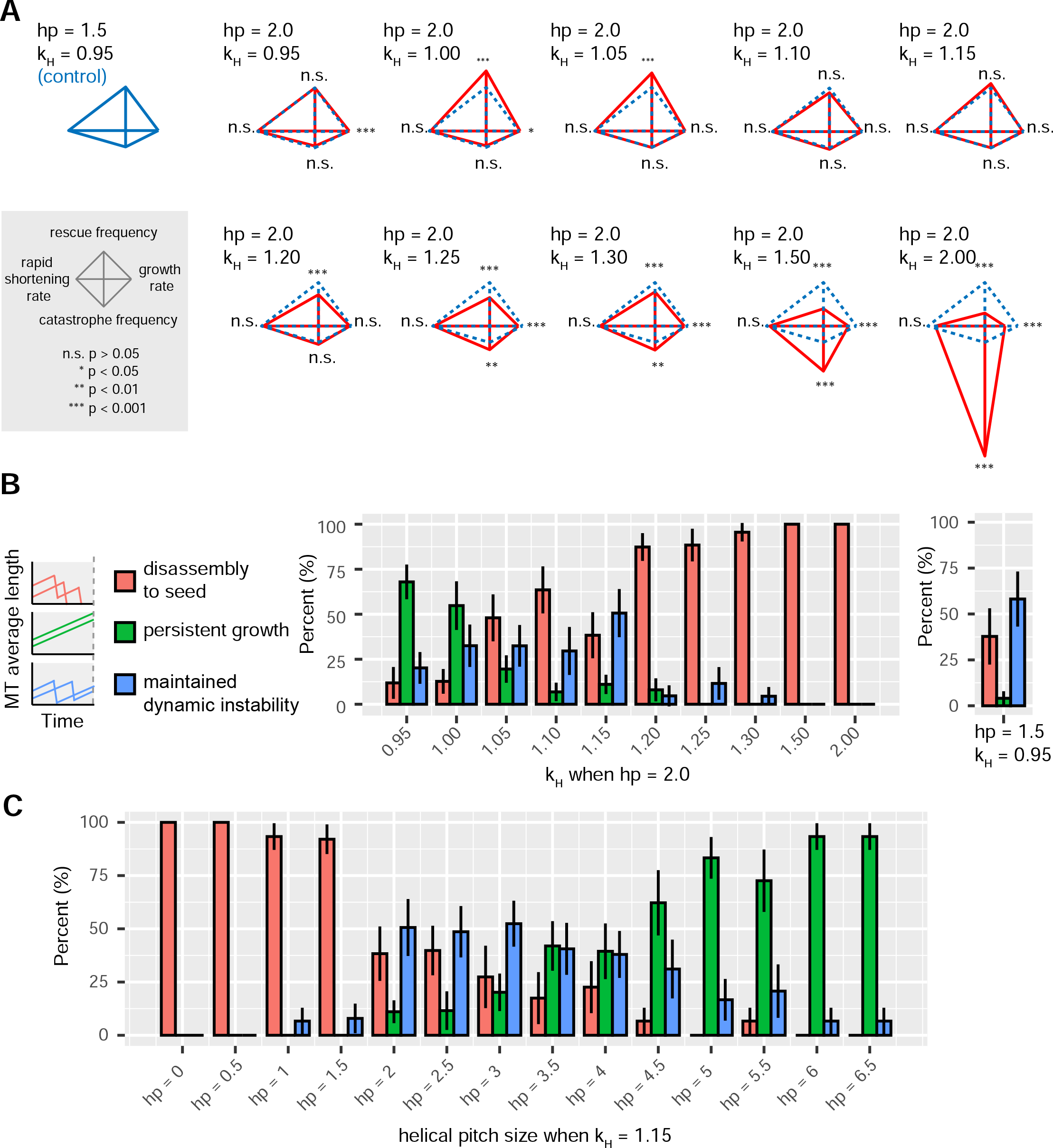
The 1.5 dimer helical pitch can be bypassed by altering hydrolysis rate. (A) Diamond graphs using two pair of individually normalized parameters describing MT plus end dynamic instability. Mean values were plotted in the diamond graphs. With the historical outcome (hp = 1.5, kH = 0.95) as control, Krustal-Wallis and post hoc Wilcoxon pairwise tests were performed for assessing variations among and between different hydrolysis rate (k_H_) conditions (at 2 dimers helical pitch) regarding rates. One-way ANOVA and post hoc pairwise t-tests were performed for assessing variations among and between different hydrolysis rate (k_H_) conditions (at 2 dimers helical pitch) regarding frequencies (see **Methods**, and the **Supplemental Statistics Data** for more details). (B) The percentage of total simulation runs that either disassemble to the seed (red), undergo persistent growth (green), or undergo maintained dynamic instability (blue), by the end of the 5 min simulation period when hydrolysis rate (k_H_) was systematically altered at a given helical pitch of 2 dimers (hp = 2). Chi-squared test and post hoc pairwise comparisons of proportions were performed for assessing variations among and between helical pitch sizes. The percentages of each category for the historical outcome (hp = 1.5, kH = 0.95) were also plotted separately (right panel) for comparison. The percentage of “maintained dynamic instability” in MT when K_h_ = 1.15 molecule^-1^ sec^-1^ at 2 dimers helical pitch is not significantly different from that in MT when k_H_ = 0.95 molecule^-1^ sec^-1^ at 1.5 dimers helical pitch. (see **Methods**, and the **Supplemental Statistics Data** for more details). Means ± SD are plotted. (C) The percentage of total simulation runs that either disassemble to the seed (red), undergo persistent growth (green), or undergo maintained dynamic instability (blue), by the end of the 5 min simulation period when helical pitch size was systematically altered and hydrolysis rate (k_H_) is given at 1.15 molecule^-1^s^-1^. Chi-squared test and post hoc pairwise comparisons of proportions were performed for assessing variations among and between helical pitch sizes. The percentage of “maintained dynamic instability” in MT with 2, 2.5, and 3 dimers helical pitch (when k_H_ = 1.15 molecule^-1^ sec^-1^) are not significantly different from that in MT with 1.5 dimer helical pitch (when k_H_ = 0.95 molecule^-1^ sec^-1^). Helical pitches sized at 3.5, 4, 4.5, 5, 5.5 dimers were included here but not in earlier simulations because a clear gap of patterns between hp=3 and hp=6 is observed here but not before (see **Methods**, and the **Supplemental Statistics Data** for more details). Means ± SD are plotted.

## DISCUSSION

Structural organizations of tubulin subunits underly the dynamic behaviors of microtubules. Despite being a possible ancient feature of subunit organization in MT-like filaments[51, 52], the helical pitch of microtubule lattice remains one of the most understudied structural features of microtubules. Particularly, it remains unclear what is the role of helical symmetry in microtubule plus end dynamic instability. This is partly because most present-day microtubules have a 1.5 dimer helical pitch, thus the low occurrence of MTs with non 1.5 dimer helical pitch provides limited variations for systematic experimental comparison [31, 33, 53-56]. Also limited is the ability to experimentally manipulate the geometry of *in vitro* assembled MTs, precisely at half-dimer resolution. Although helically symmetric B-lattice MTs without A-lattice seam can be built [27], the number of PFs are simultaneously changed besides the size of helical pitch, thus it remains challenging to probe the role of helical pitch *per se*[30]. Here by using computational simulation we demonstrate that helical pitch plays important roles in MT plus end dynamic instability. Aspects of MT plus end dynamics are quantitatively scaled with the size of helical pitch, rather than being categorically determined by the presence/absence of axial stagger or helical symmetry. Importantly, the 1.5 dimer helical pitch confers intermediate levels of growth/shrinkage rates and catastrophe/rescue frequencies and gives rise to the highest percentage of MTs that undergo alternating growth and shrinkage without being disassembled completely (“maintained dynamic instability”). Despite being uniquely supportive of “maintained dynamic instability” under present-day parameters, the requirement for 1.5 dimer helical pitch probably could have been bypassed by a modest increase (∼20%) in the GTP hydrolysis rate constant, well within the range of plausible kinetic rates. Collectively our results indicate helical pitch can be an important regulation point in the evolution of microtubule structures and functions (**Figure 5, Figure S12**).

**Figure 5.**
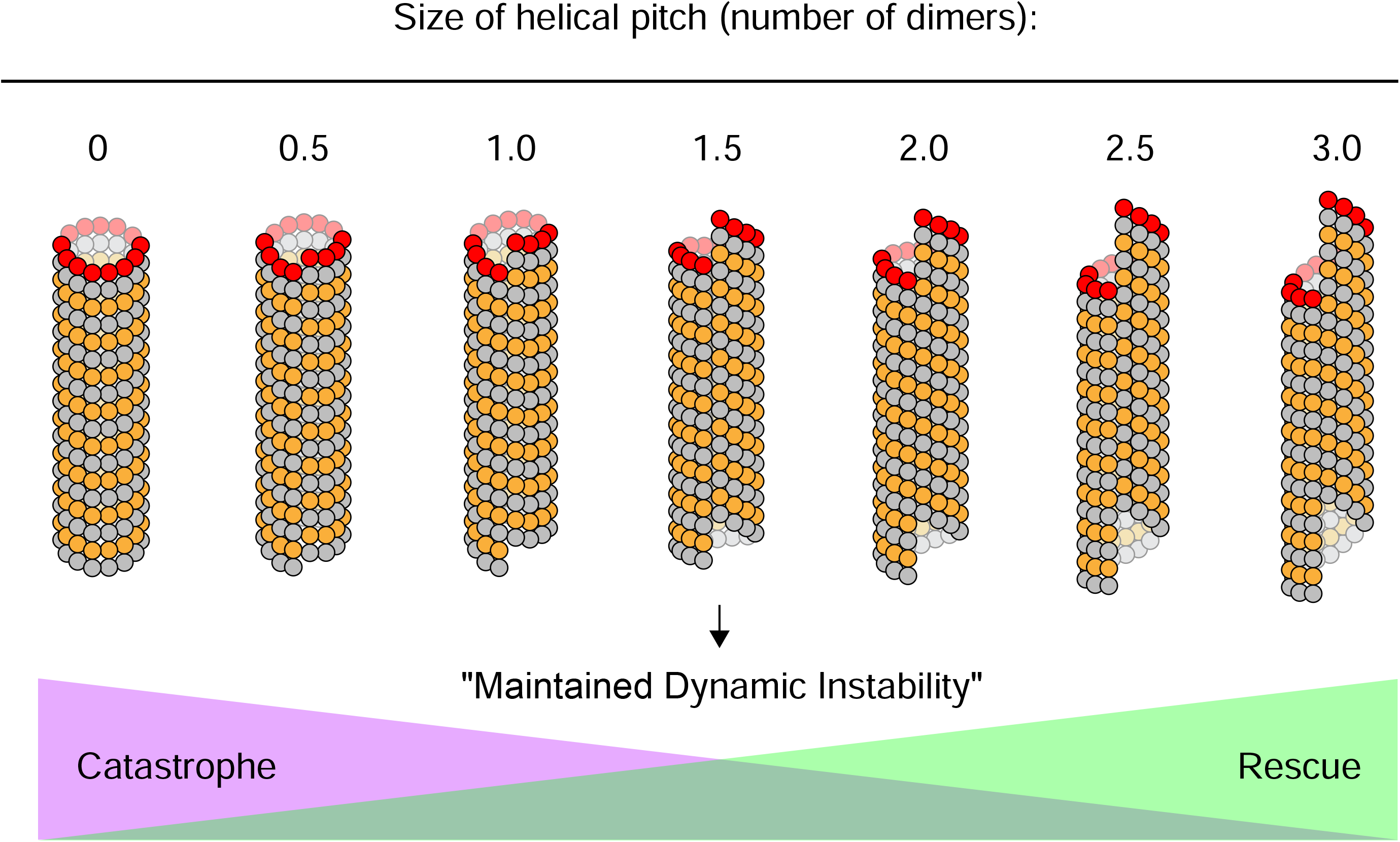
The size of helical pitch is important for microtubule plus end dynamic instability. Schematics demonstrating how MT plus end dynamics tend to change as the size of helical pitch is altered. The pitch size of 1.5 dimers by conferring intermediate levels of growth and shrinkage rates and catastrophe and rescue frequencies, yields the highest percentage of MTs that undergo “maintained dynamic instability”. The plus end of the MTs are marked by red-colored β-tubulins.

Our computational simulation enables the direct analysis of specific contribution of helical pitch on MT dynamics without being confounded by other structural features such as number of protofilaments that are changed in experiments[27]. Another major advantage of *in silico* comparison is the ability to sample through a large parameter space by systematically assigning many numerical possibilities to variables of interests. In our simulations, we were able to deviate from the historical outcome of helical pitch (hp = 1.5 dimers) and hydrolysis rate (k_H_ = 0.95 molecule^-1^s^-1^) by assigning a wide range of numerical values that is not found in present-day organisms. This method provides insights into the systems’ behavior that would be otherwise difficult to appreciate on a narrower range. For instance, we found that across a wide range of k_H_, the local maxima for the percentage of “maintained dynamic instability” correlates with a decrease in the percentage of “disassembly to seed”; whereas across a wide range of hp, the local maxima for the percentage of “maintained dynamic instability” involves a concomitant decrease in the percentage of “persistent growth”, indicating antagonistic effects of hydrolysis rate and helical pitch size in regulating dynamic instability. Such pattern might have been less obvious should only one or two pitch sizes or hydrolysis rate constants be available for experimental measurement. Also, with a wide range of helical pitch sizes, we were able to examine if readouts of dynamic instability cluster based on whether the MT has a seam or not. Our results provide direct evidence for the implication from previous experimental studies, that dynamic instability does not strictly require a seam, underscoring the power of computational simulation[57].

We limited our modeling to a single set of bond energy parameters that are used to simulate dynamics for bovine tubulin *in vitro* [37, 39]. However, as the offset initiated by helical pitch would be distributed among all 13 PFs, it will result in a stagger between each PF. Here we assume the bond energies between all neighboring tubulin subunits to be the same as the ordinary condition with a 1.5 dimer helical pitch and a hydrolysis rate at 0.95 molecule^-1^s^-1^, despite the apparently different stagger between PFs under each pitch size. Interestingly, cryo-electron microscopy has shown that the bridging densities in between adjacent PFs in A-lattice resembles that in B-lattice [58], suggesting our use of the same bond energy across a range of helical pitch sizes could be a reasonable estimate of the actual bond energies under respective scenarios. Moreover, the fact that MTs with PFs numbers other than 13 can form *in vitro* suggests there is some flexibility in the bonds between adjacent PFs[30]. Nevertheless, it is still possible through the course of evolution, lateral bond energies were adjusted to accommodate the offsets between PFs. Given the observation of non 13PF MTs which might have non-3 start helical pitch[33], experimental measurement specifically of the dynamics of non-3 start MTs would be informative for estimating bond energies. Alternatively, molecular dynamics and Brownian dynamics simulations could provide additional insights in estimating lateral bond energy given different neighboring situations caused by helical pitch [38, 59]. Meanwhile, although our simulations only focused on helical pitch and hydrolysis rate, other variables can also serve important roles in the evolution of MT structures. One such variable is the total number of PFs in a MT. MTs with divergent PF numbers were observed *in vivo* from diverse organisms[35], and MTs composed of 9 to 16 PFs were observed *in vitro* from purified tubulin [31, 33]. In some of these MTs, helical pitch sizes were observed to deviate from 1.5 dimer. Recently, “noncanonical” MTs with 11PFs found in *Caenorhabditis elegans* ventral cord neurons as well as embryos were reconstituted and measured for its dynamics *in vitro*[60], and human β-tubulin isotype was shown to control PFs number and MT stability[33]. These findings will pave the way for linking experimental measurements and systematic simulations to probe how PFs number interact with hp during the evolution of the structure and emergent properties of MTs.

In this study, our empirical definition/characterization of three types of dynamic instability is based on simulations that were run on biologically relevant time scales[42-46]. The type of simulated runs showing ‘maintained dynamic instability’ corresponds to the populations of MTs that undergo alternating growth and rapid shortening without being disassembled completely. This could be an economic and efficient way to probe the space surrounding an existing nucleation site since *de novo* nucleation could be time consuming as the kinetic barrier to nucleation makes it the rate limiting step in nucleated polymerization [5, 61]. For instance, such ‘maintained dynamic instability’ could provide a mechanism for efficient search and capture of chromosomes and timely correction of error-prone MT-kinetochore attachment, during mitosis and meiosis [62-64]. On the other hand, our results suggest that, although the 1.5 dimer helical pitch MTs is quite unique under present-day parameters as it confers the highest percentage of maintained dynamic instability, however, if evolution had started from a helical pitch size of 2.0 dimers instead of 1.5 dimers, for example, a MT with quite different structure yet performing similarly to present-day 13-3B lattice MTs could probably have resulted. The ‘alternative’ combination of parameters (e.g. hp = 2.0 dimers, k_H_ = 1.15 molecule^-1^s^-1^) we found, nevertheless, is only one scenario among all the possibilities of the vast parameter space. It remains unclear whether this ‘alternative’ scenario could be similarly accessible from a certain ancestral MT structure, compared to the present-day ‘historical outcome’ of 13-3 B-lattice MTs[65]. Future work will include (1) computationally ‘resurrecting’ the putative ancestral MT structure, and (2) building a network of possible combinatorial parameters that would potentially uncover many alternative MT structures which perform the derived function (emergent properties of dynamic instability) at least as well as the historical outcome. Recent discoveries of the bacterial BtubAB mini-microtubule that has a one-monomer helical pitch and a seam, and displays dynamic instability, illustrates what a putative “ancestral” MT might be like[51, 52]. By analyzing the number of successive changes between each functional node (a polymorphic MT with a specific set of combinatorial parameters) of the built network, one can start to examine if certain paths are uniquely accessible from the ancestral MT, and if the 1.5 dimer helical pitch is on the shortest path, or the only path should there be positive selection[65]. Meanwhile the evolutionary changes that precede the emergence of 1.5 dimer helical pitch are unknown, and answers to this question would shed light on the order of contingency, for example, whether k_H_ being 1.15molecule^-1^s^-1^ could be the permissive step required for the evolution of a non-1.5-dimer helical pitch[66, 67].

Finally, it remains unclear, on the molecular level, how the parameter space we have probed as well as that yet to be explored, could relate to the ‘sequence space’ of genes encoding tubulin isoforms/isotypes that natural selection might have acted upon. Notably, certain neurons of *C*.*elegans* have MTs with either 11 or 15 PFs, and these MTs are composed of tubulin isoforms that have specific amino acids changes compared to classic α/β-tubulin[68-70]. The wealth of mutations in these tubulin isoform genes as well as mutations that affect posttranslational modifications of these isoforms[56], could provide a useful resource for linking sequence space to the parameter space of MT dynamics. Meanwhile, it was shown that human β-tubulin isotypes can control PF number and MT dynamics[33], however, it remains unclear if specific isoforms can be used to consistently enrich MTs with non-3 start helical pitch, as measuring the dynamics of such MTs will still be required to estimate parameters and model their dynamics *in silico*. Moreover, as recent work has shown mutations that block GTP-hydrolysis result in persistent and stabilized MT growth[49], identification of more mutations/polymorphisms among tubulin genes that affect hydrolysis would also help link the sequence space with the parameter space describing MT dynamics. Despite all the possible alternative scenarios, the 1.5 dimer helical pitch of MTs is quite conserved among present-day eukaryotes. Mapping alternative evolutionary trajectories, based on connections between sequence space, parameter space and the built evolutionary networks, will help further untangle whether the 1.5 dimer helical pitch is deterministically selected, contingent upon certain prior stochastic permissive changes, or stochastically settled upon but subsequently ‘locked in’ because of other parameters accumulated after it was evolved[65, 71].

## Supporting information

Supplemental Statistics Data

## ACKNOWLEDGEMENTS

The authors thank Dr. David J. Odde and Dr. Brian Castle (University of Minnesota) for instrumental and insightful discussions on simulating microtubule dynamic instability. C.L. also thanks Dr. Tang Tang (Yale University) for stimulating discussions and critically reading the manuscript. This work was supported by the Bruce and Betty Alberts Endowed Scholarship in Physiology to C.L., a 3M Science & Technology Doctoral Fellowship through the University of Minnesota and National Science Foundation Graduate Research Fellowship 00039202 to L.S.P., and the National Institute of Health grants R01 GM89768 to Y.M.

## METHODS

### The model and simulation

The predominant 13-3 B-lattice MT with one seam is considered as the “ordinary state” for a naturally occurring MT. The construction of the model was based on previous studies [36]. Essentially, the simulation was performed with three types of events considered: GTP-tubulin being added on to the plus end of the MT lattice (“on-event”), GTP-tubulin or GDP-tubulin being removed from the polymer (“off-event”), and the GTP-tubulin subunits being hydrolyzed into GDP-tubulin that post mechanically strains to the lattice (“hydrolysis-event”). Gillespie next reaction algorithm was then deployed as the core when making stochastic decisions among the three types of events, by choosing the one with the shortest execution time. The exponentially distributed time (t) for each individual event is given by:

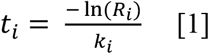

where *i* is the index of one particular event, *R* is a uncorrelated uniformly distributed random number between 0 and 1, and *k* is the (pseudo) first-order rate constant of that event (s^-1^), which was computed as following, depending on the specific type of event [36, 72, 73].

(a) For an “on-event”, the pseudo-first order rate constant (s^-1^) is given by the k_on_ which is a constant (4 µM^-1^s^-1^PF^-1^) multiplied by the concentration of GTP-tubulin and the total number of PFs.

(b)For an “off-event”, the first order rate constant k_off_ (s^-1^) is determined by the energy of the subunit under consideration. Based on the standard Gibbs free energy change:

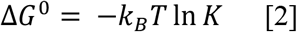

where k_B_ is Bolzmann’s constant, T is temperature (Kelvin), and K is the equilibrium constant (M^-1^) given by:

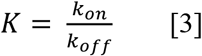

the first order rate constant of an “off-event” would be directly governed by the energy instead of the concentration of tubulin subunits:

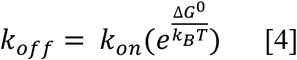

To obtain the standard Gibbs free energy change (ΔG^0^) of a reversible polymerization reaction, total bond energies of each single subunit under consideration is summed up [36]. Every subunit (either at the plus end tip or embedded in the lattice of MTs) has one longitudinal bond with the subunit right below it toward the minus end of the MT, and the longitudinal bond energy (including immobilizing energy) is 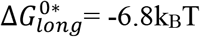. Each subunit also has a specific number of lateral bonds with its lateral neighbors (if any) at any given moment, and the lateral bond energy for a GTP-tubulin subunit under consideration is 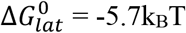. If the current subunit under consideration is a GDP-tubulin, its lateral bond energy is adjusted by adding an extra kinking energy that emulates the mechanical instability caused by the GDP-tubulin. Based on how the kinking energy is derived [36], the kinking energy Δ G_kink_ = 2.5k_B_T is added per lateral bond for a subunit under consideration, not per subunit. It is important that an “off-event” must be allowed anywhere inside the MT (either at the plus end tip or embedded in the lattice), and Δ G_kink_ is added per lateral bond, so that rapid shortening can occur at a velocity comparable to experimental measurements (data not shown). Meanwhile, due to the mechanical instability introduced to the model by GDP-tubulin, additional mechanical rules were followed to “tag” incoming tip GTP-tubulin subunit as if it is a GDP-tubulin if at least two out of its four adjacent neighbors (at the same depth or more toward the plus end) are GDP-tubulin [36].

The total standard Gibbs free energy change (ΔG^0^) is then computed by:

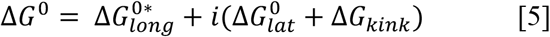

Where i is the number of lateral bonds for the subunit under consideration. Δ G_kink_ is zero if the subunit under consideration is GTP-tubulin, and takes the value of 2.5k_B_T if the subunit under consideration is a bona fide GDP-tubulin or “tagged” GDP-tubulin.

Finally, the number of lateral bonds is counted for every single subunit in the MT, either at the plus end tip or buried in the lattice (for both the single state model and the complete model, when stochastically deciding which subunit should undergo the “off-event” to dissociate from MT). Notably, because removing a subunit under consideration would also require that all subunits’ lateral bonds be broken between this subunit and the plus end tip along that PF, therefore cumulative Δ *GG*^0^ needs to be obtained as the total free energy cost used for computing the execution time for an “off-event”. If the subunit under consideration is deeply embedded inside the MT lattice, the free energy cost for removing it would be enormous (**Figure S2A**), therefore reducing the probability for deeply embedded subunits being taken off spontaneously.

Because of the offset between PF1 and PF13 at the seam (depending on how big the helical pitch is), additional adjustments were made when counting lateral neighbors for subunits of PF1 and PF13, namely the amount of increment in the number of lateral neighbors. Please refer to **Figure S2B** for more details and graphical illustration.

(c) For a “hydrolysis-event”, the pseudo-first order rate constant (s^-1^) is determined by the k_H_ which is a constant (0.95 molecule^-1^s^-1^) multiplied by the total number of GTP-tubulin that are available to be hydrolyzed. And hydrolysis is allowed for any GTP-tubulin in a MT.

After choosing the shortest execution time for deciding what to happen at one time step, the simulation procedure is implemented by updating the MT structure. If an “on-event” is chosen, secondary random numbers are generated to decide which particular PF to add the incoming GTP-tubulin to. If a “hydrolysis-event” is chosen, secondary random numbers are generated to decide which particular GTP-tubulin should be changed into the status of GDP-tubulin that will subsequently influence the mechanical status in the next round of simulation time step. If an “off-event” is chosen, the subunit under consideration together with all the subunits above it (toward the plus end) on that PF is peeled off, by breaking one longitudinal bond and all the lateral bonds for subunits to be removed.

Finally, when simulating plus end dynamics for MTs with varied helical pitch sizes, an upper limit of the size of helical pitch was noticed beyond which homotypic inter-monomer contacts would be lost. Because of the relation between pitch size and the amount of staggering between adjacent PFs:

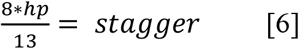

where 8 is the length of each tubulin dimer (8nm), hp is the size of helical pitch (number of dimers), 13 is the total number of PFs per MT, and the stagger is the offset between adjacent PFs. In the case of ordinary MTs, the stagger is (8 * 1.5)/13 ≈ 0.9 nm. When homotypic contacts between subunit is just about to be lost, stagger would be 4nm, therefore the upper limit for the size of helical pitch would be 6.5 dimers.

All simulations were carried out at the same free GTP-tubulin concentration (10µM). Helical pitches sized at 3.5, 4, 4.5, 5, 5.5 dimers were included only when k_H_ = 1.15 molecule^-1^sec^-1^ but not under other circumstances because a clear gap of patterns between hp=3 and hp=6 was only observed when k_H_ = 1.15 molecule^-1^sec^-1^.

### Automated post-simulation data processing

The direct output of our simulation includes documented MT lengths over time (based on which kymograph was plotted), GTP-cap size (estimated as the total number of GTP-tubulin subunits in the microtubule) over time, standard deviation among all PFs’ length at each time step, etc. In order to compute rates and frequencies, unbiased marking of catastrophe and rescue events are needed. Essentially, marking is performed using the GTP-cap dataset. Catastrophe event is defined when the cap size first has a value of zero and rescue event is defined as the last time point when the cap size is at the beginning of being non-zero for five seconds consecutively. In particular, for GTP cap size, if:

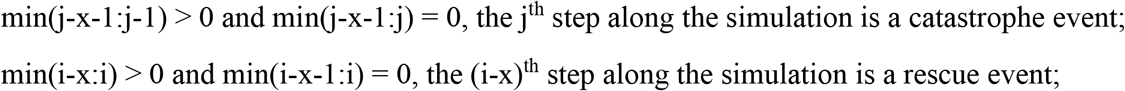

where x is the intervals of steps corresponding to 5 sec in time, as demonstrated by the schematics in **Figure S5B**.

Rates (velocities) and frequencies were computed based on the marked events. In particular, frequencies are defined by the total number of events divided by the total time accumulated in each phase leading to the events: For catastrophe frequency, total number of catastrophe events was divided by the total time spent on growth phases. For rescue frequency, total number of rescue events was divided by the total time spent on rapid shortening phases.

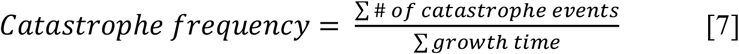

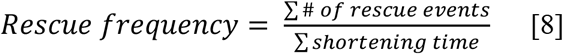

Therefore with multiple simulated trajectories (or simulated trajectories for longer time) the total time and the total number of events over all simulations can be added up. For a particular simulated trajectory, even if a catastrophe or rescue event does not occur (number of event is zero), the time spent in the growth or shorting phase still contributes to the total time. Here 18 simulation trajectories were pooled to compute the frequencies as one datapoint, and multiple repeats were used for estimating sample means.

For categorizing kymographs into three different subtypes, automatically marked kymographs were evaluated at the end of simulation. If a simulation trajectory has one or more catastrophe events while the mean MT length is zero (or total simulation ends before 5 min), the trajectory is categorized as “disassembly to seed”. If a simulation trajectory has no catastrophe event over the entire 5 min simulation time, it is categorized as “persistent growth”. If a simulation trajectory has at least one catastrophe event and the mean MT length is non zero and still growing, it is categorized as “maintained dynamic instability” (**Figure S5C**). The situation where a simulation trajectory has at least one catastrophe event and the mean MT length is non zero and is undergoing shortening is not included in our analysis due to the uncertainty of fate in the vicinity of the boundary of 5 min mark (i.e. could be either “disassembly to seed” or “maintained dynamic instability”).

### Statistics and plotting

All statistical tests (Chi-squared test and post hoc pairwise comparisons of proportions, Krustal-Wallis and post hoc Wilcoxon pairwise tests, One-way ANOVA and post hoc pairwise t-test) are noted in respective figure legends and were performed in R (RStudio, Version 1.2.5033), Graphpad Prism [16, 74, 75], or MATLAB (Mathworks, R2018a) [76]. Details of all statistics can be found in **Supplemental Statistics Data**.

To examine if aspects of MT plus end dynamic instability are categorically determined by the presence of absence of seam (i.e. helical symmetry), hierarchical and K-medoids clustering were performed in R using custom code with libraries including tidyverse, cluster, factoextra, RColorBrewer, dendextend and gplots. Unsupervised clustering was performed using both hierarchical clustering and k-medoids clustering. For hierarchical clustering, complete linkage method was used[77-80] as it generates well-balanced and compact clusters[81], and correlation matrix heatmaps and dendrograms were combined to show the resultant clusters. Average linkage method was also used and led to the same conclusions. For K-medoids clustering, partitioning around medoids (PAM) algorithm was used [82-84], and correlation matrix heatmaps were shown by indexing on the output of the ordered cluster assignments (k = 2 was predetermined to test if helically symmetric MTs and helically asymmetric MTs will form two clusters).

All plots were made in R, MATLAB or Graphpad Prism. In particular, diamond graphs were generated in MATLAB using custom code with the glyphplot function. Boxplots, barplots and scatterplots were created in R using the library ggplot2.

## Abbreviations

MT: microtubule
PF: protofilament
hp: helical pitch.

---

**Figure S1.**
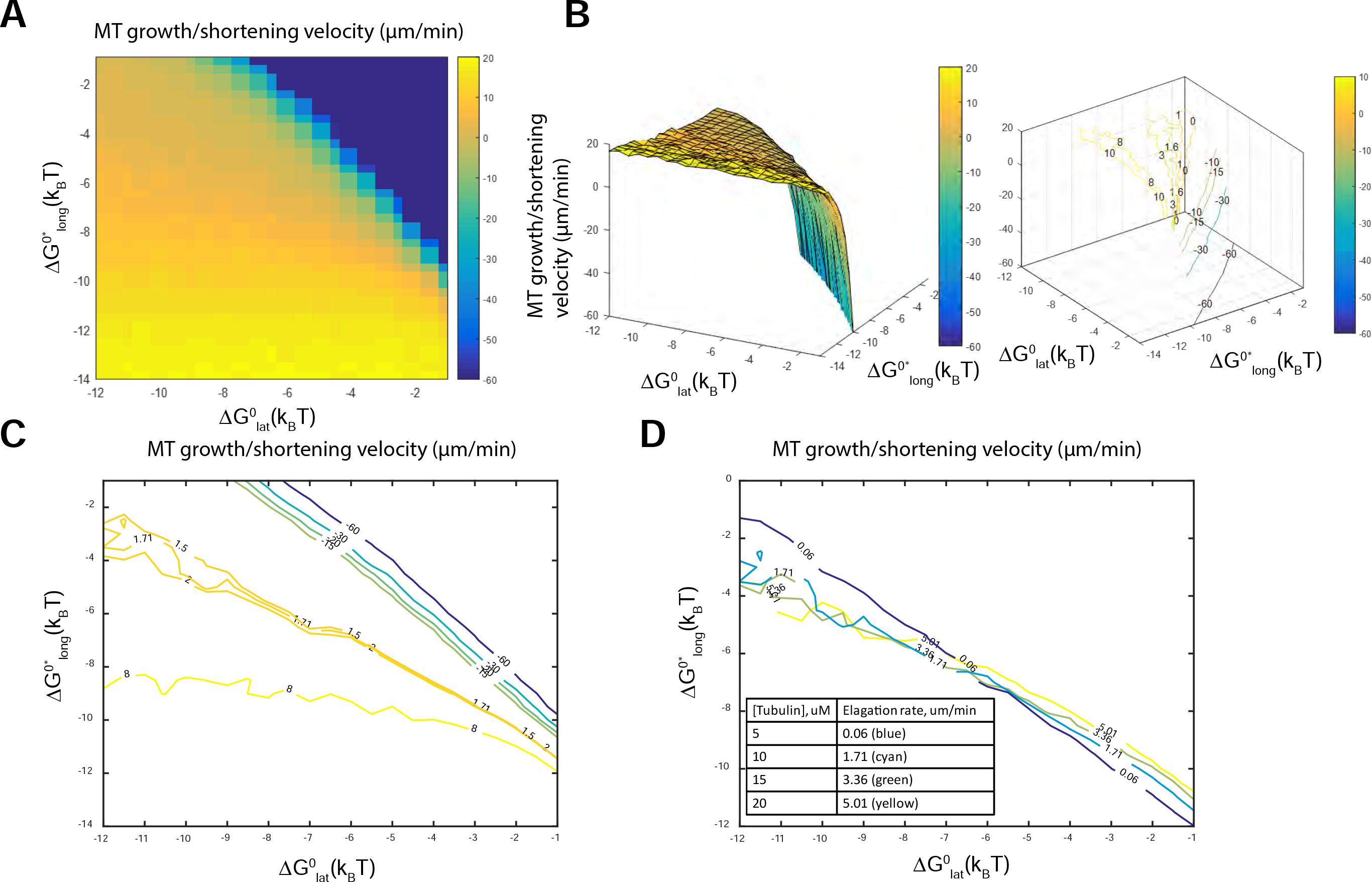
The single state model, without hydrolysis and mechanical rules being invoked, recapitulates energetic landscape in estimating lateral and longitudinal bond energies of tubulin-tubulin interactions. (A and B) XY-view and XYZ-view of MT growth/shortening velocities mapped over the entire bond energy space. Velocities are color-coded in both panels. Using these kinetic Monte Carlo simulations, we systematically investigated how changes in lateral and longitudinal bond energies affect MT growth/shortening velocity. A sharp transition from growth to rapid shortening was observed with relatively high bond energies. (C and D) Typical MT growth/shortening velocities were plotted as contours (C), and experimentally measured MT growth velocities at different tubulin concentrations were plotted (D). Notably, when plotted as contours at typical and experimentally measured MT growth/shortening velocities, our model reproduces previous model and *in vitro* experimental measurements of MT dynamic instability (VanBuren et al., PNAS, 2002 [36]).

**Figure S2.**
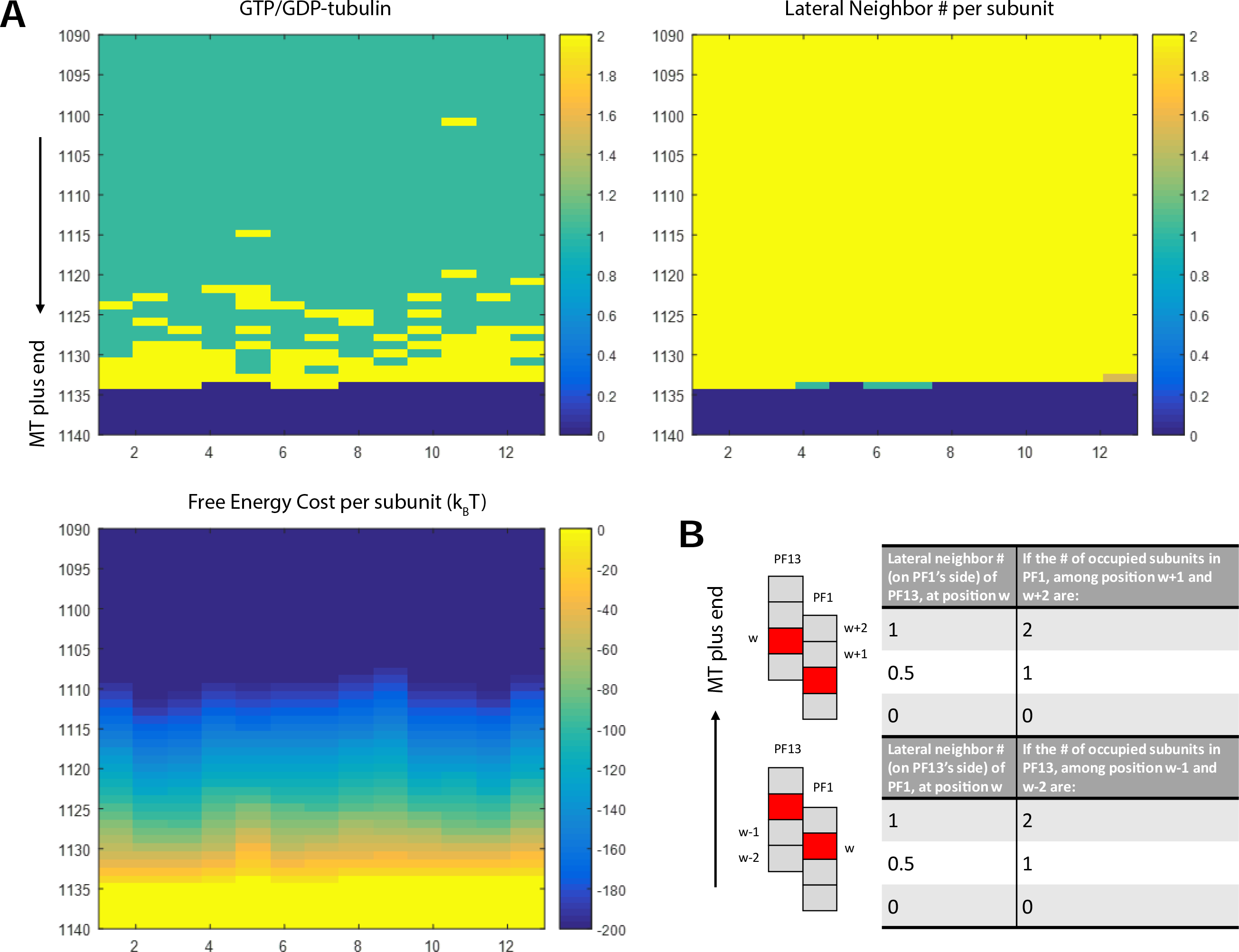
Matrices structures in the complete model, after hydrolysis and mechanical rules were invoked. (A) A snapshot during growth phase is shown, visualized by three matrices. In the GTP/GDP-tubulin matrix (upper left), each GTP-tubulin is depicted in yellow (value of 2 in the colormap scale), while each GDP-tubulin in green (value of 1 in the colormap scale), and all empty space beyond the MT plus end in dark blue (value of 0 in the colormap scale). In the lateral neighbor # matrix (upper right), the number of lateral neighbors for each subunit (tubulin dimer) is color coded (possible values include 0, 0.5, 1, 1.5, 2). Finally, the free energy cost for removing each subunit at the tip or from inside the lattice is computed (lower left). For all panels, X-axes correspond to protofilaments PF1∼PF13, Y-axes are the row numbers in the actual matrices corresponding to the position of each tubulin dimer in the lattice. MT plus ends are pointing down in all three panels. (B) Graphic illustration of how the numbers of lateral neighbors are counted at the seam. For simplicity, only the case where helical pitch has 1.5 dimers is shown, and the number of lateral neighbors under other conditions with different helical pitch sizes can be counted similarly. Each box represents one tubulin dimer, red-colored boxes are the corresponding tubulin dimer at the same depth in the MT lattice before the offset occurs. MT plus ends are pointing up.

**Figure S3.**
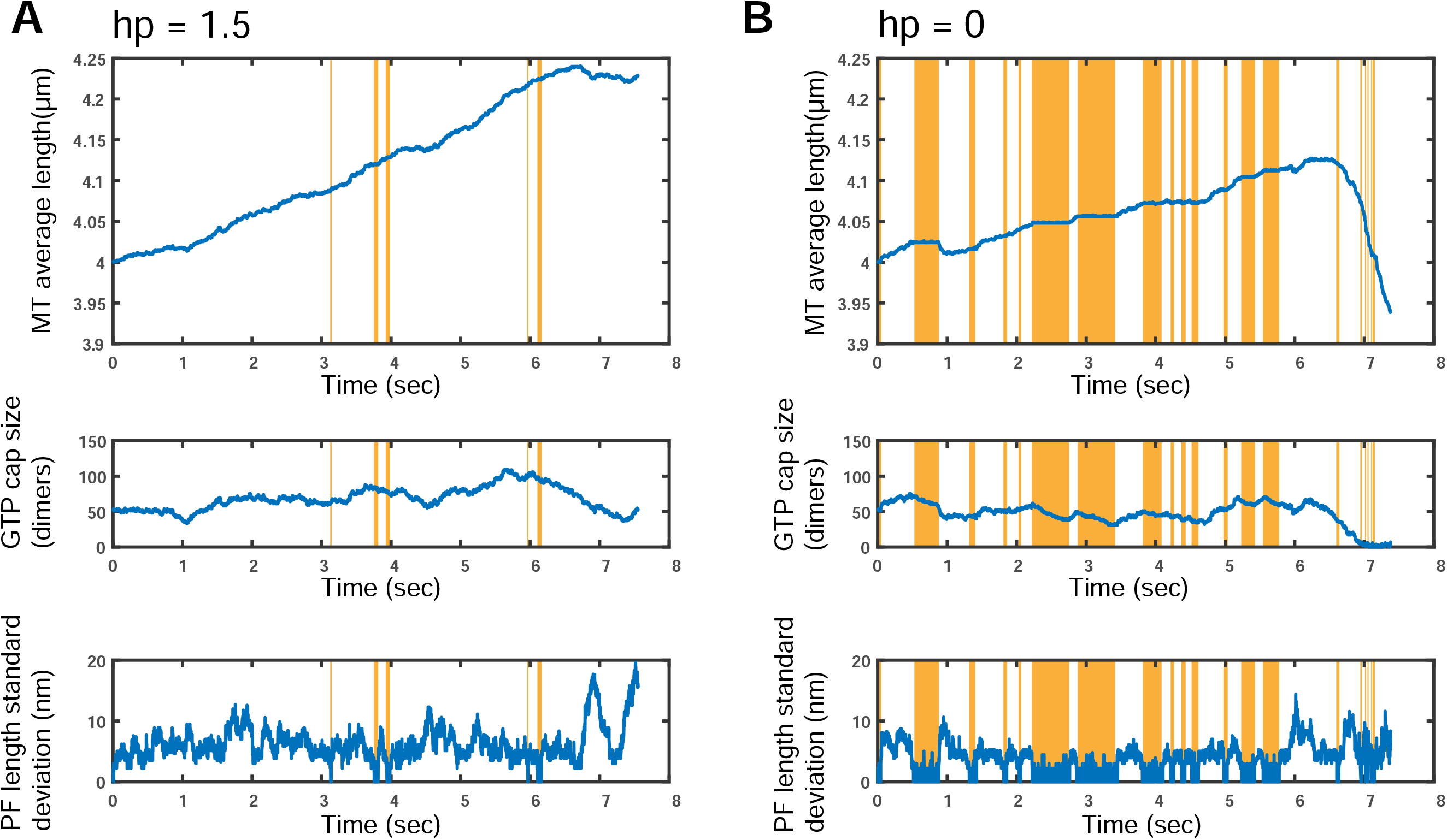
Structural details underlying short term simulations with or without helical pitch. GTP-tubulin cap sizes and protofilament (PF) length standard deviation were shown for each condition. (A) In the case of 13-3 B-lattice MT structure (hp = 1.5), GTP cap size is generally about 50 or higher, and the PF standard deviation is seldom zero. (B) In the case of MT structure without helical pitch (hp = 0), GTP cap size is on average smaller than that in (A) and the PF standard deviation is more often to be zero. For both (A) and (B), orange colored vertical bars highlight the segments of simulation time when all PFs are of the same length. During these segments of time, MT average length stays at plateau, while GTP cap size is decreasing (although when the GTP cap size is decreasing, all the PFs are not necessarily of the same length).

**Figure S4.**
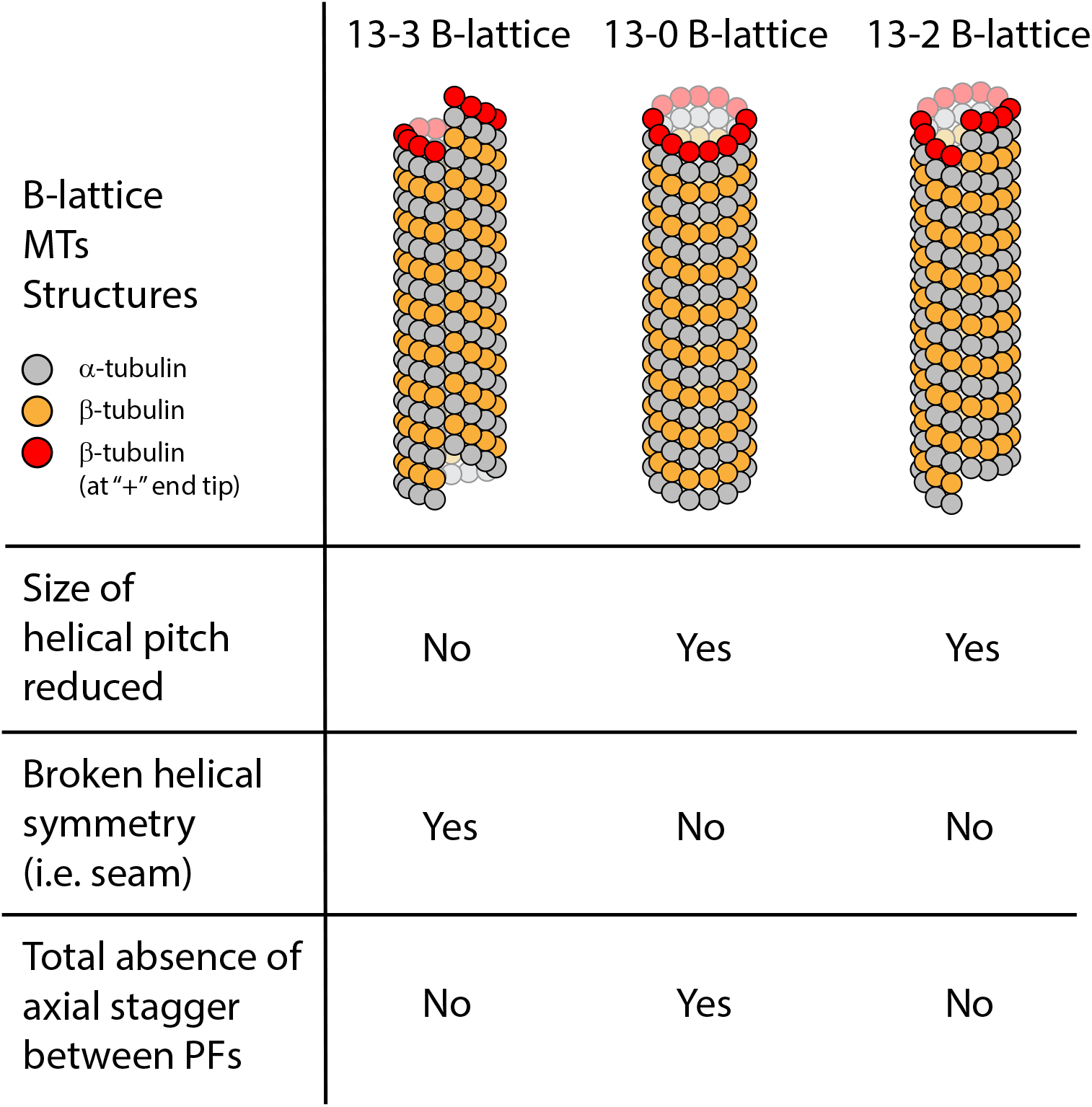
Comparing three aspects of MT plus end structures, which are not mutually exclusive. Given the three representative scenarios (i.e. 13-3 B-lattice, 13-0 B-lattice and 13-2 B-lattice), it is obvious that although a blunt MT (13-0 B-lattice) has reduced helical pitch, a MT with reduced helical pitch does not have to be blunt (e.g. 13-2 B-lattice). Likewise, a blunt MT doesn’t have a seam, yet a MT without a seam (i.e. with helical symmetry) does not have to be blunt (e.g. 13-2 B-lattice). The plus end of the MTs are marked by red-colored β-tubulins.

**Figure S5.**
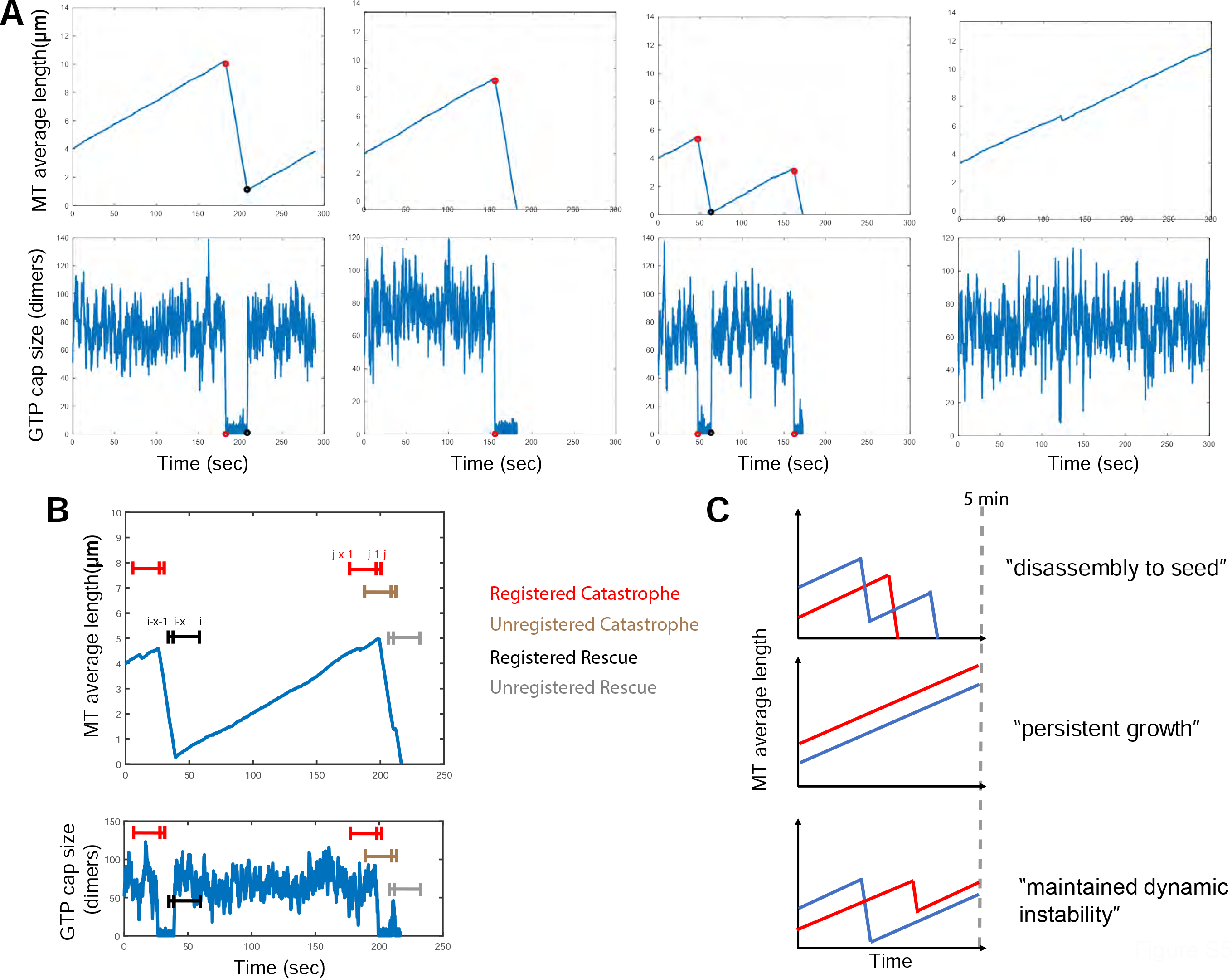
Demonstration of how catastrophe and rescue events were automatically marked for quantitation and how the three types of MT dynamics are defined. (A) Schematic demonstrations showing representative scenarios where catastrophe and rescue events are automatically marked in accordance to the “5 sec rule” (see **Methods**). Representative examples where catastrophe (red circle) and rescue (black circle) events were marked based on the size of GTP-tubulin cap. The ‘GTP cap’ sizes were plotted as functions of time, below each corresponding MT length kymograph. (B) Schematic demonstrations showing how the “5 sec rule” is applied in registering catastrophe and rescue events. Marking is performed based on GTP cap size data. Red and Black, registered events; Brown and grey, unregistered events. See **Methods** for more details. (C) Schematics demonstrating how the three categories of trajectories that have happened on or before the 5 min mark of simulation, are defined – namely “disassembly to seed”, “persistent growth”, and “maintained dynamic instability”. Trajectories not included here are dropped out when quantifying percentage. See **Methods** for more details.

**Figure S6.**
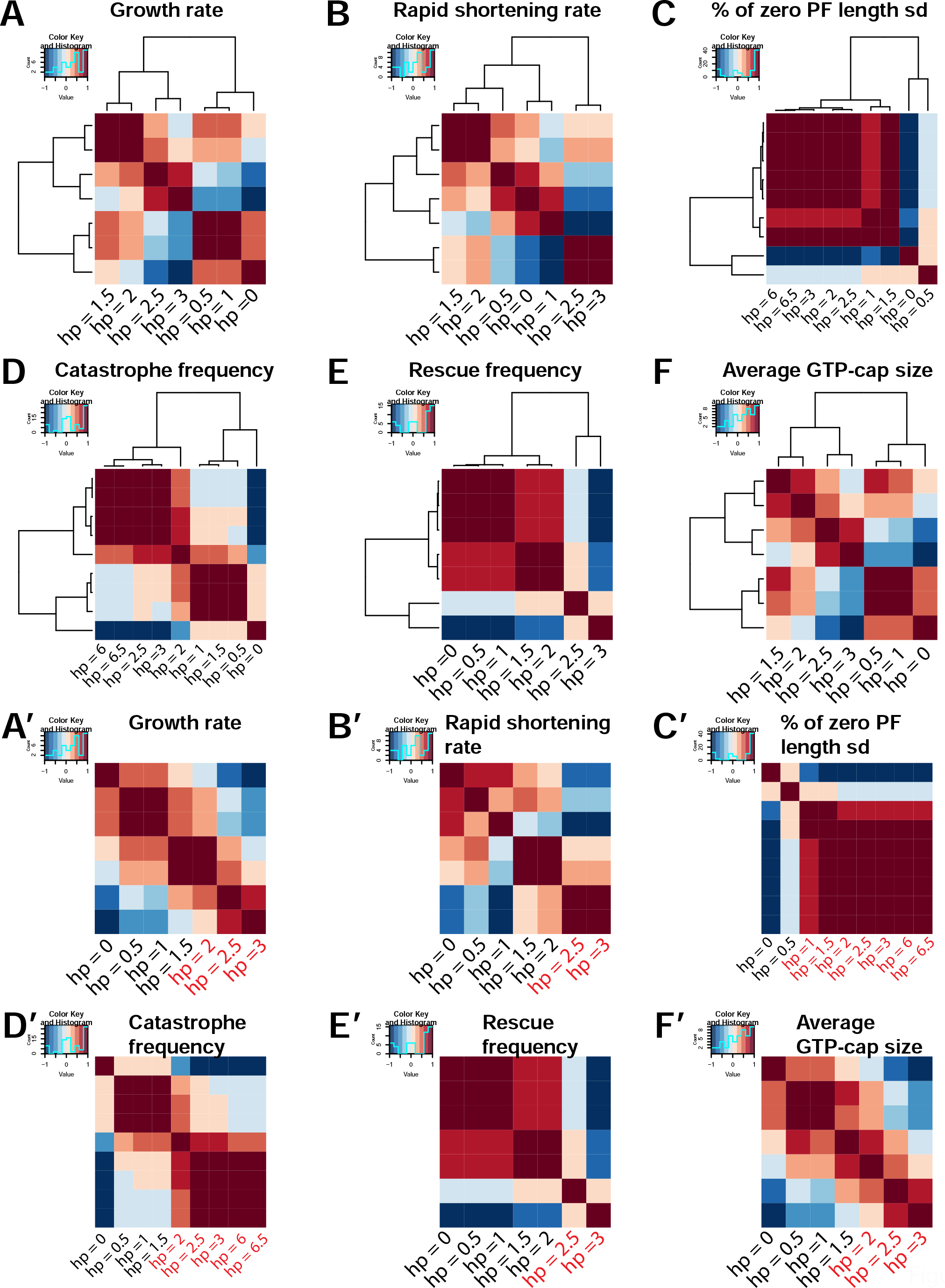
Hierarchical clustering and K-medoids clustering of aspects of MT plus end dynamic instability. Hierarchical clustering (A to F) and K-medoids clustering (A’ to F’) were performed to test if aspects of MT plus end dynamic instability was categorically determined by the presence or absence of seam (i.e. helical symmetry). Unsupervised clustering was carried out using both methods. For hierarchical clustering (complete linkage method), correlation matrix heatmaps and dendrograms were combined to show clustering outcome. For K-medoids clustering, correlation matrix heatmaps were shown by indexing on the output of the ordered cluster assignments (k = 2 was predetermined to test if MTs with seam and those without seam will form two clusters; black and red text labels indicate the two resultant clusters). The fact that helical pitch sizes with the same helical symmetry state didn’t cluster together (either with hierarchical or K-medoids method) indicate MT plus end dynamic instability is not categorically determined by the presence or absence of helical symmetry. Except for the percent of zero PF length standard deviation (“% of zero PF length sd”) and catastrophe frequency, hp = 6 and hp = 6.5 were not included due to obvious discontinuity that would confound the clustering results (see **Methods** for more details).

**Figure S7.**
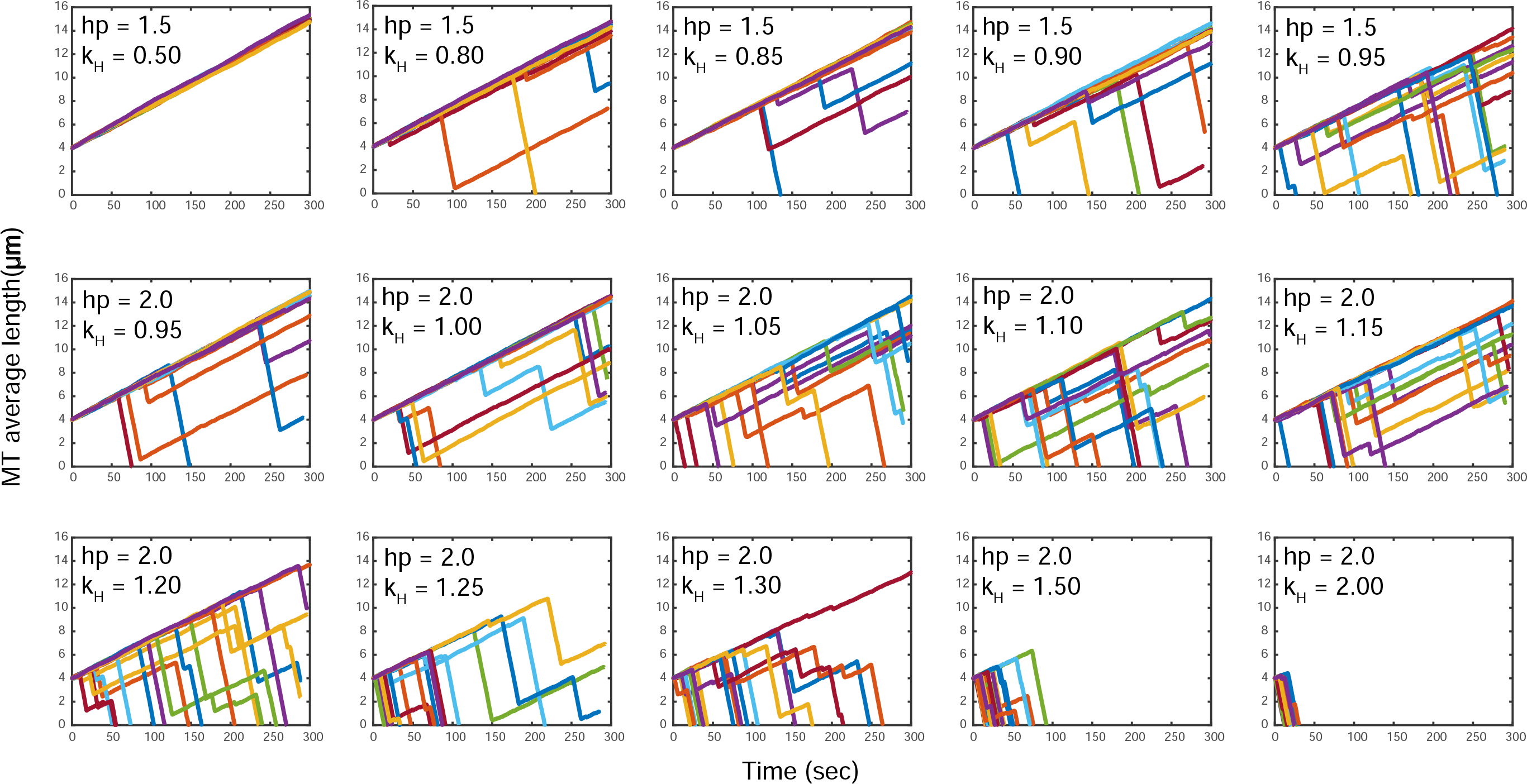
Modeling MT plus end dynamic instability by systematically altering hydrolysis rates at given certain helical pitch sizes. Graduated values of hydrolysis rates were screened across to simulate MT plus end dynamics under the condition of certain helical pitch size of either 1.5 or 2.0. Totally 18 simulation runs were plotted together under each condition.

**Figure S8.**
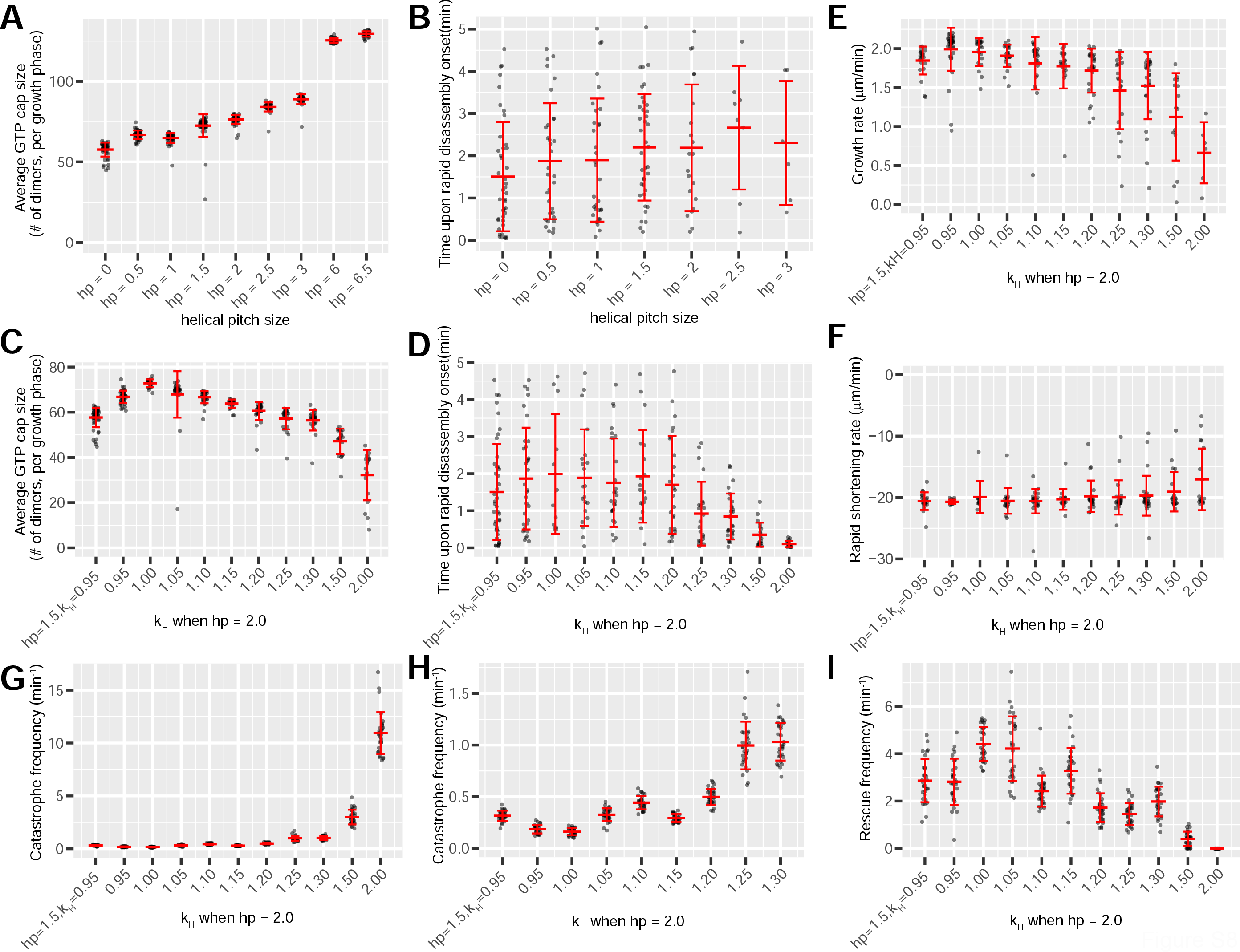
Raw scatterplots showing MT plus end dynamic instability as functions of helical pitch sizes or hydrolysis rates. (A and B) Scatter plot overlaid with Mean ± SD showing the average GTP-cap size (A) or time upon rapid disassembly onset (B), as functions of helical pitch sizes. For (A), the average size of total GTP-tubulin dimers in the entire MT (mean GTP cap size) during each growth phase was plotted as a function of helical pitch sizes. One-way ANOVA and post hoc pairwise t-test were used for assessing variations among and between helical pitch sizes. Krustal-Wallis and post hoc Wilcoxon pairwise tests were also performed for assessing variations among and between pitch sizes. For (B), Krustal-Wallis and post hoc Wilcoxon pairwise tests were performed for assessing variations among and between pitch sizes (see **Supplemental Statistics Data** for more details). (C and D) Scatter plot overlaid with Mean ± SD showing the average GTP-cap size (C) or time upon rapid disassembly onset (D), as functions of hydrolysis rates at an alternative helical pitch size of 2 dimers. The historical outcome (hp = 1.5, k_H_ = 0.95) was also plotted for comparison. For both (C) and (D), Krustal-Wallis and post hoc Wilcoxon pairwise tests were performed for assessing variations among and between pitch sizes (see **Supplemental Statistics Data** for more details). (E and F) Scatter plot overlaid with Mean ± SD showing the growth rate (E) or rapid shortening time (F), as functions of hydrolysis rates at an alternative helical pitch size of 2 dimers. The historical outcome (hp = 1.5, k_H_ = 0.95) was also plotted for comparison. For both (E) and (F), Krustal-Wallis and post hoc Wilcoxon pairwise tests were performed for assessing variations among and between pitch sizes (see **Supplemental Statistics Data** for more details). (G to I) Scatter plot overlaid with Mean ± SD showing the catastrophe frequency (G and H) or rescue frequency (I), as functions of hydrolysis rates at an alternative helical pitch size of 2 dimers. The historical outcome (hp = 1.5, k_H_ = 0.95) was also plotted for comparison. (H) is just a zoom-in of (G). For both (G) and (I), One-way ANOVA and post hoc pairwise t-test were used for assessing variations among and between helical pitch sizes (see **Supplemental Statistics Data** for more details).

**Figure S9.**
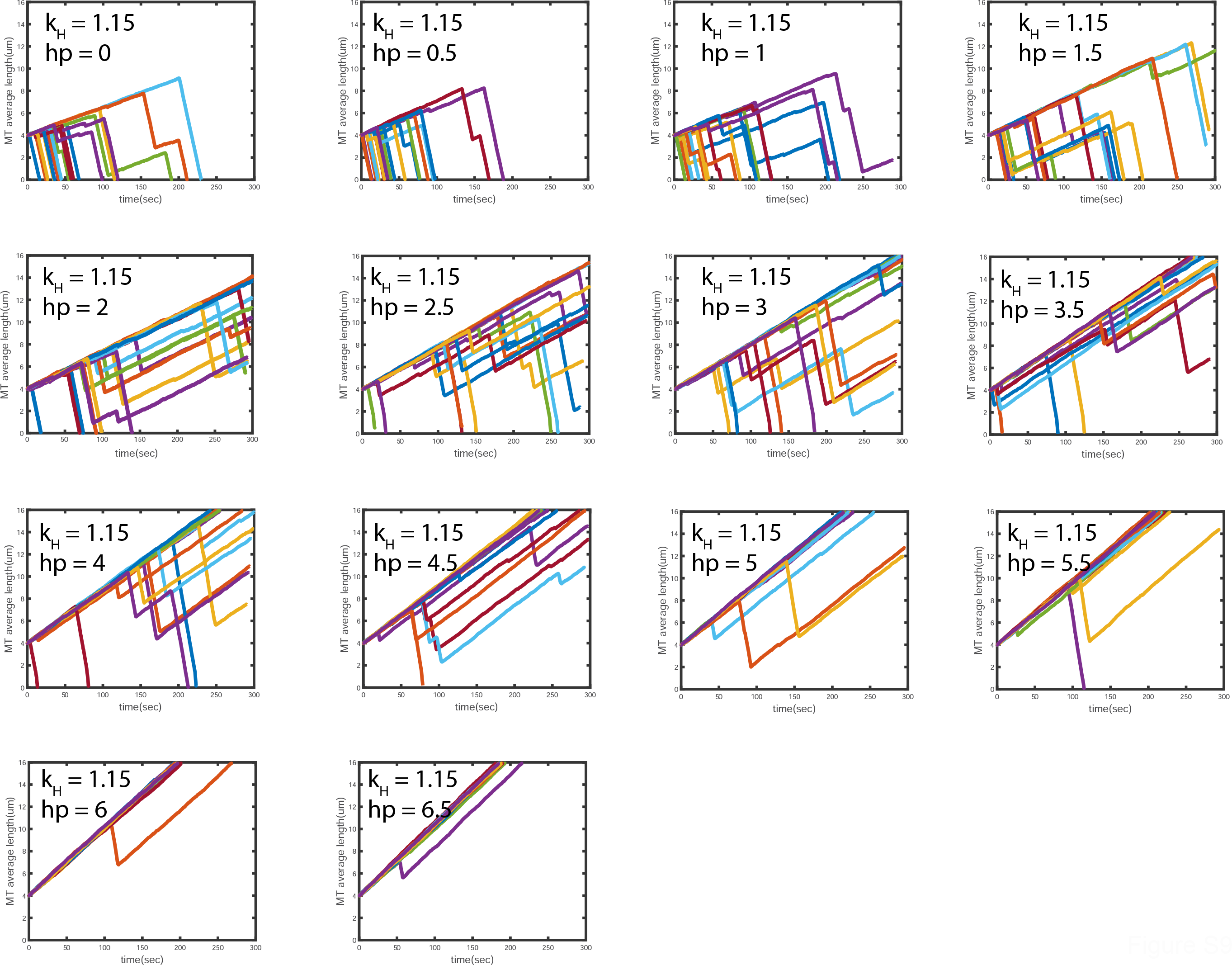
Modeling MT plus end dynamic instability by systematically altering helical pitch size at an altered hydrolysis rate (k_H_ = 1.15 molecule^-1^s^-1^). Graduated values of helical pitch sizes were screened across to simulate MT plus end dynamics under the condition of an alternative hydrolysis rate. Totally 18 simulation runs were plotted together under each condition. Helical pitches sized at 3.5, 4, 4.5, 5, 5.5 dimers were included here but not in earlier simulations because a clear gap of patterns between hp=3 and hp=6, observed here but not before.

**Figure S10.**
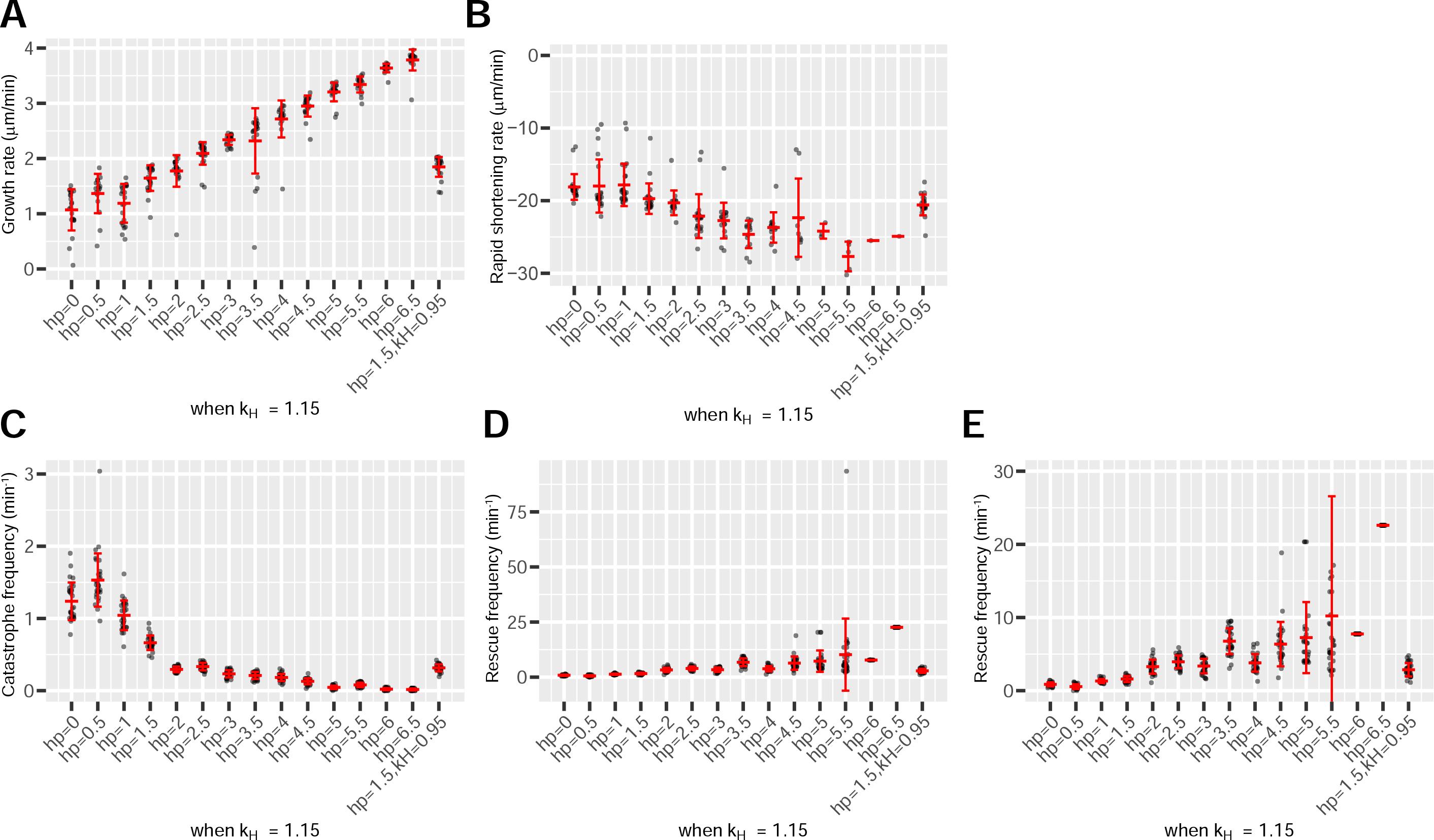
Raw scatterplots showing MT plus end dynamic instability as functions of helical pitch sizes at an altered hydrolysis rate (k_H_ = 1.15 molecule^-1^s^-1^). (A and B) Scatter plot overlaid with Mean ± SD showing the growth rate (A) or rapid shortening time (B), as functions of helical pitch sizes at an alternative hydrolysis rates (k_H_ = 1.15 molecule^- 1^s^-1^). The historical outcome (hp = 1.5, k_H_ = 0.95) was also plotted for comparison. For both (A) and (B), Krustal-Wallis and post hoc Wilcoxon pairwise tests were performed for assessing variations among and between pitch sizes (see **Supplemental Statistics Data** for more details). (C to E) Scatter plots overlaid with Mean ± SD showing the catastrophe frequency (C) or rescue frequency (D and E), as functions of helical pitch sizes at an alternative hydrolysis rates (k_H_ = 1.15 molecule^-1^s^-1^). The historical outcome (hp = 1.5, k_H_ = 0.95) was also plotted for comparison. (E) is just a zoom-in of (D) with re-scaled y-axis. For both (C) and (D), One-way ANOVA and post hoc pairwise t-test were used for assessing variations among and between helical pitch sizes (see **Supplemental Statistics Data** for more details).

**Figure S11.**
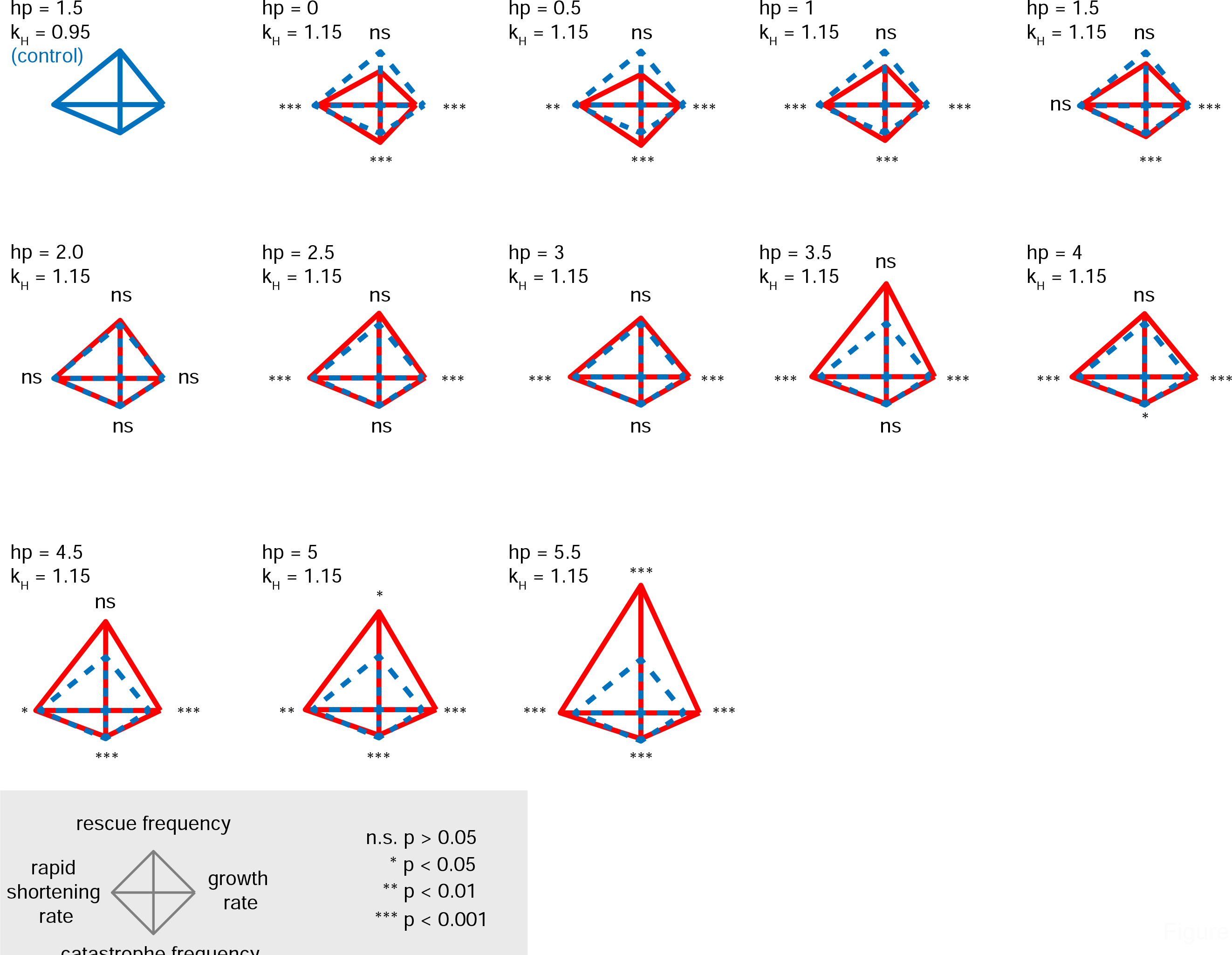
Diamond graphs showing aspects of MT plus end dynamic instability as functions of helical pitch sizes at an alternative hydrolysis rates (k_H_ = 1.15 molecule^-1^s^-1^). Diamond graphs using two pair of individually normalized parameters describing MT plus end dynamic instability. Mean values were plotted in the diamond graphs. With the historical outcome (hp = 1.5, k_H_ = 0.95) as control, Krustal-Wallis and post hoc Wilcoxon pairwise tests were performed for assessing variations among and between helical pitch sizes at an alternative hydrolysis rates (k_H_ = 1.15 molecule^-1^s^-1^) regarding rates. One-way ANOVA and post hoc pairwise t-tests were performed for assessing variations among and between different helical pitch sizes at an alternative hydrolysis rates (k_H_ = 1.15 molecule^-1^s^-1^) regarding frequencies (see **Methods**, and the **Supplemental Statistics Data** for more details). Diamond graphs for hp = 6 and hp = 6.5 are not included because some parameters have limited sample sizes.

**Figure S12.**
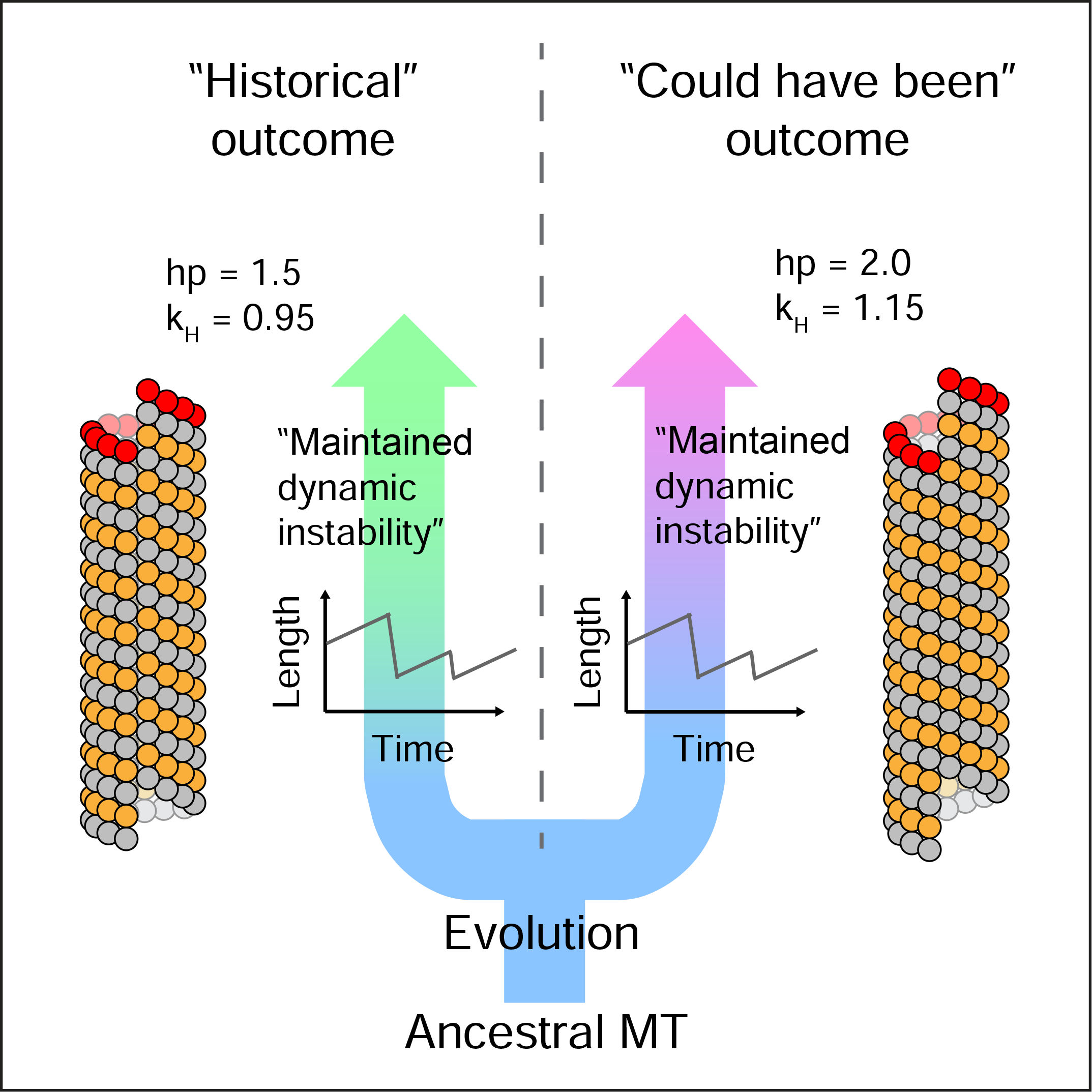
Schematics showing that while the “historical outcome” of 1.5 dimer helical pitch is conserved among most present-day MTs, this structural feature probably could have been bypassed if evolution had started differently. The plus end of the MTs are marked by red-colored β-tubulins.

## REFERENCES

1. Kline-Smith, S.L. and C.E. Walczak, Mitotic Spindle Assembly and Chromosome Segregation: Refocusing on Microtubule Dynamics. Molecular Cell, 2004. 15(3): p. 317–327.

2. Gardner, M.K. and D.J. Odde, Modeling of chromosome motility during mitosis. Current Opinion in Cell Biology, 2006. 18(6): p. 639–647.

3. Etienne-Manneville, S., Microtubules in Cell Migration. Annual Review of Cell and Developmental Biology, 2013. 29(1): p. 471–499.

4. Vale, R.D., Intracellular Transport Using Microtubule-Based Motors. Annual Review of Cell Biology, 1987. 3(1): p. 347–378.

5. Desai, A. and T.J. Mitchison, MICROTUBULE POLYMERIZATION DYNAMICS. Annual Review of Cell and Developmental Biology, 1997. 13(1): p. 83–117.

6. Prahl, L.S., et al., Microtubule-Based Control of Motor-Clutch System Mechanics in Glioma Cell Migration. Cell Reports, 2018. 25(9): p. 2591-2604.e8.

7. Wu, J. and A. Akhmanova, Microtubule-Organizing Centers. Annual Review of Cell and Developmental Biology, 2017. 33(1): p. 51–75.

8. Brouhard, G.J. and L.M. Rice, Microtubule dynamics: an interplay of biochemistry and mechanics. Nature Reviews Molecular Cell Biology, 2018. 19(7): p. 451–463.

9. Mitchison, T. and M. Kirschner, Dynamic instability of microtubule growth. Nature, 1984. 312(5991): p. 237–242.

10. Brouhard, G.J., Dynamic instability 30 years later: complexities in microtubule growth and catastrophe. Molecular biology of the cell, 2015. 26(7): p. 1207–1210.

11. Heald, R. and A. Khodjakov, Thirty years of search and capture: The complex simplicity of mitotic spindle assembly. Journal of Cell Biology, 2015. 211(6): p. 1103–1111.

12. Burdyniuk, M., et al., F-Actin nucleated on chromosomes coordinates their capture by microtubules in oocyte meiosis. Journal of Cell Biology, 2018. 217(8): p. 2661–2674.

13. Vaart, Babet van d., A. Akhmanova, and A. Straube, Regulation of microtubule dynamic instability. Biochemical Society Transactions, 2009. 37(5): p. 1007–1013.

14. Akhmanova, A. and M.O. Steinmetz, Tracking the ends: a dynamic protein network controls the fate of microtubule tips. Nat Rev Mol Cell Biol, 2008. 9(4): p. 309–322.

15. Amos, L.A. and D. Schlieper, Microtubules and Maps. Advances in Protein Chemistry, 2005. 71: p. 257–298.

16. Liu, C., et al., A dynein independent role of Tctex-1 at the kinetochore. Cell Cycle, 2015. 14(9): p. 1379–1388.

17. Schiff, P.B., J. Fant, and S.B. Horwitz, Promotion of microtubule assembly in vitro by taxol. Nature, 1979. 277(5698): p. 665–667.

18. Castle, B.T., et al., Mechanisms of kinetic stabilization by the drugs paclitaxel and vinblastine. Molecular Biology of the Cell, 2017. 28(9): p. 1238–1257.

19. Bowne-Anderson, H., et al., Microtubule dynamic instability: a new model with coupled GTP hydrolysis and multistep catastrophe. BioEssays: news and reviews in molecular, cellular and developmental biology, 2013. 35(5): p. 452–461.

20. Nogales, E., Structural Insights into Microtubule Function. Annual Review of Biochemistry, 2000. 69(1): p. 277–302.

21. Wang, H.-W. and E. Nogales, Nucleotide-dependent bending flexibility of tubulin regulates microtubule assembly. Nature, 2005. 435(7044): p. 911–915.

22. Alushin, Gregory M., et al., High-Resolution Microtubule Structures Reveal the Structural Transitions in αβ-Tubulin upon GTP Hydrolysis. Cell, 2014. 157(5): p. 1117–1129.

23. Mandelkow, E.M., E. Mandelkow, and R.A. Milligan, Microtubule dynamics and microtubule caps: a time-resolved cryo-electron microscopy study. Journal of Cell Biology, 1991. 114(5): p. 977–991.

24. Tilney, L.G., et al., Microtubules: evidence for 13 protofilaments. The Journal of cell biology, 1973. 59(2 Pt 1): p. 267–275.

25. McIntosh, J.R., et al., Lattice structure of cytoplasmic microtubules in a cultured Mammalian cell. Journal of molecular biology, 2009. 394(2): p. 177–182.

26. Hyman, A.A., et al., Role of GTP hydrolysis in microtubule dynamics: information from a slowly hydrolyzable analogue, GMPCPP. Molecular biology of the cell, 1992. 3(10): p. 1155–1167.

27. Mandelkow, E.M., et al., On the surface lattice of microtubules: helix starts, protofilament number, seam, and handedness. The Journal of Cell Biology, 1986. 102(3): p. 1067–1073.

28. Hunyadi, V., et al., Why is the microtubule lattice helical? Biology of the Cell, 2007. 099(2): p. 117–128.

29. Katsuki, M., D.R. Drummond, and R.A. Cross, Ectopic A-lattice seams destabilize microtubules. Nature Communications, 2014. 5: p. 3094.

30. Amos, L.A., Microtubule structure and its stabilisation. Organic & Biomolecular Chemistry, 2004. 2(15): p. 2153–2160.

31. Chrétien, D. and R.H. Wade, New data on the microtubule surface lattice. Biology of the Cell, 1991. 71(1-2): p. 161–174.

32. Pampaloni, F. and E.-L. Florin, Microtubule architecture: inspiration for novel carbon nanotube-based biomimetic materials. Trends in Biotechnology, 2008. 26(6): p. 302–310.

33. Ti, S.-C., G.M. Alushin, and T.M. Kapoor, Human β-Tubulin Isotypes Can Regulate Microtubule Protofilament Number and Stability. Developmental Cell, 2018. 47(2): p. 175-190.e5.

34. Zhang, R. and E. Nogales, A new protocol to accurately determine microtubule lattice seam location. Journal of Structural Biology, 2015. 192(2): p. 245–254.

35. Chaaban, S. and G.J. Brouhard, A microtubule bestiary: structural diversity in tubulin polymers. Molecular Biology of the Cell, 2017. 28(22): p. 2924–2931.

36. VanBuren, V., D.J. Odde, and L. Cassimeris, Estimates of lateral and longitudinal bond energies within the microtubule lattice. Proceedings of the National Academy of Sciences, 2002. 99(9): p. 6035–6040.

37. Gardner, Melissa K., et al., Rapid Microtubule Self-Assembly Kinetics. Cell, 2011. 146(4): p. 582–592.

38. Castle, Brian T. and David J. Odde, Brownian Dynamics of Subunit Addition-Loss Kinetics and Thermodynamics in Linear Polymer Self-Assembly. Biophysical Journal, 2013. 105(11): p. 2528–2540.

39. Walker, R.A., et al., Dynamic instability of individual microtubules analyzed by video light microscopy: rate constants and transition frequencies. The Journal of Cell Biology, 1988. 107(4): p. 1437–1448.

40. Walker, R.A., N.K. Pryer, and E.D. Salmon, Dilution of individual microtubules observed in real time in vitro: evidence that cap size is small and independent of elongation rate. The Journal of Cell Biology, 1991. 114(1): p. 73.

41. Drechsel, D.N. and M.W. Kirschner, The minimum GTP cap required to stabilize microtubules. Current Biology. 4(12): p. 1053–1061.

42. Saxton, W.M., et al., Tubulin dynamics in cultured mammalian cells. The Journal of Cell Biology, 1984. 99(6): p. 2175.

43. Zhai, Y., P.J. Kronebusch, and G.G. Borisy, Kinetochore microtubule dynamics and the metaphase-anaphase transition. The Journal of Cell Biology, 1995. 131(3): p. 721.

44. Vicente, J.J. and L. Wordeman, Mitosis, microtubule dynamics and the evolution of kinesins. Experimental Cell Research, 2015. 334(1): p. 61–69.

45. Gorbsky, G.J. and G.G. Borisy, Microtubules of the kinetochore fiber turn over in metaphase but not in anaphase. Journal of Cell Biology, 1989. 109(2): p. 653–662.

46. Bakhoum, S.F., et al., Genome stability is ensured by temporal control of kinetochore-microtubule dynamics. Nat Cell Biol, 2009. 11(1): p. 27–35.

47. VanBuren, V., L. Cassimeris, and D.J. Odde, Mechanochemical Model of Microtubule Structure and Self-Assembly Kinetics. Biophysical Journal, 2005. 89(5): p. 2911–2926.

48. Lacroix, B., et al., In Situ Imaging in C.elegans Reveals Developmental Regulation of Microtubule Dynamics. Developmental Cell, 2014. 29(2): p. 203–216.

49. Roostalu, J., et al., The speed of GTP hydrolysis determines GTP cap size and controls microtubule stability. eLife, 2020. 9: p. e51992.

50. Trogden, K.P. and S.L. Rogers, TOG Proteins Are Spatially Regulated by Rac-GSK3β to Control Interphase Microtubule Dynamics. PLOS ONE, 2015. 10(9): p. e0138966.

51. Pilhofer, M., et al., Microtubules in Bacteria: Ancient Tubulins Build a Five-Protofilament Homolog of the Eukaryotic Cytoskeleton. PLOS Biology, 2011. 9(12): p. e1001213.

52. Deng, X., et al., Four-stranded mini microtubules formed by <em>Prosthecobacter</em> BtubAB show dynamic instability. Proceedings of the National Academy of Sciences, 2017. 114(29): p. E5950.

53. Chrétien, D. and S.D. Fuller, Microtubules switch occasionally into unfavorable configurations during elongation1. Journal of Molecular Biology, 2000. 298(4): p. 663–676.

54. Kaye, J.S., THE FINE STRUCTURE AND ARRANGEMENT OF MICROCYLINDERS IN THE LUMINA OF FLAGELLAR FIBERS IN CRICKET SPERMATIDS. The Journal of Cell Biology, 1970. 45(2): p. 416.

55. Dallai, R. and B.A. Afzelius, Microtubular diversity in insect spermatozoa: Results obtained with a new fixative. Journal of Structural Biology, 1990. 103(2): p. 164–179.

56. Cueva, Juan G., et al., Posttranslational Acetylation of α-Tubulin Constrains Protofilament Number in Native Microtubules. Current Biology, 2012. 22(12): p. 1066–1074.

57. Manka, S.W. and C.A. Moores, Microtubule structure by cryo-EM: snapshots of dynamic instability. Essays in biochemistry, 2018. 62(6): p. 737–751.

58. Sui, H. and K.H. Downing, Structural Basis of Interprotofilament Interaction and Lateral Deformation of Microtubules. Structure, 2010. 18(8): p. 1022–1031.

59. Hemmat, M., et al., Multiscale Computational Modeling of Tubulin-Tubulin Lateral Interaction. Biophysical Journal, 2019. 117(7): p. 1234–1249.

60. Chaaban, S., et al., The Structure and Dynamics of <em>C. elegans</em> Tubulin Reveals the Mechanistic Basis of Microtubule Growth. Developmental Cell, 2018. 47(2): p. 191-204.e8.

61. Erickson, H.P. and W.A. Voter, Nucleation of Microtubule Assembly. Annals of the New York Academy of Sciences, 1986. 466(1): p. 552–565.

62. Cimini, D., et al., Merotelic kinetochore orientation occurs frequently during early mitosis in mammalian tissue cells and error correction is achieved by two different mechanisms. Journal of Cell Science, 2003. 116(20): p. 4213–4225.

63. Lampson, M.A. and I.M. Cheeseman, Sensing centromere tension: Aurora B and the regulation of kinetochore function. Trends in cell biology, 2011. 21(3): p. 133–140.

64. Kitajima, Tomoya S., M. Ohsugi, and J. Ellenberg, Complete Kinetochore Tracking Reveals Error-Prone Homologous Chromosome Biorientation in Mammalian Oocytes. Cell, 2011. 146(4): p. 568–581.

65. Starr, T.N., L.K. Picton, and J.W. Thornton, Alternative evolutionary histories in the sequence space of an ancient protein. Nature, 2017. 549(7672): p. 409–413.

66. McKeown, A.N., et al., Evolution of DNA specificity in a transcription factor family produced a new gene regulatory module. Cell, 2014. 159(1): p. 58–68.

67. Bloom, J.D., L.I. Gong, and D. Baltimore, Permissive secondary mutations enable the evolution of influenza oseltamivir resistance. Science (New York, N.Y.), 2010. 328(5983): p. 1272–1275.

68. Chalfie, M. and J.N. Thomson, Structural and functional diversity in the neuronal microtubules of Caenorhabditis elegans. The Journal of cell biology, 1982. 93(1): p. 15–23.

69. Savage, C., et al., mec-7 is a beta-tubulin gene required for the production of 15-protofilament microtubules in Caenorhabditis elegans. Genes & Development, 1989. 3(6): p. 870–881.

70. Fukushige, T., et al., MEC-12, an alpha-tubulin required for touch sensitivity in C. elegans. Journal of Cell Science, 1999. 112(3): p. 395.

71. Shah, P., D.M. McCandlish, and J.B. Plotkin, Contingency and entrenchment in protein evolution under purifying selection. Proceedings of the National Academy of Sciences, 2015. 112(25): p. E3226.

72. Gillespie, D.T., Exact stochastic simulation of coupled chemical reactions. The Journal of Physical Chemistry, 1977. 81(25): p. 2340–2361.

73. Martin, S.R., M.J. Schilstra, and P.M. Bayley, Dynamic instability of microtubules: Monte Carlo simulation and application to different types of microtubule lattice. Biophysical Journal, 1993. 65(2): p. 578–596.

74. Kutscheidt, S., et al., FHOD1 interaction with nesprin-2G mediates TAN line formation and nuclear movement. Nat Cell Biol, 2014. 16(7): p. 708–715.

75. Liu, C., R. Zhu, and Y. Mao, Nuclear Actin Polymerized by mDia2 Confines Centromere Movement during CENP-A Loading. iScience, 2018. 9: p. 314–327.

76. Liu, C. and Y. Mao, Diaphanous formin mDia2 regulates CENP-A levels at centromeres. The Journal of Cell Biology, 2016. 213(4): p. 415–424.

77. Sabine Landau, B.E.M.L., et al., Cluster Analysis. 2001: Wiley.

78. Wentzensen, N., et al., Hierarchical Clustering of Human Papilloma Virus Genotype Patterns in the ASCUS-LSIL Triage Study. Cancer Research, 2010. 70(21): p. 8578.

79. Franzén, O., et al., Improved OTU-picking using long-read 16S rRNA gene amplicon sequencing and generic hierarchical clustering. Microbiome, 2015. 3(1): p. 43.

80. Schmidt, T.S.B., J.F. Matias Rodrigues, and C. von Mering, Ecological Consistency of SSU rRNA-Based Operational Taxonomic Units at a Global Scale. PLOS Computational Biology, 2014. 10(4): p. e1003594.

81. de Brevern, A.G., S. Hazout, and A. Malpertuy, Influence of microarrays experiments missing values on the stability of gene groups by hierarchical clustering. BMC Bioinformatics, 2004. 5(1): p. 114.

82. Leonard Kaufman, P.R., Clustering by Means of medoids. Statistical Data Analysis Based on the L1-Norm and Related Methods, 1987: p. 405–416.

83. Johnson, M.G., et al., A Universal Probe Set for Targeted Sequencing of 353 Nuclear Genes from Any Flowering Plant Designed Using k-Medoids Clustering. Systematic Biology, 2018. 68(4): p. 594–606.

84. Brennecke, P., et al., Single-cell transcriptome analysis reveals coordinated ectopic gene-expression patterns in medullary thymic epithelial cells. Nature Immunology, 2015. 16(9): p. 933–941.

