## Supplemental Statistics Data for "The size of helical pitch is important for microtubule plus end dynamic instability"

### Table of content:

Output of statistical analyses for data presented in main figures and supplemental figures are as following:

| **Output of statistical analyses** | **Figure(s)** |
|---|---|
| Statistics on the growth phase velocity (growth rate, μm/min) as a function of hp (with all $k_H$ = 0.95) | Figure 3A |
| Statistics on the rapid shortening phase velocity (rapid shortening rate, μm/min) as a function of hp (with all $k_H$ = 0.95) | Figure 3B |
| Statistics on the catastrophe frequency as a function of hp (with all $k_H$ = 0.95) | Figure 3C |
| Statistics on the rescue frequency as a function of hp (with all $k_H$ = 0.95) | Figure 3D |
| Statistics on the percentage of zero protofilament (PF) length standard deviation as a function of hp (with all $k_H$ = 0.95) | Figure 3E |
| Statistics on the proportion of resilient growth as a function of hp | Figure 3F |
| Statistics on the proportion of resilient growth as a function of $k_H$ (when hp = 2.0) with hp = 1.5, $k_H$ = 0.95 as control | Figure 4B |
| Statistics on the proportion of resilient growth as a function of hp (when $k_H$ = 1.15) with hp = 1.5, $k_H$ = 0.95 as control | Figure 4C |
| Statistics on the average GTP cap size (# of dimers, per growth phase) as a function of hp (with all $k_H$ = 0.95) | Figure S8A |
| Statistics on time upon rapid disassembly onset(min) as a function of hp (with all $k_H$ = 0.95) | Figure S8B |
| Statistics on the average GTP cap size (# of dimers, per growth phase) as a function of $k_H$ (when hp = 2.0) with hp = 1.5, $k_H$ = 0.95 as control | Figure S8C |
| Statistics on time upon rapid disassembly onset(min) as a function of $k_H$ (when hp = 2.0) with hp = 1.5, $k_H$ = 0.95 as control | Figure S8D |
| Statistics on the growth phase velocity as a function of $k_H$ (when hp = 2.0) with hp = 1.5, $k_H$ = 0.95 as control | Figure 4A, Figure S8E |
| Statistics on the rapid shortening phase velocity as a function of $k_H$ (when hp = 2.0) with hp = 1.5, $k_H$ = 0.95 as control | Figure 4A, Figure S8F |
| Statistics on the catastrophe frequency as a function of $k_H$ (when hp = 2.0) with hp = 1.5, $k_H$ = 0.95 as control | Figure 4A, Figure S8G, S8H |
| Statistics on the rescue frequency as a function of $k_H$ (when hp = 2.0) with hp = 1.5, $k_H$ = 0.95 as control | Figure 4A, Figure S8I |
| Statistics on the growth phase velocity as a function of hp (when $k_H$ = 1.15) with hp = 1.5, $k_H$ = 0.95 as control | Figure S10A, S11 |
| Statistics on the rapid shortening phase velocity as a function of hp (when $k_H$ = 1.15) with hp = 1.5, $k_H$ = 0.95 as control | Figure S10B, S11 |
| Statistics on the catastrophe frequency as a function of hp (when $k_H$ = 1.15) with hp = 1.5, $k_H$ = 0.95 as control | Figure S10C, S11 |
| Statistics on the rescue frequency as a function of hp (when $k_H$ = 1.15) with hp = 1.5, $k_H$ = 0.95 as control | Figure S10D, S10E, S11 |

Statistics on the growth phase velocity (growth rate, µm/min) as a function of hp (with all $k_H$ = 0.95)

Sample means:

```
    hp=0   hp=0.5     hp=1   hp=1.5     hp=2   hp=2.5     hp=3     hp=6   hp=6.5 
1.270376 1.580965 1.535697 1.868835 1.992308 2.330100 2.533078 3.805731 3.948211 
```

sample standard deviations:

```
      hp=0     hp=0.5       hp=1     hp=1.5       hp=2     hp=2.5       hp=3       hp=6 
0.31726654 0.28838727 0.23991835 0.15894604 0.27570592 0.20242206 0.06122140 0.05201569 
    hp=6.5 
0.06204380 
```

sample sizes:

```
hp=0   hp=0.5     hp=1   hp=1.5     hp=2   hp=2.5     hp=3     hp=6   hp=6.5 
  40       42       41       43       39       36       37       36       36 
```

Kruskal-Wallis rank sum test

```
data:  dat15$V_g by id
Kruskal-Wallis chi-squared = 317.74, df = 8, p-value < 2.2e-16
```

Pairwise comparisons using Wilcoxon rank sum test

```
data:  dat15$V_g and id 

       hp=0    hp=0.5  hp=1    hp=1.5  hp=2    hp=2.5  hp=3    hp=6   
hp=0.5 3.2e-09 -       -       -       -       -       -       -      
hp=1   2.5e-08 0.021   -       -       -       -       -       -      
hp=1.5 < 2e-16 1.2e-12 5.2e-14 -       -       -       -       -      
hp=2   6.0e-15 4.8e-13 4.5e-13 1.0e-07 -       -       -       -      
hp=2.5 < 2e-16 < 2e-16 < 2e-16 < 2e-16 < 2e-16 -       -       -      
hp=3   < 2e-16 < 2e-16 < 2e-16 < 2e-16 < 2e-16 2.6e-16 -       -      
hp=6   < 2e-16 < 2e-16 < 2e-16 < 2e-16 < 2e-16 < 2e-16 < 2e-16 -      
hp=6.5 < 2e-16 < 2e-16 < 2e-16 < 2e-16 < 2e-16 < 2e-16 < 2e-16 1.5e-14
P value adjustment method: BH 
```

Statistics on the rapid shortening phase velocity (rapid shortening rate, µm/min) as a function of hp (with all $k_H$ = 0.95)

Sample means:
```
      hp=0    hp=0.5      hp=1    hp=1.5      hp=2    hp=2.5      hp=3
-18.56246 -19.39622 -18.02959 -20.28878 -20.70298 -22.36629 -22.05768
```
sample standard deviations:
```
     hp=0    hp=0.5      hp=1    hp=1.5      hp=2    hp=2.5      hp=3
0.4565820 0.7278660 3.0523940 1.6630858 0.3458078 1.7633153 1.9078037
```

sample sizes:
```
     hp=0    hp=0.5      hp=1    hp=1.5      hp=2    hp=2.5      hp=3
       40        31        28        35        18         8         5
```

Kruskal-Wallis rank sum test
```
data:  dat16$V_rs by id
Kruskal-Wallis chi-squared = 102.05, df = 6, p-value < 2.2e-16
```

Pairwise comparisons using Wilcoxon rank sum test
```
data:  dat16$V_rs and id
```

| | hp=0 | hp=0.5 | hp=1 | hp=1.5 | hp=2 | hp=2.5 |
|---|---|---|---|---|---|---|
| hp=0.5 | 1.1e-08 | - | - | - | - | - |
| hp=1 | 0.0062 | 0.0742 | - | - | - | - |
| hp=1.5 | 1.4e-11 | 7.0e-05 | 9.5e-07 | - | - | - |
| hp=2 | 9.3e-14 | 1.6e-08 | 9.7e-10 | 0.2799 | - | - |
| hp=2.5 | 3.7e-08 | 8.7e-05 | 2.6e-05 | 0.0015 | 0.0024 | - |
| hp=3 | 4.3e-06 | 4.1e-05 | 2.0e-05 | 0.0189 | 0.0062 | 0.4351 |

P value adjustment method: BH

Statistics on the catastrophe frequency as a function of hp (with all $k_H$ = 0.95)

sample means:

```
      hp=0     hp=0.5       hp=1     hp=1.5       hp=2     hp=2.5       hp=3       hp=6     hp=6.5 
0.51120147 0.29385814 0.25926083 0.26799410 0.13014901 0.04400852 0.03540214 0.00000000 0.00000000 
```

sample standard deviations:

```
      hp=0     hp=0.5       hp=1     hp=1.5       hp=2     hp=2.5       hp=3       hp=6 
0.13031532 0.08953915 0.05694861 0.06671413 0.06750933 0.02301158 0.02170364 0.00000000 
    hp=6.5 
0.00000000 
```

sample sizes:

```
      hp=0     hp=0.5       hp=1     hp=1.5       hp=2     hp=2.5       hp=3       hp=6 
        60         60         60         60         60         60         60         30 
    hp=6.5 
        60 
```

One-way ANOVA:

```
             Df Sum Sq Mean Sq F value Pr(>F)    
hp_id         8 13.629  1.7037   378.2 <2e-16 ***
Residuals   501  2.257  0.0045                   
---
Signif. codes:  0 ‘***’ 0.001 ‘**’ 0.01 ‘*’ 0.05 ‘.’ 0.1 ‘ ’ 1
```

Pairwise comparisons using t tests with pooled SD

```
data:  df_raw_comb$F_cat and hp_id 
       hp=0    hp=0.5  hp=1    hp=1.5  hp=2    hp=2.5 hp=3  hp=6 
hp=0.5 < 2e-16 -       -       -       -       -      -     -    
hp=1   < 2e-16 0.178   -       -       -       -      -     -    
hp=1.5 < 2e-16 1.000   1.000   -       -       -      -     -    
hp=2   < 2e-16 < 2e-16 < 2e-16 < 2e-16 -       -      -     -    
hp=2.5 < 2e-16 < 2e-16 < 2e-16 < 2e-16 2.5e-10 -      -     -    
hp=3   < 2e-16 < 2e-16 < 2e-16 < 2e-16 2.1e-12 1.000  -     -    
hp=6   < 2e-16 < 2e-16 < 2e-16 < 2e-16 2.1e-15 0.127  0.674 -    
hp=6.5 < 2e-16 < 2e-16 < 2e-16 < 2e-16 < 2e-16 0.013  0.145 1.000
P value adjustment method: bonferroni 
```

Statistics on the rescue frequency as a function of hp (with all $k_H$ = 0.95)

sample means:

```
   hp=0   hp=0.5     hp=1   hp=1.5     hp=2   hp=2.5     hp=3
1.931931 1.880937 1.811089 2.789971 2.673283 5.998727 8.838105
```

sample standard deviations:

```
     hp=0    hp=0.5      hp=1    hp=1.5      hp=2    hp=2.5      hp=3
0.8535576 0.8922721 0.5448699 0.9374734 1.3872433 4.9973357 7.5233288
```

sample sizes:

```
hp=0    hp=0.5      hp=1    hp=1.5      hp=2    hp=2.5      hp=3
  60        60        60        60        60        59        56
```

One-way ANOVA:

```
             Df Sum Sq Mean Sq F value Pr(>F)
hp_id         6   2502   417.0    35.2 <2e-16 ***
Residuals   408   4834    11.8
---
Signif. codes:  0 ‘***’ 0.001 ‘**’ 0.01 ‘*’ 0.05 ‘.’ 0.1 ‘ ’ 1
```

Pairwise comparisons using t tests with pooled SD

```
data:  df_raw_comb1$F_res and hp_id
       hp=0    hp=0.5  hp=1    hp=1.5  hp=2    hp=2.5
hp=0.5 1.00000 -       -       -       -       -
hp=1   1.00000 1.00000 -       -       -       -
hp=1.5 1.00000 1.00000 1.00000 -       -       -
hp=2   1.00000 1.00000 1.00000 1.00000 -       -
hp=2.5 6.9e-09 4.2e-09 2.2e-09 1.2e-05 4.7e-06 -
hp=3   < 2e-16 < 2e-16 < 2e-16 < 2e-16 < 2e-16 0.00026
P value adjustment method: bonferroni
```

### Statistics on the percentage of zero protofilament (PF) length standard deviation as a function of hp (with all $k_H$ = 0.95)

Sample means:

| hp=0 | hp=0.5 | hp=1 | hp=1.5 | hp=2 | hp=2.5 | hp=3 | hp=6 | hp=6.5 |
|---|---|---|---|---|---|---|---|---|
| 13.01411444 | 7.43130211 | 1.85465272 | 1.22093611 | 0.46108377 | 0.32015625 | 0.13814359 | 0.02152778 | 0.01103935 |

sample standard deviations:

| hp=0 | hp=0.5 | hp=1 | hp=1.5 | hp=2 | hp=2.5 | hp=3 | hp=6 | hp=6.5 |
|---|---|---|---|---|---|---|---|---|
| 0.890902911 | 0.387349597 | 0.133793787 | 0.100221864 | 0.052947790 | 0.053197966 | 0.037740987 | 0.007663572 | 0.007176525 |

sample sizes:
n = 18 for all conditions

Kruskal-Wallis rank sum test
data:  dat11$percent_std by hp_id
Kruskal-Wallis chi-squared = 158.08, df = 8, p-value < 2.2e-16

Pairwise comparisons using Wilcoxon rank sum test

data:  dat11$percent_std and hp_id

| | hp=0 | hp=0.5 | hp=1 | hp=1.5 | hp=2 | hp=2.5 | hp=3 | hp=6 |
|---|---|---|---|---|---|---|---|---|
| hp=0.5 | 3.4e-07 | - | - | - | - | - | - | - |
| hp=1 | 1.3e-09 | 3.4e-07 | - | - | - | - | - | - |
| hp=1.5 | 1.3e-09 | 3.4e-07 | 1.3e-09 | - | - | - | - | - |
| hp=2 | 3.4e-07 | 3.4e-07 | 3.4e-07 | 3.4e-07 | - | - | - | - |
| hp=2.5 | 1.3e-09 | 3.4e-07 | 1.3e-09 | 1.3e-09 | 3.7e-06 | - | - | - |
| hp=3 | 3.4e-07 | 3.4e-07 | 3.4e-07 | 3.4e-07 | 3.4e-07 | 3.4e-07 | - | - |
| hp=6 | 3.4e-07 | 3.4e-07 | 3.4e-07 | 3.4e-07 | 3.4e-07 | 3.4e-07 | 3.4e-07 | - |
| hp=6.5 | 3.4e-07 | 3.4e-07 | 3.4e-07 | 3.4e-07 | 3.4e-07 | 3.4e-07 | 3.4e-07 | 0.00047 |

P value adjustment method: BH

Statistics on the proportion of resilient growth as a function of hp (treated as continuous quantitative variable)

Sample means:

```
     hp=0    hp=0.5      hp=1    hp=1.5      hp=2    hp=2.5      hp=3      hp=6    hp=6.5
0.2960784 0.2333333 0.3729167 0.5814815 0.2018519 0.1555556 0.1111111 0.0000000 0.0000000
```

Sample standard deviations:

```
      hp=0     hp=0.5       hp=1     hp=1.5       hp=2     hp=2.5       hp=3       hp=6
0.13332041 0.11297343 0.12225800 0.15002010 0.08812330 0.09040847 0.07720093 0.00000000
    hp=6.5
0.00000000
```

Sample sizes:

n = 30 for all conditions

One-way ANOVA:

```
             Df Sum Sq Mean Sq F value Pr(>F)
hp_id         8  8.192   1.024   102.5 <2e-16 ***
Residuals   261  2.607   0.010
---
Signif. codes:  0 ‘***’ 0.001 ‘**’ 0.01 ‘*’ 0.05 ‘.’ 0.1 ‘ ’ 1
```

Pairwise comparisons using t tests with pooled SD

```
data:  df_raw_0$percent_r and hp_id
       hp=0    hp=0.5  hp=1    hp=1.5  hp=2    hp=2.5  hp=3    hp=6
hp=0.5 0.56541 -       -       -       -       -       -       -
hp=1   0.11436 5.1e-06 -       -       -       -       -       -
hp=1.5 < 2e-16 < 2e-16 8.5e-13 -       -       -       -       -
hp=2   0.01133 1.00000 6.9e-09 < 2e-16 -       -       -       -
hp=2.5 4.3e-06 0.10190 8.9e-14 < 2e-16 1.00000 -       -       -
hp=3   2.8e-10 0.00013 < 2e-16 < 2e-16 0.01856 1.00000 -       -
hp=6   < 2e-16 1.3e-15 < 2e-16 < 2e-16 4.6e-12 2.0e-07 0.00085 -
hp=6.5 < 2e-16 1.3e-15 < 2e-16 < 2e-16 4.6e-12 2.0e-07 0.00085 1.00000
P value adjustment method: bonferroni
```

Statistics on the proportion of resilient growth as a function of hp
(1, hp=0; 2, hp=0.5; 3, hp=1; 4, hp=1.5; 5, hp=2; 6, hp=2.5; 7, hp=3; 8, hp=6; 9, hp=6.5)

9-sample test for equality of proportions without continuity correction:

data: count of resilient growth out of total count
X-squared = 865.62, df = 8, p-value < 2.2e-16
alternative hypothesis: two.sided

sample estimates:

| prop 1 | prop 2 | prop 3 | prop 4 | prop 5 | prop 6 | prop 7 |
|---|---|---|---|---|---|---|
| 0.2960784 | 0.2333333 | 0.3729167 | 0.5814815 | 0.2018519 | 0.1555556 | 0.1111111 |

| prop 8 | prop 9 |
|---|---|
| 0.0000000 | 0.0000000 |

sample size:

n = 30 for each hp condition

Post hoc pairwise comparison of proportions:

data: count of resilient growth out of total count

| | 1 | 2 | 3 | 4 | 5 | 6 | 7 | 8 |
|---|---|---|---|---|---|---|---|---|
| 2 | 0.973 | - | - | - | - | - | - | - |
| 3 | 0.440 | 8.4e-05 | - | - | - | - | - | - |
| 4 | < 2e-16 | < 2e-16 | 1.5e-09 | - | - | - | - | - |
| 5 | 0.019 | 1.000 | 7.4e-08 | < 2e-16 | - | - | - | - |
| 6 | 2.5e-06 | 0.065 | 1.5e-13 | < 2e-16 | 1.000 | - | - | - |
| 7 | 4.8e-12 | 7.7e-06 | < 2e-16 | < 2e-16 | 0.002 | 1.000 | - | - |
| 8 | < 2e-16 | < 2e-16 | < 2e-16 | < 2e-16 | < 2e-16 | < 2e-16 | 1.6e-13 | - |
| 9 | < 2e-16 | < 2e-16 | < 2e-16 | < 2e-16 | < 2e-16 | < 2e-16 | 1.6e-13 | - |

P value adjustment method: bonferroni

Statistics on the proportion of resilient growth as a function of $k_H$ (when hp = 2.0) with hp = 1.5, $k_H$ = 0.95 as control (treated as continuous quantitative variable)

Sample means:

| kH=0.95 | kH=1.00 | kH=1.05 | kH=1.10 | kH=1.15 | kH=1.20 |
|---|---|---|---|---|---|
| 0.20185185 | 0.32500000 | 0.32444444 | 0.29607843 | 0.50625000 | 0.04666667 |
| **kH=1.25** | **kH=1.30** | **kH=1.50** | **kH=2.00** | **hp=1.5,kH=0.95** | |
| 0.11568627 | 0.04444444 | 0.00000000 | 0.00000000 | 0.58148148 | |

Sample standard deviations:

| kH=0.95 | kH=1.00 | kH=1.05 | kH=1.10 | kH=1.15 | kH=1.20 |
|---|---|---|---|---|---|
| 0.08812330 | 0.11767320 | 0.11577933 | 0.13332041 | 0.13369540 | 0.05846045 |
| **kH=1.25** | **kH=1.30** | **kH=1.50** | **kH=2.00** | **hp=1.5,kH=0.95** | |
| 0.09071345 | 0.05137530 | 0.00000000 | 0.00000000 | 0.15002010 | |

Sample sizes:

n = 30 for all conditions

One-way ANOVA:

| | Df | Sum Sq | Mean Sq | F value | Pr(>F) | |
|---|---|---|---|---|---|---|
| hp_id | 10 | 12.275 | 1.2275 | 125.7 | <2e-16 | *** |
| Residuals | 319 | 3.116 | 0.0098 | | | |

---

Signif. codes:  0 ‘***’ 0.001 ‘**’ 0.01 ‘*’ 0.05 ‘.’ 0.1 ‘ ’ 1

Pairwise comparisons using t tests with pooled SD

data:  df_raw_0$percent_r and hp_id

| | kH=0.95 | kH=1.00 | kH=1.05 | kH=1.10 | kH=1.15 | kH=1.20 | kH=1.25 | kH=1.30 | kH=1.50 | kH=2.00 |
|---|---|---|---|---|---|---|---|---|---|---|
| kH=1.00 | 0.00012 | - | - | - | - | - | - | - | - | - |
| kH=1.05 | 0.00013 | 1.00000 | - | - | - | - | - | - | - | - |
| kH=1.10 | 0.01437 | 1.00000 | 1.00000 | - | - | - | - | - | - | - |
| kH=1.15 | < 2e-16 | 4.4e-10 | 3.9e-10 | 2.6e-13 | - | - | - | - | - | - |
| kH=1.20 | 1.9e-07 | < 2e-16 | < 2e-16 | < 2e-16 | < 2e-16 | - | - | - | - | - |
| kH=1.25 | 0.04539 | 3.2e-13 | 3.7e-13 | 5.4e-10 | < 2e-16 | 0.39638 | - | - | - | - |
| kH=1.30 | 1.2e-07 | < 2e-16 | < 2e-16 | < 2e-16 | < 2e-16 | 1.00000 | 0.30583 | - | - | - |
| kH=1.50 | 2.3e-12 | < 2e-16 | < 2e-16 | < 2e-16 | < 2e-16 | 1.00000 | 0.00045 | 1.00000 | - | - |
| kH=2.00 | 2.3e-12 | < 2e-16 | < 2e-16 | < 2e-16 | < 2e-16 | 1.00000 | 0.00045 | 1.00000 | 1.00000 | - |
| hp=1.5,kH=0.95 | < 2e-16 | < 2e-16 | < 2e-16 | < 2e-16 | 0.18891 | < 2e-16 | < 2e-16 | < 2e-16 | < 2e-16 | < 2e-16 |

P value adjustment method: bonferroni

Statistics on the proportion of resilient growth as a function of $k_H$ (when hp = 2.0) with hp = 1.5, $k_H$ = 0.95 as control

(1, $k_H$=0.95; 2, $k_H$=1.00; 3, $k_H$=1.05; 4, $k_H$=1.10; 5, $k_H$=1.15; 6, $k_H$=1.20; 7, $k_H$=1.25; 8, $k_H$=1.30; 9, $k_H$=1.50; 10, $k_H$=2.00; 11, hp=1.5, $k_H$=0.95)

11-sample test for equality of proportions without continuity correction:

data: count of resilient growth out of total count
X-squared = 1234.1, df = 10, p-value < 2.2e-16
alternative hypothesis: two.sided

sample estimates:

| prop 1 | prop 2 | prop 3 | prop 4 | prop 5 | prop 6 | prop 7 |
|---|---|---|---|---|---|---|
| 0.20185185 | 0.32500000 | 0.32444444 | 0.29607843 | 0.50625000 | 0.04666667 | 0.11568627 |

| prop 8 | prop 9 | prop 10 | prop 11 |
|---|---|---|---|
| 0.04444444 | 0.00000000 | 0.00000000 | 0.58148148 |

sample size:

n = 30 for each hp condition

Post hoc Pairwise comparison of proportions:

data: count of resilient growth out of total count

| | 1 | 2 | 3 | 4 | 5 | 6 | 7 | 8 | 9 | 10 |
|---|---|---|---|---|---|---|---|---|---|---|
| 2 | 0.00057 | - | - | - | - | - | - | - | - | - |
| 3 | 0.00085 | 1.00000 | - | - | - | - | - | - | - | - |
| 4 | 0.02874 | 1.00000 | 1.00000 | - | - | - | - | - | - | - |
| 5 | < 2e-16 | 9.6e-07 | 1.5e-06 | 1.2e-09 | - | - | - | - | - | - |
| 6 | 6.5e-11 | < 2e-16 | < 2e-16 | < 2e-16 | < 2e-16 | - | - | - | - | - |
| 7 | 0.01066 | 1.4e-13 | 3.4e-13 | 9.9e-11 | < 2e-16 | 0.00978 | - | - | - | - |
| 8 | 4.0e-13 | < 2e-16 | < 2e-16 | < 2e-16 | < 2e-16 | 1.00000 | 0.00171 | - | - | - |
| 9 | < 2e-16 | < 2e-16 | < 2e-16 | < 2e-16 | < 2e-16 | 6.6e-05 | 6.6e-14 | 0.00011 | - | - |
| 10 | < 2e-16 | < 2e-16 | < 2e-16 | < 2e-16 | < 2e-16 | 6.6e-05 | 6.6e-14 | 0.00011 | - | - |
| 11 | < 2e-16 | 2.1e-14 | 6.2e-14 | < 2e-16 | 1.00000 | < 2e-16 | < 2e-16 | < 2e-16 | < 2e-16 | < 2e-16 |

P value adjustment method: bonferroni

Statistics on the proportion of resilient growth as a function of hp (when $k_H$ = 1.15) with hp = 1.5, $k_H$ = 0.95 as control (treated as continuous quantitative variable)

Sample means:

```
        hp=0        hp=0.5          hp=1        hp=1.5          hp=2        hp=2.5
  0.00000000    0.00000000    0.06666667    0.07916667    0.50625000    0.48627451
        hp=3        hp=3.5          hp=4        hp=4.5          hp=5        hp=5.5
  0.52407407    0.40588235    0.37962963    0.31111111    0.16666667    0.20740741
        hp=6        hp=6.5 hp=1.5,kH=0.95
  0.06666667    0.06666667    0.58148148
```

Sample standard deviations:

```
        hp=0        hp=0.5          hp=1        hp=1.5          hp=2        hp=2.5
  0.00000000    0.00000000    0.06258243    0.06950668    0.13369540    0.12058535
        hp=3        hp=3.5          hp=4        hp=4.5          hp=5        hp=5.5
  0.10788746    0.12197907    0.11022958    0.13810121    0.09787004    0.12544797
        hp=6        hp=6.5 hp=1.5,kH=0.95
  0.06258243    0.06258243    0.15002010
```

Sample sizes:

n = 30 for all conditions

One-way ANOVA:

```
             Df Sum Sq Mean Sq F value Pr(>F)
hp_id        14 18.434  1.3167   127.8 <2e-16 ***
Residuals   435  4.482  0.0103
---
Signif. codes:  0 ‘***’ 0.001 ‘**’ 0.01 ‘*’ 0.05 ‘.’ 0.1 ‘ ’ 1
```

Pairwise comparisons using t tests with pooled SD

```
data:  df_raw_0$percent_r and hp_id
               hp=0    hp=0.5  hp=1    hp=1.5  hp=2    hp=2.5  hp=3    hp=3.5  hp=4    hp=4.5
hp=0.5         1.00000 -       -       -       -       -       -       -       -       -
hp=1           1.00000 1.00000 -       -       -       -       -       -       -       -
hp=1.5         0.28054 0.28054 1.00000 -       -       -       -       -       -       -
hp=2           < 2e-16 < 2e-16 < 2e-16 < 2e-16 -       -       -       -       -       -
hp=2.5         < 2e-16 < 2e-16 < 2e-16 < 2e-16 1.00000 -       -       -       -       -
hp=3           < 2e-16 < 2e-16 < 2e-16 < 2e-16 1.00000 1.00000 -       -       -       -
hp=3.5         < 2e-16 < 2e-16 < 2e-16 < 2e-16 0.01547 0.24095 0.00088 -       -       -
hp=4           < 2e-16 < 2e-16 < 2e-16 < 2e-16 0.00020 0.00589 6.4e-06 1.00000 -       -
hp=4.5         < 2e-16 < 2e-16 < 2e-16 2.3e-15 5.5e-11 7.5e-09 4.9e-13 0.03510 0.97145 -
hp=5           5.4e-08 5.4e-08 0.01634 0.09605 < 2e-16 < 2e-16 < 2e-16 2.8e-16 4.9e-13 6.4e-06
hp=5.5         2.2e-12 2.2e-12 1.3e-05 0.00015 < 2e-16 < 2e-16 < 2e-16 2.3e-11 1.5e-08 0.00931
hp=6           1.00000 1.00000 1.00000 1.00000 < 2e-16 < 2e-16 < 2e-16 < 2e-16 < 2e-16 < 2e-16
hp=6.5         1.00000 1.00000 1.00000 1.00000 < 2e-16 < 2e-16 < 2e-16 < 2e-16 < 2e-16 < 2e-16
hp=1.5,kH=0.95 < 2e-16 < 2e-16 < 2e-16 < 2e-16 0.45144 0.03298 1.00000 6.8e-09 9.6e-12 < 2e-16
               hp=5    hp=5.5  hp=6    hp=6.5
hp=0.5         -       -       -       -
hp=1           -       -       -       -
hp=1.5         -       -       -       -
hp=2           -       -       -       -
hp=2.5         -       -       -       -
hp=3           -       -       -       -
hp=3.5         -       -       -       -
hp=4           -       -       -       -
hp=4.5         -       -       -       -
hp=5           -       -       -       -
hp=5.5         1.00000 -       -       -
hp=6           0.01634 1.3e-05 -       -
hp=6.5         0.01634 1.3e-05 1.00000 -
hp=1.5,kH=0.95 < 2e-16 < 2e-16 < 2e-16 < 2e-16

P value adjustment method: bonferroni
```

## kH = 1.15

<u>Statistics on the proportion of resilient growth as a function of hp (when $k_H$ = 1.15) with hp = 1.5, $k_H$ = 0.95 as control</u>
(1, hp = 0; 2, hp = 0.5; 3, hp = 1; 4, hp = 1.5; 5, hp = 2; 6, hp = 2.5; 7, hp = 3; 8, hp = 3.5; 9, hp = 4; 10, hp = 4.5; 11, hp = 5; 12, hp = 5.5; 13, hp = 6; 14, hp = 6.5; 15, hp=1.5, kH=0.95)

15-sample test for equality of proportions without continuity correction
data:  V_count_r out of V_count_total
X-squared = 1706.9, df = 14, p-value < 2.2e-16
alternative hypothesis: two.sided

sample estimates:

| prop 1 | prop 2 | prop 3 | prop 4 | prop 5 | prop 6 | prop 7 | prop 8 | prop 9 |
|---|---|---|---|---|---|---|---|---|
| 0.00000000 | 0.00000000 | 0.06666667 | 0.07916667 | 0.50625000 | 0.48627451 | 0.52407407 | 0.40588235 | 0.37962963 |

| prop 10 | prop 11 | prop 12 | prop 13 | prop 14 | prop 15 |
|---|---|---|---|---|---|
| 0.31111111 | 0.16666667 | 0.20740741 | 0.06666667 | 0.06666667 | 0.58148148 |

sample size:
n = 30 for each hp condition

Pairwise comparisons using Pairwise comparison of proportions

data:  V_count_r out of V_count_total

| | 1 | 2 | 3 | 4 | 5 | 6 | 7 | 8 | 9 | 10 | 11 | 12 | 13 |
|---|---|---|---|---|---|---|---|---|---|---|---|---|---|
| 2 | - | - | - | - | - | - | - | - | - | - | - | - | - |
| 3 | 3.1e-07 | 3.1e-07 | - | - | - | - | - | - | - | - | - | - | - |
| 4 | 8.5e-09 | 8.5e-09 | 1.00000 | - | - | - | - | - | - | - | - | - | - |
| 5 | < 2e-16 | < 2e-16 | < 2e-16 | < 2e-16 | - | - | - | - | - | - | - | - | - |
| 6 | < 2e-16 | < 2e-16 | < 2e-16 | < 2e-16 | 1.00000 | - | - | - | - | - | - | - | - |
| 7 | < 2e-16 | < 2e-16 | < 2e-16 | < 2e-16 | 1.00000 | 1.00000 | - | - | - | - | - | - | - |
| 8 | < 2e-16 | < 2e-16 | < 2e-16 | < 2e-16 | 0.19735 | 1.00000 | 0.01664 | - | - | - | - | - | - |
| 9 | < 2e-16 | < 2e-16 | < 2e-16 | < 2e-16 | 0.00648 | 0.06396 | 0.00026 | 1.00000 | - | - | - | - | - |
| 10 | < 2e-16 | < 2e-16 | < 2e-16 | < 2e-16 | 3.6e-08 | 1.0e-06 | 2.1e-10 | 0.17644 | 1.00000 | - | - | - | - |
| 11 | < 2e-16 | < 2e-16 | 5.3e-05 | 0.00401 | < 2e-16 | < 2e-16 | < 2e-16 | 1.5e-15 | 7.2e-13 | 4.1e-06 | - | - | - |
| 12 | < 2e-16 | < 2e-16 | 3.3e-09 | 1.4e-06 | < 2e-16 | < 2e-16 | < 2e-16 | 4.6e-10 | 8.2e-08 | 0.01393 | 1.00000 | - | - |
| 13 | 3.1e-07 | 3.1e-07 | 1.00000 | 1.00000 | < 2e-16 | < 2e-16 | < 2e-16 | < 2e-16 | < 2e-16 | < 2e-16 | 5.3e-05 | 3.3e-09 | - |
| 14 | 3.1e-07 | 3.1e-07 | 1.00000 | 1.00000 | < 2e-16 | < 2e-16 | < 2e-16 | < 2e-16 | < 2e-16 | < 2e-16 | 5.3e-05 | 3.3e-09 | 1.00000 |
| 15 | < 2e-16 | < 2e-16 | < 2e-16 | < 2e-16 | 1.00000 | 0.25469 | 1.00000 | 1.9e-06 | 5.0e-09 | < 2e-16 | < 2e-16 | < 2e-16 | < 2e-16 |

| | 14 |
|---|---|
| 2 | - |
| 3 | - |
| 4 | - |
| 5 | - |
| 6 | - |
| 7 | - |
| 8 | - |
| 9 | - |
| 10 | - |
| 11 | - |
| 12 | - |
| 13 | - |
| 14 | - |
| 15 | < 2e-16 |

P value adjustment method: bonferroni

Statistics on the average GTP cap size (# of dimers, per growth phase) as a function of hp (with all $k_H$ = 0.95)

Sample means:

```
     hp=0    hp=0.5      hp=1    hp=1.5      hp=2    hp=2.5      hp=3      hp=6    hp=6.5
 57.65886  66.81967  64.85393  72.51097  76.33327  84.08044  88.87342 125.40117 129.40041
```

sample standard deviations:

```
    hp=0   hp=0.5     hp=1   hp=1.5     hp=2   hp=2.5     hp=3     hp=6   hp=6.5
4.335404 2.711147 3.112423 6.963318 2.763598 2.813121 3.086727 1.444057 1.642250
```

sample sizes:

```
    hp=0   hp=0.5     hp=1   hp=1.5     hp=2   hp=2.5     hp=3     hp=6   hp=6.5
      54       49       51       62       48       41       41       36       36
```

Kruskal-Wallis rank sum test

```
data:  dat17$GTP by id
Kruskal-Wallis chi-squared = 393.58, df = 8, p-value < 2.2e-16
```

Pairwise comparisons using Wilcoxon rank sum test

```
data:  dat17$GTP and id

       hp=0    hp=0.5  hp=1    hp=1.5  hp=2    hp=2.5  hp=3    hp=6
hp=0.5 < 2e-16 -       -       -       -       -       -       -
hp=1   4.8e-16 0.00021 -       -       -       -       -       -
hp=1.5 < 2e-16 9.3e-16 < 2e-16 -       -       -       -       -
hp=2   < 2e-16 < 2e-16 2.4e-16 4.3e-11 -       -       -       -
hp=2.5 < 2e-16 < 2e-16 3.1e-16 4.4e-16 < 2e-16 -       -       -
hp=3   < 2e-16 < 2e-16 3.1e-16 3.1e-16 < 2e-16 < 2e-16 -       -
hp=6   1.4e-15 < 2e-16 2.9e-15 3.1e-16 < 2e-16 < 2e-16 < 2e-16 -
hp=6.5 1.4e-15 < 2e-16 2.9e-15 3.1e-16 < 2e-16 < 2e-16 < 2e-16 1.7e-14
P value adjustment method: BH
```

One-way ANOVA

```
             Df Sum Sq Mean Sq F value Pr(>F)
id            8 216156   27019    1837 <2e-16 ***
Residuals   409   6015      15
---
Signif. codes:  0 ‘***’ 0.001 ‘**’ 0.01 ‘*’ 0.05 ‘.’ 0.1 ‘ ’ 1
```

Pairwise comparisons using t tests with pooled SD

```
data:  dat17$GTP and id

       hp=0    hp=0.5  hp=1    hp=1.5  hp=2    hp=2.5  hp=3    hp=6
hp=0.5 < 2e-16 -       -       -       -       -       -       -
hp=1   < 2e-16 0.38707 -       -       -       -       -       -
hp=1.5 < 2e-16 2.4e-12 < 2e-16 -       -       -       -       -
hp=2   < 2e-16 < 2e-16 < 2e-16 1.2e-05 -       -       -       -
hp=2.5 < 2e-16 < 2e-16 < 2e-16 < 2e-16 < 2e-16 -       -       -
hp=3   < 2e-16 < 2e-16 < 2e-16 < 2e-16 < 2e-16 1.0e-06 -       -
hp=6   < 2e-16 < 2e-16 < 2e-16 < 2e-16 < 2e-16 < 2e-16 < 2e-16 -
hp=6.5 < 2e-16 < 2e-16 < 2e-16 < 2e-16 < 2e-16 < 2e-16 < 2e-16 0.00045

P value adjustment method: bonferroni
```

Statistics on time upon rapid disassembly onset(min) as a function of hp (with all $k_H$ = 0.95)

Sample means:

| hp=0 | hp=0.5 | hp=1 | hp=1.5 | hp=2 | hp=2.5 | hp=3 | hp=6 | hp=6.5 |
|---|---|---|---|---|---|---|---|---|
| 1.506268 | 1.870229 | 1.898159 | 2.199986 | 2.189203 | 2.665500 | 2.302813 | NA | NA |

sample standard deviations:

| hp=0 | hp=0.5 | hp=1 | hp=1.5 | hp=2 | hp=2.5 | hp=3 | hp=6 | hp=6.5 |
|---|---|---|---|---|---|---|---|---|
| 1.294270 | 1.373167 | 1.457356 | 1.260246 | 1.498254 | 1.466161 | 1.466352 | NA | NA |

sample sizes:

| hp=0 | hp=0.5 | hp=1 | hp=1.5 | hp=2 | hp=2.5 | hp=3 | hp=6 | hp=6.5 |
|---|---|---|---|---|---|---|---|---|
| 42 | 33 | 32 | 38 | 21 | 8 | 6 | NA | NA |

Kruskal-Wallis rank sum test

data:  dat19$time by id

Kruskal-Wallis chi-squared = 10.22, df = 6, p-value = 0.1157

Pairwise comparisons using Wilcoxon rank sum test

data:  dat19$time and id

| | hp=0 | hp=0.5 | hp=1 | hp=1.5 | hp=2 | hp=2.5 |
|---|---|---|---|---|---|---|
| hp=0.5 | 0.48 | - | - | - | - | - |
| hp=1 | 0.48 | 0.91 | - | - | - | - |
| hp=1.5 | 0.23 | 0.48 | 0.48 | - | - | - |
| hp=2 | 0.37 | 0.64 | 0.64 | 0.91 | - | - |
| hp=2.5 | 0.37 | 0.48 | 0.48 | 0.64 | 0.64 | - |
| hp=3 | 0.48 | 0.64 | 0.64 | 0.91 | 0.91 | 0.91 |

P value adjustment method: BH

Statistics on the average GTP cap size (# of dimers, per growth phase) as a function of $k_H$ (when hp = 2.0) with hp = 1.5, $k_H$ = 0.95 as control

Sample means:

| hp=1.5,kH=0.95 | hp=2,kH=0.95 | hp=2,kH=1.00 | hp=2,kH=1.05 | hp=2,kH=1.10 | hp=2,kH=1.15 |
|---|---|---|---|---|---|
| 57.65886 | 66.81967 | 72.80067 | 67.86704 | 66.63726 | 63.82437 |

| hp=2,kH=1.20 | hp=2,kH=1.25 | hp=2,kH=1.30 | hp=2,kH=1.50 | hp=2,kH=2.00 |
|---|---|---|---|---|
| 60.60838 | 57.17088 | 56.38166 | 47.14012 | 32.17991 |

sample standard deviations:

| hp=1.5,kH=0.95 | hp=2,kH=0.95 | hp=2,kH=1.00 | hp=2,kH=1.05 | hp=2,kH=1.10 | hp=2,kH=1.15 |
|---|---|---|---|---|---|
| 4.335404 | 2.711147 | 1.744213 | 10.265340 | 2.699324 | 1.800352 |

| hp=2,kH=1.20 | hp=2,kH=1.25 | hp=2,kH=1.30 | hp=2,kH=1.50 | hp=2,kH=2.00 |
|---|---|---|---|---|
| 3.974162 | 4.790612 | 4.495587 | 5.597565 | 11.155981 |

sample sizes:

| hp=1.5,kH=0.95 | hp=2,kH=0.95 | hp=2,kH=1.00 | hp=2,kH=1.05 | hp=2,kH=1.10 | hp=2,kH=1.15 |
|---|---|---|---|---|---|
| 54 | 49 | 26 | 30 | 33 | 30 |

| hp=2,kH=1.20 | hp=2,kH=1.25 | hp=2,kH=1.30 | hp=2,kH=1.50 | hp=2,kH=2.00 |
|---|---|---|---|---|
| 30 | 26 | 28 | 20 | 18 |

Kruskal-Wallis rank sum test

data: dat18$GTP by id

Kruskal-Wallis chi-squared = 286.28, df = 10, p-value < 2.2e-16

Pairwise comparisons using Wilcoxon rank sum test

data: dat18$GTP and id

| | hp=1.5,kH=0.95 | hp=2,kH=0.95 | hp=2,kH=1.00 | hp=2,kH=1.05 | hp=2,kH=1.10 | hp=2,kH=1.15 |
|---|---|---|---|---|---|---|
| hp=2,kH=0.95 | < 2e-16 | - | - | - | - | - |
| hp=2,kH=1.00 | 1.4e-12 | 6.9e-14 | - | - | - | - |
| hp=2,kH=1.05 | 6.9e-11 | 1.4e-07 | 2.6e-07 | - | - | - |
| hp=2,kH=1.10 | 8.0e-13 | 0.9700 | 3.9e-14 | 3.7e-08 | - | - |
| hp=2,kH=1.15 | 1.4e-11 | 2.2e-07 | 2.4e-15 | 1.8e-10 | 1.2e-07 | - |
| hp=2,kH=1.20 | 1.1e-05 | 1.0e-13 | 2.4e-15 | 1.6e-10 | 1.1e-11 | 2.7e-07 |
| hp=2,kH=1.25 | 0.7680 | < 2e-16 | 2.5e-14 | 4.8e-10 | 8.0e-13 | 1.4e-11 |
| hp=2,kH=1.30 | 0.0089 | < 2e-16 | 7.3e-15 | 2.7e-10 | 1.4e-13 | 1.6e-13 |
| hp=2,kH=1.50 | 3.7e-08 | 2.4e-16 | 9.3e-13 | 5.7e-10 | 4.9e-14 | 1.6e-13 |
| hp=2,kH=2.00 | 4.5e-10 | 2.3e-15 | 4.5e-12 | 2.9e-10 | 2.3e-13 | 8.0e-13 |

| | hp=2,kH=1.20 | hp=2,kH=1.25 | hp=2,kH=1.30 | hp=2,kH=1.50 |
|---|---|---|---|---|
| hp=2,kH=0.95 | - | - | - | - |
| hp=2,kH=1.00 | - | - | - | - |
| hp=2,kH=1.05 | - | - | - | - |
| hp=2,kH=1.10 | - | - | - | - |
| hp=2,kH=1.15 | - | - | - | - |
| hp=2,kH=1.20 | - | - | - | - |
| hp=2,kH=1.25 | 1.3e-05 | - | - | - |
| hp=2,kH=1.30 | 3.2e-07 | 0.0647 | - | - |
| hp=2,kH=1.50 | 5.5e-11 | 3.7e-08 | 2.5e-09 | - |
| hp=2,kH=2.00 | 1.4e-12 | 4.6e-11 | 1.2e-10 | 4.9e-07 |

P value adjustment method: BH

Statistics on time upon rapid disassembly onset(min) as a function of $k_H$ (when hp = 2.0) with hp = 1.5, $k_H$ = 0.95 as control

Sample means:

| hp=1.5,kH=0.95 | hp=2,kH=0.95 | hp=2,kH=1.00 | hp=2,kH=1.05 | hp=2,kH=1.10 | hp=2,kH=1.15 |
|---|---|---|---|---|---|
| 1.5062679 | 1.8702288 | 1.9922596 | 1.8906100 | 1.7583203 | 1.9316962 |
| hp=2,kH=1.20 | hp=2,kH=1.25 | hp=2,kH=1.30 | hp=2,kH=1.50 | hp=2,kH=2.00 | |
| 1.7007026 | 0.9234347 | 0.8455804 | 0.3550947 | 0.1021790 | |

sample standard deviations:

| hp=1.5,kH=0.95 | hp=2,kH=0.95 | hp=2,kH=1.00 | hp=2,kH=1.05 | hp=2,kH=1.10 | hp=2,kH=1.15 |
|---|---|---|---|---|---|
| 1.29427007 | 1.37316670 | 1.62133223 | 1.30465468 | 1.19464001 | 1.24972650 |
| hp=2,kH=1.20 | hp=2,kH=1.25 | hp=2,kH=1.30 | hp=2,kH=1.50 | hp=2,kH=2.00 | |
| 1.32029105 | 0.86031692 | 0.61907080 | 0.32505877 | 0.08519724 | |

sample sizes:

| hp=1.5,kH=0.95 | hp=2,kH=0.95 | hp=2,kH=1.00 | hp=2,kH=1.05 | hp=2,kH=1.10 | hp=2,kH=1.15 |
|---|---|---|---|---|---|
| 42 | 33 | 12 | 21 | 26 | 19 |
| hp=2,kH=1.20 | hp=2,kH=1.25 | hp=2,kH=1.30 | hp=2,kH=1.50 | hp=2,kH=2.00 | |
| 26 | 23 | 27 | 20 | 18 | |

Kruskal-Wallis rank sum test
data:  dat20$time by id
Kruskal-Wallis chi-squared = 85.825, df = 10, p-value = 3.588e-14

Pairwise comparisons using Wilcoxon rank sum test
data:  dat20$time and id

| | hp=1.5,kH=0.95 | hp=2,kH=0.95 | hp=2,kH=1.00 | hp=2,kH=1.05 | hp=2,kH=1.10 | hp=2,kH=1.15 |
|---|---|---|---|---|---|---|
| hp=2,kH=0.95 | 0.32259 | - | - | - | - | - |
| hp=2,kH=1.00 | 0.44484 | 0.89092 | - | - | - | - |
| hp=2,kH=1.05 | 0.28114 | 0.91004 | 0.90518 | - | - | - |
| hp=2,kH=1.10 | 0.35654 | 0.98775 | 1.00000 | 0.81338 | - | - |
| hp=2,kH=1.15 | 0.22991 | 0.81338 | 0.81338 | 0.97130 | 0.69770 | - |
| hp=2,kH=1.20 | 0.69053 | 0.81338 | 0.69770 | 0.69770 | 0.81338 | 0.64734 |
| hp=2,kH=1.25 | 0.16901 | 0.01090 | 0.06624 | 0.01090 | 0.00682 | 0.00544 |
| hp=2,kH=1.30 | 0.12630 | 0.00544 | 0.02861 | 0.00375 | 0.00426 | 0.00194 |
| hp=2,kH=1.50 | 0.00041 | 1.6e-06 | 0.00017 | 9.0e-06 | 3.7e-07 | 2.1e-06 |
| hp=2,kH=2.00 | 1.0e-07 | 1.8e-10 | 1.8e-07 | 9.6e-08 | 6.4e-10 | 9.6e-08 |

| | hp=2,kH=1.20 | hp=2,kH=1.25 | hp=2,kH=1.30 | hp=2,kH=1.50 |
|---|---|---|---|---|
| hp=2,kH=0.95 | - | - | - | - |
| hp=2,kH=1.00 | - | - | - | - |
| hp=2,kH=1.05 | - | - | - | - |
| hp=2,kH=1.10 | - | - | - | - |
| hp=2,kH=1.15 | - | - | - | - |
| hp=2,kH=1.20 | - | - | - | - |
| hp=2,kH=1.25 | 0.04110 | - | - | - |
| hp=2,kH=1.30 | 0.04110 | 0.89092 | - | - |
| hp=2,kH=1.50 | 3.0e-05 | 0.03017 | 0.00426 | - |
| hp=2,kH=2.00 | 2.4e-09 | 9.0e-06 | 2.2e-07 | 0.00251 |

P value adjustment method: BH

### Statistics on the growth phase velocity as a function of $k_H$ (when hp = 2.0) with hp = 1.5, $k_H$ = 0.95 as control

#### sample means:

| hp=1.5,kH=0.95 | hp=2,kH=0.95 | hp=2,kH=1.00 | hp=2,kH=1.05 | hp=2,kH=1.10 | hp=2,kH=1.15 | hp=2,kH=1.20 | hp=2,kH=1.25 | hp=2,kH=1.30 | hp=2,kH=1.50 | hp=2,kH=2.00 |
|---|---|---|---|---|---|---|---|---|---|---|
| 1.8478436 | 1.9923080 | 1.9567969 | 1.9099146 | 1.8119577 | 1.7745154 | 1.7178058 | 1.4617121 | 1.5247851 | 1.1242615 | 0.6626331 |

#### sample standard deviations:

| hp=1.5,kH=0.95 | hp=2,kH=0.95 | hp=2,kH=1.00 | hp=2,kH=1.05 | hp=2,kH=1.10 | hp=2,kH=1.15 | hp=2,kH=1.20 | hp=2,kH=1.25 | hp=2,kH=1.30 | hp=2,kH=1.50 | hp=2,kH=2.00 |
|---|---|---|---|---|---|---|---|---|---|---|
| 0.1793722 | 0.2757059 | 0.1770787 | 0.1425470 | 0.3359776 | 0.2861416 | 0.2824627 | 0.4960710 | 0.4314553 | 0.5609555 | 0.3932447 |

#### sample sizes:

| hp=1.5,kH=0.95 | hp=2,kH=0.95 | hp=2,kH=1.00 | hp=2,kH=1.05 | hp=2,kH=1.10 | hp=2,kH=1.15 | hp=2,kH=1.20 | hp=2,kH=1.25 | hp=2,kH=1.30 | hp=2,kH=1.50 | hp=2,kH=2.00 |
|---|---|---|---|---|---|---|---|---|---|---|
| 23 | 39 | 21 | 23 | 27 | 21 | 27 | 20 | 27 | 15 | 6 |

#### Kruskal-Wallis rank sum test

data: growth rate (μm/min)
Kruskal-Wallis chi-squared = 119.03, df = 10, p-value < 2.2e-16

#### Pairwise comparisons using Wilcoxon rank sum test

data: growth rate (μm/min)

| | hp=1.5,kH=0.95 | hp=2,kH=0.95 | hp=2,kH=1.00 | hp=2,kH=1.05 | hp=2,kH=1.10 | hp=2,kH=1.15 | hp=2,kH=1.20 | hp=2,kH=1.25 | hp=2,kH=1.30 | hp=2,kH=1.50 |
|---|---|---|---|---|---|---|---|---|---|---|
| hp=2,kH=0.95 | 2.1e-05 | - | - | - | - | - | - | - | - | - |
| hp=2,kH=1.00 | 0.01403 | 0.05704 | - | - | - | - | - | - | - | - |
| hp=2,kH=1.05 | 0.22074 | 0.00046 | 0.09895 | - | - | - | - | - | - | - |
| hp=2,kH=1.10 | 0.65514 | 2.1e-05 | 0.01853 | 0.45290 | - | - | - | - | - | - |
| hp=2,kH=1.15 | 0.16107 | 3.5e-06 | 0.00280 | 0.01853 | 0.12353 | - | - | - | - | - |
| hp=2,kH=1.20 | 0.08558 | 1.0e-06 | 0.00039 | 0.00671 | 0.04866 | 0.57090 | - | - | - | - |
| hp=2,kH=1.25 | 0.00088 | 1.7e-07 | 2.3e-05 | 3.8e-05 | 0.00099 | 0.01683 | 0.06402 | - | - | - |
| hp=2,kH=1.30 | 6.8e-05 | 7.4e-09 | 4.3e-06 | 2.4e-06 | 0.00012 | 0.00605 | 0.03482 | 0.95753 | - | - |
| hp=2,kH=1.50 | 2.4e-06 | 7.2e-08 | 3.8e-07 | 1.7e-07 | 4.6e-06 | 5.3e-06 | 0.00011 | 0.05017 | 0.00641 | - |
| hp=2,kH=2.00 | 1.5e-05 | 4.6e-06 | 2.1e-05 | 1.5e-05 | 5.4e-05 | 0.00017 | 2.1e-05 | 0.00562 | 0.00078 | 0.05607 |

P value adjustment method: BH

### Statistics on the rapid shortening phase velocity as a function of $k_H$ (when hp = 2.0) with hp = 1.5, $k_H$ = 0.95 as control

#### sample means:

| hp=1.5,kH=0.95 | hp=2,kH=0.95 | hp=2,kH=1.00 | hp=2,kH=1.05 | hp=2,kH=1.10 | hp=2,kH=1.15 | hp=2,kH=1.20 | hp=2,kH=1.25 | hp=2,kH=1.30 | hp=2,kH=1.50 | hp=2,kH=2.00 |
|---|---|---|---|---|---|---|---|---|---|---|
| -20.59369 | -20.70298 | -19.91251 | -20.55753 | -20.61000 | -20.29957 | -19.80598 | -19.99354 | -19.70679 | -19.06036 | -17.05007 |

#### sample standard deviations:

| hp=1.5,kH=0.95 | hp=2,kH=0.95 | hp=2,kH=1.00 | hp=2,kH=1.05 | hp=2,kH=1.10 | hp=2,kH=1.15 | hp=2,kH=1.20 | hp=2,kH=1.25 | hp=2,kH=1.30 | hp=2,kH=1.50 | hp=2,kH=2.00 |
|---|---|---|---|---|---|---|---|---|---|---|
| 1.4304924 | 0.3458078 | 2.6319800 | 2.0822486 | 1.9900719 | 1.7010728 | 2.5652344 | 2.7865287 | 3.2502470 | 3.2245247 | 5.0180157 |

#### sample sizes:

| hp=1.5,kH=0.95 | hp=2,kH=0.95 | hp=2,kH=1.00 | hp=2,kH=1.05 | hp=2,kH=1.10 | hp=2,kH=1.15 | hp=2,kH=1.20 | hp=2,kH=1.25 | hp=2,kH=1.30 | hp=2,kH=1.50 | hp=2,kH=2.00 |
|---|---|---|---|---|---|---|---|---|---|---|
| 21 | 18 | 10 | 18 | 25 | 17 | 23 | 23 | 27 | 20 | 18 |

#### Kruskal-Wallis rank sum test

data: rapid shortening rate (μm/min)
Kruskal-Wallis chi-squared = 12.627, df = 10, p-value = 0.2453

#### Pairwise comparisons using Wilcoxon rank sum test

data: rapid shortening rate (μm/min)

| | hp=1.5,kH=0.95 | hp=2,kH=0.95 | hp=2,kH=1.00 | hp=2,kH=1.05 | hp=2,kH=1.10 | hp=2,kH=1.15 |
|---|---|---|---|---|---|---|
| hp=2,kH=0.95 | 0.90 | - | - | - | - | - |
| hp=2,kH=1.00 | 0.90 | 0.85 | - | - | - | - |
| hp=2,kH=1.05 | 0.85 | 0.90 | 0.85 | - | - | - |
| hp=2,kH=1.10 | 0.90 | 0.65 | 0.90 | 0.84 | - | - |
| hp=2,kH=1.15 | 0.90 | 0.84 | 0.90 | 0.84 | 0.90 | - |
| hp=2,kH=1.20 | 0.86 | 0.72 | 0.90 | 0.84 | 0.98 | 0.91 |
| hp=2,kH=1.25 | 0.90 | 0.85 | 0.90 | 0.85 | 0.86 | 0.90 |
| hp=2,kH=1.30 | 0.84 | 0.59 | 0.85 | 0.59 | 0.90 | 0.85 |
| hp=2,kH=1.50 | 0.84 | 0.63 | 0.85 | 0.59 | 0.84 | 0.85 |
| hp=2,kH=2.00 | 0.40 | 0.24 | 0.59 | 0.24 | 0.57 | 0.57 |

| | hp=2,kH=1.20 | hp=2,kH=1.25 | hp=2,kH=1.30 | hp=2,kH=1.50 |
|---|---|---|---|---|
| hp=2,kH=0.95 | - | - | - | - |
| hp=2,kH=1.00 | - | - | - | - |
| hp=2,kH=1.05 | - | - | - | - |
| hp=2,kH=1.10 | - | - | - | - |
| hp=2,kH=1.15 | - | - | - | - |
| hp=2,kH=1.20 | - | - | - | - |
| hp=2,kH=1.25 | 0.87 | - | - | - |
| hp=2,kH=1.30 | 0.90 | 0.84 | - | - |
| hp=2,kH=1.50 | 0.85 | 0.84 | 0.90 | - |
| hp=2,kH=2.00 | 0.59 | 0.40 | 0.63 | 0.82 |

P value adjustment method: BH

### Statistics on the catastrophe frequency as a function of $k_H$ (when hp = 2.0) with hp = 1.5, $k_H$ = 0.95 as control

**sample means:**

| hp=1.5,kH=0.95 | hp=2,kH=0.95 | hp=2,kH=1.00 | hp=2,kH=1.05 | hp=2,kH=1.10 | hp=2,kH=1.15 | hp=2,kH=1.20 |
|---|---|---|---|---|---|---|
| 0.3159058 | 0.1865708 | 0.1614239 | 0.3264407 | 0.4431386 | 0.2948904 | 0.4987426 |

| hp=2,kH=1.25 | hp=2,kH=1.30 | hp=2,kH=1.50 | hp=2,kH=2.00 |
|---|---|---|---|
| 0.9959550 | 1.0315921 | 3.0008960 | 10.9433265 |

**sample standard deviations:**

| hp=1.5,kH=0.95 | hp=2,kH=0.95 | hp=2,kH=1.00 | hp=2,kH=1.05 | hp=2,kH=1.10 | hp=2,kH=1.15 |
|---|---|---|---|---|---|
| 0.04956530 | 0.04141973 | 0.03426123 | 0.06191876 | 0.06503169 | 0.03909026 |

| hp=2,kH=1.20 | hp=2,kH=1.25 | hp=2,kH=1.30 | hp=2,kH=1.50 | hp=2,kH=2.00 |
|---|---|---|---|---|
| 0.07591367 | 0.23083175 | 0.18072435 | 0.69630712 | 1.97280951 |

**sample sizes:**

n = 30 for all conditions

**One-way ANOVA:**

| | Df | Sum Sq | Mean Sq | F value | Pr(>F) | |
|---|---|---|---|---|---|---|
| hp_id | 10 | 3045 | 304.52 | 747.1 | <2e-16 | *** |
| Residuals | 319 | 130 | 0.41 | | | |

---

Signif. codes: 0 ‘***’ 0.001 ‘**’ 0.01 ‘*’ 0.05 ‘.’ 0.1 ‘ ’ 1

**Pairwise comparisons using t tests with pooled SD**

data: df_raw_0$F_cat and hp_id

| | hp=1.5,kH=0.95 | hp=2,kH=0.95 | hp=2,kH=1.00 | hp=2,kH=1.05 | hp=2,kH=1.10 | hp=2,kH=1.15 | hp=2,kH=1.20 |
|---|---|---|---|---|---|---|---|
| hp=2,kH=0.95 | 1.0000 | - | - | - | - | - | - |
| hp=2,kH=1.00 | 1.0000 | 1.0000 | - | - | - | - | - |
| hp=2,kH=1.05 | 1.0000 | 1.0000 | 1.0000 | - | - | - | - |
| hp=2,kH=1.10 | 1.0000 | 1.0000 | 1.0000 | 1.0000 | - | - | - |
| hp=2,kH=1.15 | 1.0000 | 1.0000 | 1.0000 | 1.0000 | 1.0000 | - | - |
| hp=2,kH=1.20 | 1.0000 | 1.0000 | 1.0000 | 1.0000 | 1.0000 | 1.0000 | - |
| hp=2,kH=1.25 | 0.0026 | 8.0e-05 | 3.9e-05 | 0.0034 | 0.0492 | 0.0015 | 0.1520 |
| hp=2,kH=1.30 | 0.0010 | 2.8e-05 | 1.3e-05 | 0.0014 | 0.0227 | 0.0006 | 0.0745 |
| hp=2,kH=1.50 | < 2e-16 | < 2e-16 | < 2e-16 | < 2e-16 | < 2e-16 | < 2e-16 | < 2e-16 |
| hp=2,kH=2.00 | < 2e-16 | < 2e-16 | < 2e-16 | < 2e-16 | < 2e-16 | < 2e-16 | < 2e-16 |

| | hp=2,kH=1.25 | hp=2,kH=1.30 | hp=2,kH=1.50 |
|---|---|---|---|
| hp=2,kH=0.95 | - | - | - |
| hp=2,kH=1.00 | - | - | - |
| hp=2,kH=1.05 | - | - | - |
| hp=2,kH=1.10 | - | - | - |
| hp=2,kH=1.15 | - | - | - |
| hp=2,kH=1.20 | - | - | - |
| hp=2,kH=1.25 | - | - | - |
| hp=2,kH=1.30 | 1.0000 | - | - |
| hp=2,kH=1.50 | < 2e-16 | < 2e-16 | - |
| hp=2,kH=2.00 | < 2e-16 | < 2e-16 | < 2e-16 |

P value adjustment method: bonferroni

Statistics on the rescue frequency as a function of $k_H$ (when hp = 2.0) with hp = 1.5, $k_H$ = 0.95 as control

**sample means:**

| hp=1.5,kH=0.95 | hp=2,kH=0.95 | hp=2,kH=1.00 | hp=2,kH=1.05 | hp=2,kH=1.10 | hp=2,kH=1.15 | hp=2,kH=1.20 |
|---|---|---|---|---|---|---|
| 2.8614859 | 2.8187026 | 4.4079023 | 4.2181390 | 2.4215961 | 3.2819451 | 1.7211113 |

| hp=2,kH=1.25 | hp=2,kH=1.30 | hp=2,kH=1.50 | hp=2,kH=2.00 |
|---|---|---|---|
| 1.4486349 | 1.9824224 | 0.4050684 | 0.0000000 |

**sample standard deviations:**

| hp=1.5,kH=0.95 | hp=2,kH=0.95 | hp=2,kH=1.00 | hp=2,kH=1.05 | hp=2,kH=1.10 | hp=2,kH=1.15 |
|---|---|---|---|---|---|
| 0.9113240 | 0.9741829 | 0.7158820 | 1.3591723 | 0.6555230 | 0.9679373 |

| hp=2,kH=1.20 | hp=2,kH=1.25 | hp=2,kH=1.30 | hp=2,kH=1.50 | hp=2,kH=2.00 |
|---|---|---|---|---|
| 0.6061353 | 0.4681297 | 0.6288714 | 0.3032989 | 0.0000000 |

**sample sizes:**

n = 30 for all conditions

**One-way ANOVA:**

| | Df | Sum Sq | Mean Sq | F value | Pr(>F) | |
|---|---|---|---|---|---|---|
| hp_id | 10 | 591.6 | 59.16 | 98.9 | <2e-16 | *** |
| Residuals | 319 | 190.8 | 0.60 | | | |

---
Signif. codes: 0 ‘***’ 0.001 ‘**’ 0.01 ‘*’ 0.05 ‘.’ 0.1 ‘ ’ 1

**Pairwise comparisons using t tests with pooled SD**

data: df_raw_0$F_res and hp_id

| | hp=1.5,kH=0.95 | hp=2,kH=0.95 | hp=2,kH=1.00 | hp=2,kH=1.05 | hp=2,kH=1.10 | hp=2,kH=1.15 | hp=2,kH=1.20 |
|---|---|---|---|---|---|---|---|
| hp=2,kH=0.95 | 1.00000 | - | - | - | - | - | - |
| hp=2,kH=1.00 | 7.1e-12 | 1.7e-12 | - | - | - | - | - |
| hp=2,kH=1.05 | 3.0e-09 | 8.0e-10 | 1.00000 | - | - | - | - |
| hp=2,kH=1.10 | 1.00000 | 1.00000 | < 2e-16 | 1.2e-15 | - | - | - |
| hp=2,kH=1.15 | 1.00000 | 1.00000 | 2.1e-06 | 0.00023 | 0.00121 | - | - |
| hp=2,kH=1.20 | 1.4e-06 | 4.4e-06 | < 2e-16 | < 2e-16 | 0.02842 | 4.4e-12 | - |
| hp=2,kH=1.25 | 5.2e-10 | 2.0e-09 | < 2e-16 | < 2e-16 | 9.6e-05 | 3.1e-16 | 1.00000 |
| hp=2,kH=1.30 | 0.00081 | 0.00201 | < 2e-16 | < 2e-16 | 1.00000 | 1.6e-08 | 1.00000 |
| hp=2,kH=1.50 | < 2e-16 | < 2e-16 | < 2e-16 | < 2e-16 | < 2e-16 | < 2e-16 | 1.0e-08 |
| hp=2,kH=2.00 | < 2e-16 | < 2e-16 | < 2e-16 | < 2e-16 | < 2e-16 | < 2e-16 | 1.8e-14 |

| | hp=2,kH=1.25 | hp=2,kH=1.30 | hp=2,kH=1.50 |
|---|---|---|---|
| hp=2,kH=0.95 | - | - | - |
| hp=2,kH=1.00 | - | - | - |
| hp=2,kH=1.05 | - | - | - |
| hp=2,kH=1.10 | - | - | - |
| hp=2,kH=1.15 | - | - | - |
| hp=2,kH=1.20 | - | - | - |
| hp=2,kH=1.25 | - | - | - |
| hp=2,kH=1.30 | 0.43475 | - | - |
| hp=2,kH=1.50 | 1.7e-05 | 2.5e-12 | - |
| hp=2,kH=2.00 | 1.7e-10 | < 2e-16 | 1.00000 |

P value adjustment method: bonferroni

Statistics on the growth phase velocity as a function of hp (when $k_H$ = 1.15) with hp = 1.5, $k_H$ = 0.95 as control

sample means:

| hp=0 | hp=0.5 | hp=1 | hp=1.5 | hp=2 | hp=2.5 | hp=3 |
|---|---|---|---|---|---|---|
| 1.070330 | 1.365858 | 1.188869 | 1.645388 | 1.774515 | 2.092361 | 2.338934 |
| hp=3.5 | hp=4 | hp=4.5 | hp=5 | hp=5.5 | hp=6 | hp=6.5 |
| 2.318940 | 2.716178 | 2.949662 | 3.205003 | 3.340379 | 3.638772 | 3.784919 |
| hp=1.5,kH=0.95 | | | | | | |
| 1.847844 | | | | | | |

sample standard deviations:

| hp=0 | hp=0.5 | hp=1 | hp=1.5 | hp=2 | hp=2.5 |
|---|---|---|---|---|---|
| 0.37330312 | 0.35610312 | 0.35144047 | 0.23195548 | 0.28614157 | 0.20410773 |
| hp=3 | hp=3.5 | hp=4 | hp=4.5 | hp=5 | hp=5.5 |
| 0.09186812 | 0.59131107 | 0.33500434 | 0.19055272 | 0.16878842 | 0.14388677 |
| hp=6 | hp=6.5 | hp=1.5,kH=0.95 | | | |
| 0.07757024 | 0.19000556 | 0.17937217 | | | |

sample sizes:

| hp=0 | hp=0.5 | hp=1 | hp=1.5 | hp=2 | hp=2.5 |
|---|---|---|---|---|---|
| 22 | 17 | 22 | 24 | 21 | 24 |
| hp=3 | hp=3.5 | hp=4 | hp=4.5 | hp=5 | hp=5.5 |
| 21 | 21 | 18 | 20 | 18 | 18 |
| hp=6 | hp=6.5 | hp=1.5,kH=0.95 | | | |
| 18 | 18 | 23 | | | |

Kruskal-Wallis rank sum test

data: dat1$V_g by id

Kruskal-Wallis chi-squared = 277.12, df = 14, p-value < 2.2e-16

Pairwise comparisons using Wilcoxon rank sum test

data: dat1$V_g and id

| | hp=0 | hp=0.5 | hp=1 | hp=1.5 | hp=2 | hp=2.5 | hp=3 | hp=3.5 | hp=4 | hp=4.5 | hp=5 |
|---|---|---|---|---|---|---|---|---|---|---|---|
| hp=0.5 | 0.00202 | - | - | - | - | - | - | - | - | - | - |
| hp=1 | 0.28866 | 0.11234 | - | - | - | - | - | - | - | - | - |
| hp=1.5 | 1.4e-08 | 0.00177 | 1.7e-06 | - | - | - | - | - | - | - | - |
| hp=2 | 6.6e-09 | 1.5e-06 | 1.4e-08 | 0.01126 | - | - | - | - | - | - | - |
| hp=2.5 | 2.4e-11 | 8.1e-09 | 8.1e-11 | 1.2e-08 | 5.6e-07 | - | - | - | - | - | - |
| hp=3 | 2.7e-11 | 1.5e-10 | 2.7e-11 | 2.4e-11 | 3.1e-11 | 1.5e-07 | - | - | - | - | - |
| hp=3.5 | 2.7e-08 | 8.9e-06 | 3.5e-07 | 3.6e-05 | 0.00024 | 0.00021 | 0.00118 | - | - | - | - |
| hp=4 | 8.1e-11 | 8.6e-08 | 2.1e-09 | 2.4e-08 | 1.2e-07 | 5.9e-08 | 1.5e-07 | 5.1e-06 | - | - | - |
| hp=4.5 | 3.1e-11 | 2.7e-10 | 3.1e-11 | 2.4e-11 | 3.9e-11 | 2.4e-11 | 4.7e-09 | 4.9e-08 | 0.00019 | - | - |
| hp=5 | 5.8e-11 | 8.0e-10 | 5.8e-11 | 3.1e-11 | 7.7e-11 | 3.1e-11 | 7.7e-11 | 7.7e-11 | 5.7e-07 | 9.2e-06 | - |
| hp=5.5 | 5.8e-11 | 8.0e-10 | 5.8e-11 | 3.1e-11 | 7.7e-11 | 3.1e-11 | 7.7e-11 | 7.7e-11 | 4.4e-10 | 4.5e-08 | 0.00538 |
| hp=6 | 5.8e-11 | 8.0e-10 | 5.8e-11 | 3.1e-11 | 7.7e-11 | 3.1e-11 | 7.7e-11 | 7.7e-11 | 4.4e-10 | 1.3e-10 | 8.0e-10 |
| hp=6.5 | 5.8e-11 | 8.0e-10 | 5.8e-11 | 3.1e-11 | 7.7e-11 | 3.1e-11 | 7.7e-11 | 7.7e-11 | 4.4e-10 | 3.1e-09 | 2.7e-07 |
| hp=1.5,kH=0.95 | 2.8e-10 | 7.7e-07 | 2.3e-09 | 0.00014 | 0.14783 | 4.2e-07 | 2.4e-11 | 0.00014 | 5.0e-08 | 2.7e-11 | 4.3e-11 |

| | hp=5.5 | hp=6 | hp=6.5 |
|---|---|---|---|
| hp=0.5 | - | - | - |
| hp=1 | - | - | - |
| hp=1.5 | - | - | - |
| hp=2 | - | - | - |
| hp=2.5 | - | - | - |
| hp=3 | - | - | - |
| hp=3.5 | - | - | - |
| hp=4 | - | - | - |
| hp=4.5 | - | - | - |
| hp=5 | - | - | - |
| hp=5.5 | - | - | - |
| hp=6 | 6.1e-08 | - | - |
| hp=6.5 | 3.5e-07 | 9.4e-07 | - |
| hp=1.5,kH=0.95 | 4.3e-11 | 4.3e-11 | 4.3e-11 |

P value adjustment method: BH

### Statistics on the rapid shortening phase velocity as a function of hp (when $k_H$ = 1.15) with hp = 1.5, $k_H$ = 0.95 as control

**sample means:**

| hp=0 | hp=0.5 | hp=1 | hp=1.5 | hp=2 | hp=2.5 | hp=3 |
|---|---|---|---|---|---|---|
| -18.11516 | -17.98693 | -17.83075 | -19.72450 | -20.29957 | -22.13365 | -22.74434 |

| hp=3.5 | hp=4 | hp=4.5 | hp=5 | hp=5.5 | hp=1.5,kH=0.95 |
|---|---|---|---|---|---|
| -24.64659 | -23.69735 | -22.35211 | -24.20552 | -27.69085 | -20.59369 |

**sample standard deviations:**

| hp=0 | hp=0.5 | hp=1 | hp=1.5 | hp=2 | hp=2.5 |
|---|---|---|---|---|---|
| 1.769088 | 3.662756 | 2.919131 | 2.099426 | 1.701073 | 3.021934 |

| hp=3 | hp=3.5 | hp=4 | hp=4.5 | hp=5 | hp=5.5 |
|---|---|---|---|---|---|
| 2.457590 | 1.889195 | 2.092914 | 5.392399 | 1.026006 | 2.041285 |

| hp=1.5,kH=0.95 |
|---|
| 1.430492 |

**sample sizes:**

| hp=0 | hp=0.5 | hp=1 | hp=1.5 | hp=2 | hp=2.5 |
|---|---|---|---|---|---|
| 23 | 21 | 25 | 23 | 17 | 21 |

| hp=3 | hp=3.5 | hp=4 | hp=4.5 | hp=5 | hp=5.5 |
|---|---|---|---|---|---|
| 17 | 14 | 13 | 9 | 3 | 5 |

| hp=1.5,kH=0.95 |
|---|
| 21 |

**Kruskal-Wallis rank sum test**

data: dat2p$V_rs by id

Kruskal-Wallis chi-squared = 136.4, df = 12, p-value < 2.2e-16

**Pairwise comparisons using Wilcoxon rank sum test**

data: dat2p$V_rs and id

| | hp=0 | hp=0.5 | hp=1 | hp=1.5 | hp=2 | hp=2.5 | hp=3 | hp=3.5 | hp=4 | hp=4.5 | hp=5 | hp=5.5 |
|---|---|---|---|---|---|---|---|---|---|---|---|---|
| hp=0.5 | 0.01877 | - | - | - | - | - | - | - | - | - | - | - |
| hp=1 | 0.21446 | 0.26105 | - | - | - | - | - | - | - | - | - | - |
| hp=1.5 | 8.6e-06 | 0.01180 | 0.00012 | - | - | - | - | - | - | - | - | - |
| hp=2 | 1.8e-06 | 0.00090 | 7.9e-06 | 0.26175 | - | - | - | - | - | - | - | - |
| hp=2.5 | 1.8e-06 | 3.5e-06 | 1.2e-06 | 2.2e-06 | 9.0e-05 | - | - | - | - | - | - | - |
| hp=3 | 8.2e-07 | 1.6e-06 | 3.3e-07 | 8.8e-07 | 2.5e-05 | 0.94275 | - | - | - | - | - | - |
| hp=3.5 | 8.5e-09 | 1.7e-08 | 8.5e-09 | 8.5e-09 | 3.9e-07 | 0.00356 | 0.01210 | - | - | - | - | - |
| hp=4 | 8.5e-06 | 8.0e-06 | 2.8e-06 | 9.9e-06 | 5.7e-05 | 0.02931 | 0.06700 | 0.64009 | - | - | - | - |
| hp=4.5 | 0.01469 | 0.01093 | 0.01225 | 0.01651 | 0.03248 | 0.18002 | 0.31999 | 0.74561 | 0.87837 | - | - | - |
| hp=5 | 0.00146 | 0.00175 | 0.00122 | 0.00146 | 0.00304 | 0.10154 | 0.25122 | 0.88142 | 0.75176 | 1.00000 | - | - |
| hp=5.5 | 5.7e-05 | 7.7e-05 | 4.4e-05 | 5.7e-05 | 0.00016 | 0.00026 | 0.00165 | 0.01954 | 0.00465 | 0.01700 | 0.04567 | - |
| hp=1.5,kH=0.95 | 1.2e-06 | 0.00151 | 8.3e-06 | 0.15550 | 0.76704 | 0.00011 | 6.0e-05 | 8.2e-07 | 0.00010 | 0.03247 | 0.00642 | 7.7e-05 |

P value adjustment method: BH

(hp = 6 and hp = 6.5 were not included due to limited sample size in each case).

Statistics on the catastrophe frequency as a function of hp (when $k_H$ = 1.15) with hp = 1.5, $k_H$ = 0.95 as control

sample means:

| hp=0 | hp=0.5 | hp=1 | hp=1.5 | hp=2 | hp=2.5 | hp=3 |
|---|---|---|---|---|---|---|
| 1.23817939 | 1.53128795 | 1.04309285 | 0.66387077 | 0.29489044 | 0.33399983 | 0.23335031 |

| hp=3.5 | hp=4 | hp=4.5 | hp=5 | hp=5.5 | hp=6 | hp=6.5 |
|---|---|---|---|---|---|---|
| 0.21040483 | 0.18276485 | 0.12985627 | 0.04574410 | 0.07994544 | 0.02085179 | 0.01644581 |

| hp=1.5,kH=0.95 |
|---|
| 0.31590581 |

sample standard deviations:

| hp=0 | hp=0.5 | hp=1 | hp=1.5 | hp=2 | hp=2.5 |
|---|---|---|---|---|---|
| 0.25998775 | 0.36885877 | 0.20502140 | 0.10061850 | 0.03909026 | 0.05166671 |

| hp=3 | hp=3.5 | hp=4 | hp=4.5 | hp=5 | hp=5.5 |
|---|---|---|---|---|---|
| 0.04959144 | 0.04892537 | 0.04900185 | 0.04154192 | 0.02397111 | 0.03128181 |

| hp=6 | hp=6.5 | hp=1.5,kH=0.95 |
|---|---|---|
| 0.01527552 | 0.01377869 | 0.04956530 |

sample sizes:
n = 30 for all conditions

One-way ANOVA:

| | Df | Sum Sq | Mean Sq | F value | Pr(>F) | |
|---|---|---|---|---|---|---|
| hp_id | 14 | 95.49 | 6.821 | 374.2 | <2e-16 | *** |
| Residuals | 435 | 7.93 | 0.018 | | | |

---
Signif. codes:  0 ‘***’ 0.001 ‘**’ 0.01 ‘*’ 0.05 ‘.’ 0.1 ‘ ’ 1

Pairwise comparisons using t tests with pooled SD

data:  df_raw_0$F_cat and hp_id

| | hp=0 | hp=0.5 | hp=1 | hp=1.5 | hp=2 | hp=2.5 | hp=3 | hp=3.5 | hp=4 | hp=4.5 | hp=5 |
|---|---|---|---|---|---|---|---|---|---|---|---|
| hp=0.5 | 6.3e-14 | - | - | - | - | - | - | - | - | - | - |
| hp=1 | 4.1e-06 | < 2e-16 | - | - | - | - | - | - | - | - | - |
| hp=1.5 | < 2e-16 | < 2e-16 | < 2e-16 | - | - | - | - | - | - | - | - |
| hp=2 | < 2e-16 | < 2e-16 | < 2e-16 | < 2e-16 | - | - | - | - | - | - | - |
| hp=2.5 | < 2e-16 | < 2e-16 | < 2e-16 | < 2e-16 | 1.00000 | - | - | - | - | - | - |
| hp=3 | < 2e-16 | < 2e-16 | < 2e-16 | < 2e-16 | 1.00000 | 0.42836 | - | - | - | - | - |
| hp=3.5 | < 2e-16 | < 2e-16 | < 2e-16 | < 2e-16 | 1.00000 | 0.04561 | 1.00000 | - | - | - | - |
| hp=4 | < 2e-16 | < 2e-16 | < 2e-16 | < 2e-16 | 0.14643 | 0.00188 | 1.00000 | 1.00000 | - | - | - |
| hp=4.5 | < 2e-16 | < 2e-16 | < 2e-16 | < 2e-16 | 0.00031 | 9.8e-07 | 0.33118 | 1.00000 | 1.00000 | - | - |
| hp=5 | < 2e-16 | < 2e-16 | < 2e-16 | < 2e-16 | 4.0e-10 | 1.7e-13 | 1.3e-05 | 0.00033 | 0.01035 | 1.00000 | - |
| hp=5.5 | < 2e-16 | < 2e-16 | < 2e-16 | < 2e-16 | 1.7e-07 | 1.6e-10 | 0.00143 | 0.02170 | 0.35220 | 1.00000 | 1.00000 |
| hp=6 | < 2e-16 | < 2e-16 | < 2e-16 | < 2e-16 | 3.2e-12 | 8.5e-16 | 2.5e-07 | 9.5e-06 | 0.00047 | 0.19790 | 1.00000 |
| hp=6.5 | < 2e-16 | < 2e-16 | < 2e-16 | < 2e-16 | 1.3e-12 | 3.2e-16 | 1.2e-07 | 4.8e-06 | 0.00026 | 0.12909 | 1.00000 |
| hp=1.5,kH=0.95 | < 2e-16 | < 2e-16 | < 2e-16 | < 2e-16 | 1.00000 | 1.00000 | 1.00000 | 0.27524 | 0.01609 | 1.6e-05 | 6.9e-12 |

| | hp=5.5 | hp=6 | hp=6.5 |
|---|---|---|---|
| hp=0.5 | - | - | - |
| hp=1 | - | - | - |
| hp=1.5 | - | - | - |
| hp=2 | - | - | - |
| hp=2.5 | - | - | - |
| hp=3 | - | - | - |
| hp=3.5 | - | - | - |
| hp=4 | - | - | - |
| hp=4.5 | - | - | - |
| hp=5 | - | - | - |
| hp=5.5 | - | - | - |
| hp=6 | 1.00000 | - | - |
| hp=6.5 | 1.00000 | 1.00000 | - |
| hp=1.5,kH=0.95 | 4.4e-09 | 4.2e-14 | 1.6e-14 |

P value adjustment method: bonferroni

Statistics on the rescue frequency as a function of hp (when $k_H$ = 1.15) with hp = 1.5, $k_H$ = 0.95 as control

**sample means:**

| hp=0 | hp=0.5 | hp=1 | hp=1.5 | hp=2 | hp=2.5 | hp=3 | hp=3.5 | hp=4 | hp=4.5 | hp=5 | hp=5.5 | hp=1.5,kH=0.95 |
|---|---|---|---|---|---|---|---|---|---|---|---|---|
| 0.8643374 | 0.5560779 | 1.3219360 | 1.5996068 | 3.2819451 | 3.9462351 | 3.3653840 | 6.7664929 | 3.8043001 | 6.3634276 | 7.2552282 | 10.2151847 | 2.8614859 |

**sample standard deviations:**

| hp=0 | hp=0.5 | hp=1 | hp=1.5 | hp=2 | hp=2.5 | hp=3 | hp=3.5 | hp=4 | hp=4.5 | hp=5 | hp=5.5 | hp=1.5,kH=0.95 |
|---|---|---|---|---|---|---|---|---|---|---|---|---|
| 0.2740079 | 0.3295510 | 0.2797284 | 0.4133295 | 0.9679373 | 0.8702800 | 0.9941021 | 1.7687742 | 1.1934573 | 3.0340968 | 4.8565439 | 16.3593290 | 0.9113240 |

**sample sizes:**

n = 30 for all conditions

**One-way ANOVA:**

| | Df | Sum Sq | Mean Sq | F value | Pr(>F) | |
|---|---|---|---|---|---|---|
| hp_id | 14 | 9732 | 695.1 | 32.52 | <2e-16 | *** |
| Residuals | 418 | 8935 | 21.4 | | | |

---

Signif. codes:  0 ‘***’ 0.001 ‘**’ 0.01 ‘*’ 0.05 ‘.’ 0.1 ‘ ’ 1

17 observations deleted due to missingness

**Pairwise comparisons using t tests with pooled SD**

data:  df_raw_0$F_res and hp_id

| | hp=0 | hp=0.5 | hp=1 | hp=1.5 | hp=2 | hp=2.5 | hp=3 | hp=3.5 | hp=4 | hp=4.5 | hp=5 | hp=5.5 |
|---|---|---|---|---|---|---|---|---|---|---|---|---|
| hp=0.5 | 1.00000 | - | - | - | - | - | - | - | - | - | - | - |
| hp=1 | 1.00000 | 1.00000 | - | - | - | - | - | - | - | - | - | - |
| hp=1.5 | 1.00000 | 1.00000 | 1.00000 | - | - | - | - | - | - | - | - | - |
| hp=2 | 1.00000 | 1.00000 | 1.00000 | 1.00000 | - | - | - | - | - | - | - | - |
| hp=2.5 | 1.00000 | 0.49696 | 1.00000 | 1.00000 | 1.00000 | - | - | - | - | - | - | - |
| hp=3 | 1.00000 | 1.00000 | 1.00000 | 1.00000 | 1.00000 | 1.00000 | - | - | - | - | - | - |
| hp=3.5 | 0.00012 | 3.2e-05 | 0.00070 | 0.00198 | 0.38866 | 1.00000 | 0.48312 | - | - | - | - | - |
| hp=4 | 1.00000 | 0.71189 | 1.00000 | 1.00000 | 1.00000 | 1.00000 | 1.00000 | 1.00000 | - | - | - | - |
| hp=4.5 | 0.00057 | 0.00017 | 0.00311 | 0.00817 | 1.00000 | 1.00000 | 1.00000 | 1.00000 | 1.00000 | - | - | - |
| hp=5 | 1.9e-05 | 5.0e-06 | 0.00013 | 0.00038 | 0.11019 | 0.65618 | 0.13988 | 1.00000 | 0.45821 | 1.00000 | - | - |
| hp=5.5 | 4.2e-12 | 6.8e-13 | 5.7e-11 | 2.6e-10 | 1.3e-06 | 2.5e-05 | 1.9e-06 | 0.42697 | 1.4e-05 | 0.14189 | 1.00000 | - |
| hp=1.5,kH=0.95 | 1.00000 | 1.00000 | 1.00000 | 1.00000 | 1.00000 | 1.00000 | 1.00000 | 0.12177 | 1.00000 | 0.37123 | 0.03111 | 1.8e-07 |

P value adjustment method: Bonferroni

(hp = 6 and hp = 6.5 were not included due to limited sample size in each case).
